# Mitotree: The Universal Human Mitochondrial Reference Phylogeny at 10× the Resolution

**DOI:** 10.64898/2026.05.28.728540

**Authors:** Paul A. Maier, Göran Runfeldt, Roberta J. Estes, Pablo A. Goloboff, John Detsikas, AJ M. Burke, Michael T. Sager, Miguel G. Vilar

## Abstract

Mitochondrial DNA (mtDNA) is the oldest and most prevalent source of human molecular phylogenetic reconstruction. Everyone alive today traces their matrilines to one female African ancestor who lived some 145 thousand years ago. For decades, PhyloTree provided researchers with a moderately sized reference tree (v17: n=24,275 sequences, n=5,438 branches). However, hundreds of thousands of sequences are available today, and since PhyloTree’s retirement in 2016, there is a need for an actively maintained phylogenetic reference system. In addition, no currently published method can wield such a large de novo reconstruction while dealing with the rampant homoplasy seen in mtDNA and adhering to the accuracy standards expected in this well-studied field. In this paper we introduce Mitotree, the largest human phylogeny ever described (n≈330,000 complete sequences, n≈54,000 branches), and a new recursive phylogenetic pipeline for estimating it. We incorporate ancient DNA, relaxed clock age estimates (TMRCAs), and public databases (e.g., GenBank, 1000 Genomes, HGDP, and SGDP). We report approximately 180 previously unknown branches older than 30,000 years. The median TMRCA of terminal haplogroups is 2,000 years more recent than PhyloTree. Our pipeline offers a novel divide-and-conquer approach that tackles huge heuristic searches within tractable runtimes, provides high accuracy, and reasonable confidence estimates. Our validation found a false negative rate of 3.5%, and a false positive rate of 1.0%. Among the most striking findings are an ∼83 kya split of African L2e (the oldest in our dataset), the resolution of Ötzi’s K1f into a clade with living descendants, new ethnic founder haplogroups (e.g., Jewish diaspora), and placement of historical figures such as Abraham Lincoln. To our knowledge, Mitotree is the largest de novo haplotype phylogeny built for humanity, and is a continually improved resource for academic, genealogical, and medical research.

## Introduction

Mitochondrial DNA (mtDNA) was the first genetic marker to give humanity a glimpse into our own prehistorical origins, in the 1980s (Anderson et al., 1981). However, only a tiny fraction of our phylogenetic past was sampled in those early studies (e.g., Cann et al., 1987; Vigilant et al., 1991; Chen et al., 1995). Expanding such phylogenetic knowledge of human haplotypes remains an important goal today. Unlike most nuclear DNA (excluding Y-DNA), which recombines and hence obscures homology necessary to trace ancestor-descendant relationships more than a few generations old (Arenas and Posada, 2010; Henn et al., 2012; Rannala, 2025), mtDNA is inherited intact from mother to child. This special property of mtDNA (its lack of attenuation) allows us to reconstruct shared ancestral relationships for all matrilines of humankind. Compared to Y-DNA, mtDNA sequences are much more ubiquitous and cost-effective to produce, which in principle allows us to build a tree with massive representation from modern and ancient humans, and archaic hominins. However, practical limitations in both data availability, and phylogenetic methods, have precluded the vast majority of human mtDNA from being placed in our single shared tree.

Between 2008 and 2016, the principal reference tree for human mtDNA was PhyloTree (van Oven and Kayser, 2009; van Oven, 2015), which was subsequently discontinued (Fig. **1**). This final version of PhyloTree (v17; hereafter “PTv17”) included 24,275 full and partial mitogenome sequences, which resolved 5,438 branches or “haplogroups.” Its sampling relied on GenBank (Sayers et al., 2025) sequences from that era, and other public repositories such as the 1000 Genomes Project (Siva, 2008), while its tree topology was manually constructed via data curation and grafting onto previous trees. As previously noted, PTv17 offered a wealth of expert knowledge and annotation, but manual grafting onto previous trees can introduce bias, and prevent automated scaling to much larger phylogenies (Blanco et al., 2011). Other databases such as MITOMAP (Kogelnik, 1996), mtDB (Ingman, 2006), HmtDB (Attimonelli et al., 2005), AmtDB (Ehler et al., 2019), and mitoLEAF (Huber et al., 2025) were concurrently or subsequently founded, yet have not overcome the gap in our ability to reconstruct trees from scratch using the 100,000s of currently available full sequences (e.g., >40,000 GenBank samples added since PTv17, and >270,000 private FamilyTreeDNA samples).

**Figure 1:**
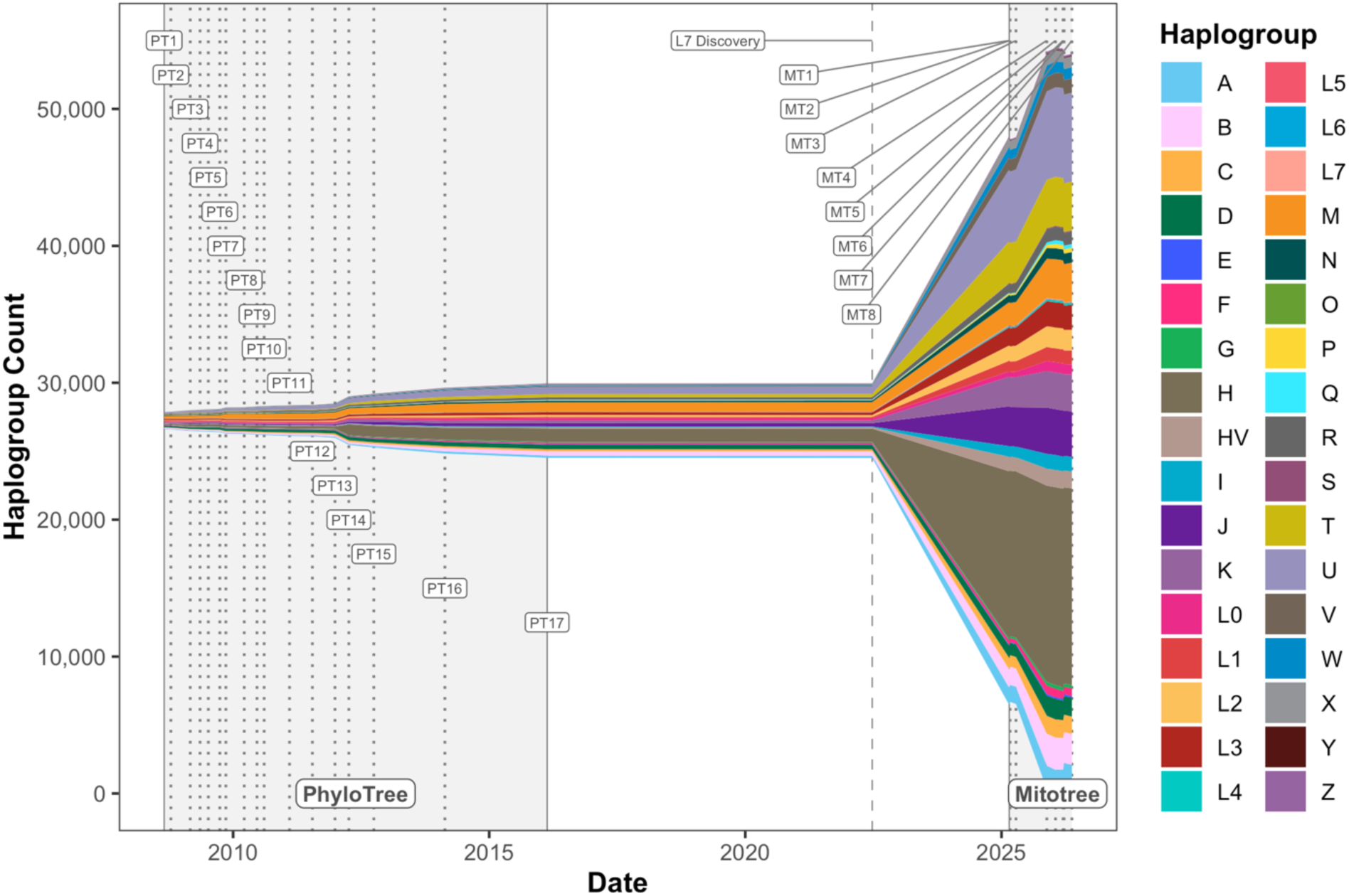
Mitochondrial Tree Growth 2008–2026: Total number of haplogroups (and split by root) over time, beginning with PhyloTree v1 (“PT1”), and ending with the current version of Mitotree (“MT8”). The L7 discovery (Maier et al., 2022) marks the beginning of Mitotree’s construction.

A genuine gap exists in large-scale phylogenetic methodology to reconstruct a modern version of PTv17 while maintaining reasonable accuracy and speed. Three categories of phylogenetic methods have attempted to address this gap: (1) scaled maximum likelihood (ML), (2) divide-and-conquer, and (3) sparse pandemic methods. ML estimation is popular due to its explicit models of evolution and statistical consistency, yet finding the optimal tree is NP-hard (Roch, 2006), hence exceedingly slow or impossible for many 1,000s of taxa. Heuristic searches have been vastly improved via better implementation and parallelization, but scaled ML tools still struggle: IQ-TREE2 (Minh et al., 2020) and RAXML-NG (Kozlov et al., 2019) either fail or give nearly 100% false negative error with 50,000 taxa (Zaharias and Warnow, 2022). Although FASTTREE2 (Price et al., 2010) and VERYFASTTREE4 (Piñeiro and Pichel, 2024) can complete runs with 1,000,000 taxa, benchmarks show much lower accuracy (Sayyari et al., 2017; Lees et al., 2018; Smirnov and Warnow, 2021). Divide-and-conquer approaches such as NJMERGE (Molloy and Warnow, 2019a), TREEMERGE (Molloy and Warnow, 2019b), and GTM (Smirnov and Warnow, 2020) develop a promising strategy: compute a “rough” starting tree, partition it and perform high-quality ML analyses on each subset, then merge the parts together. However, only GTM has been successfully benchmarked with 50,000 taxa, and although it approaches RAXML-NG accuracy, it suffers slower runtimes than FASTTREE2 (Park et al., 2021). No divide-and-conquer approaches have been scaled to 100,000s of sequences for runtime or accuracy, considered multiple replicates, or addressed the problematic attributes of mtDNA such as extreme homoplasy and heterotachy. Pandemic methods such as MAPLE (De Maio et al., 2023), USHER (Turakhia et al., 2021), and MATOPTIMIZE (Ye et al., 2022) successfully handle 1,000,000s of taxa up to thousands of times faster than other ML methods. However, they exploit unique properties of SARS-CoV-2 genomes: recent divergence with sparse mutations (<0.1% pairwise divergence), no rate heterogeneity, no heterotachy, and no saturated sites from homoplasy, all violated in human mtDNA evolution (Endicott and Ho, 2008).

To overcome these problems, we developed a novel pipeline to build Mitotree, the largest human phylogeny to date (Fig. **2**). Our criteria included: (1) accuracy, (2) efficiency on huge datasets, (3) consistency between frequent re-estimations with new data, (4) handling of homoplasy, rate heterogeneity, and heterotachy, and (5) continuity of branch naming and variant annotations with PTv17. Weighted maximum parsimony (WMP) was chosen in a divide-and-conquer framework to balance the clear tradeoff between speed and accuracy. Data-driven weighting of characters based on observed levels of homoplasy (i.e., implied weighting;

**Figure 2:**
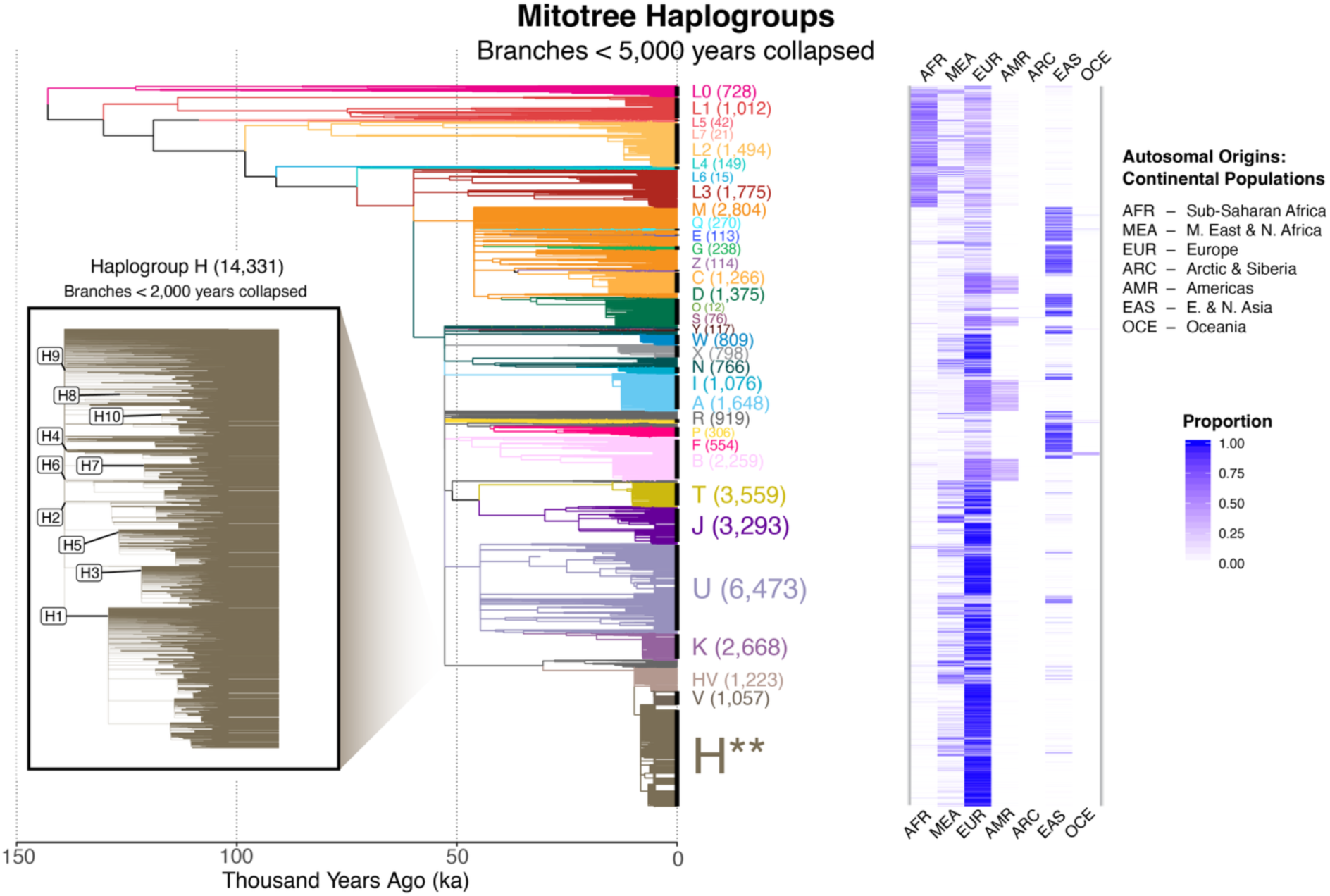
Mitotree Haplogroup Counts and Population Origins. Mitotree (53,588 branches) summarized by collapsing younger branches (<5 kya). Haplogroup H (comprising more than 25% of the tree) is shown as an inset with branches <2 kya collapsed. Autosomal population origins are summarized at the continental level where available from myOrigins 3.0 (Maier et al., 2021) for comparison.

Goloboff, 1993b) can actually outperform the ML and Bayesian methods of binning high gamma-rate mutations (+Γ), and with clear speed advantages (Goloboff et al., 2018; Torres et al., 2022; Morales and Goloboff, 2023; Wright and Wynd, 2024). We aim to release regular updates by re-estimating Mitotree from scratch, as placement-only methods can fail to detect global inaccuracies (e.g., Haplogroup L7 went undetected in PTv17 despite being 100,000 years old; Maier et al., 2022). Mitotree provides unprecedented resolution (10× PhyloTree), over 180 new haplogroups older than 30,000 years, and is a freely available resource for all academic, medical, and other non-commercial use.

## Materials and methods

### Data sources and filtering

We used 331,221 full mitochondrial sequences (n=177,196 unique haplotypes) primarily from seven sources (Table **S1**): FamilyTreeDNA (FTDNA) mtFull database (n=272,787), NCBI GenBank (n=49,625; Sayers et al., 2025), 1000 Genomes (n=3,484; Siva, 2008), HGDP (n=776; Bergström et al., 2020), GGVP (n=392; Band et al., 2019), SGDP (n=243; Mallick et al., 2016), CNCB (n=179), and miscellaneous academic studies (n=3,735), which comprised most of the 3,206 ancient DNA (aDNA) samples. We defined “full” sequences to contain 16,450–16,700 nt with ≤ 10 (modern samples) or ≤ 500 (ancient samples) ambiguous bases. For compatibility with PhyloTree (final v17), slightly modified by the recent L7 discovery (Maier et al., 2022), we included as many representatives of those 5,470 branches as possible (Table **S2**). In cases where samples originally picked by PhyloTree were missing, incomplete, or otherwise incompatible with their branch, we chose from available substitutes (n=72 branches; n=77 samples). Requisite samples were available to attempt reconstruction of 5,448 out of 5,470 (99.6%) PhyloTree branches, although this does not guarantee their existence in Mitotree (e.g., if new samples infer a different topology). Eighteen PhyloTree branches were represented by aDNA samples and therefore were given the same priority as modern samples. Otherwise, aDNA samples were given second priority, for both topological reconstruction and age estimates. All aDNA samples were required to have an estimated mean age, and modern samples were given a default birth year of 1950 unless known. The outgroup was the inferred Reconstructed Sapiens Reference Sequence (RSRS; Behar et al., 2012), which represents the root state of the modern human mtDNA tree.

Previous concerns have been raised about variability in data quality for GenBank sequences (Harris, 2003; Yao et al., 2009). Therefore, we used the following criteria to screen and exclude problematic samples: (1) more than 3 heteroplasmies, (2) more than 10 ambiguous bases, (3) more than two missing upstream mutations (excluding hypervariable sites), (4) more than one heterozygous upstream mutation (excluding hypervariable sites), (5) more than two mutations that are RSRS × rCRS differences (excluding hypervariable sites), or (6) entire studies if more than 10% of samples are in haplogroup H2a2a, indicating extreme rCRS bias.

All subsequent pipeline steps described below—from sequence generation/alignment through haplogroup naming—are fully automated and can be rerun without human supervision as new sequences are incorporated.

### NGS data pipeline

FTDNA customer sequences were generated as previously described (Maier et al., 2022). When next-generation sequencing (NGS) data were available from modern (e.g., 1000 Genomes) and aDNA studies, FASTA consensus sequences were generated using pipelines adapted from the GATK Best Practices for mitochondrial short variant discovery (GATK Team, 2025), using GATK v4.6.2.0, BWA-MEM v0.7.17, and BCFTOOLS v1.22. mtDNA-mapped reads were realigned to (1) the standard RSRS and (2) a shifted RSRS offset by 8 kb (placeholder N bases removed). Variants were called in haploid mode. Modern samples were called using GATK *HaplotypeCaller* with parameters -ERC BP_RESOLUTION --output-mode EMIT_ALL_ACTIVE_SITES. Ancient samples were called using BCFTOOLS multiallelic caller, requiring minimum base and mapping qualities of 30. To mitigate post-mortem damage (PMD) artifacts, sites showing low-level C→T or G→A heterozygosity (minor T/A allele <20% of total depth) had their genotype quality (GQ) scores adjusted to prevent erroneous no-calls. The resulting VCF files from both alignments were lifted over to standard RSRS coordinates using GATK *LiftoverVcf* and merged using GATK *MergeVcfs*. Sites with a minor allele depth/total depth (AD/DP) ratio >0.2 were recoded as heterozygous, except for C/T and G/A sites in ancient samples where the minor allele was A or T; these were retained as homozygous for the major allele to avoid PMD-induced artifactual heterozygotes. Modern sample sites with QUAL <20, DP <1, or GQ <20, and ancient sample sites with QUAL <10, DP <1, or GQ <13, were set to no-call. Consensus FASTA sequences were generated using GATK *FastaAlternateReferenceMaker*, incorporating IUPAC ambiguity codes for heterozygous sites and masking no-called positions as ‘N’.

### Alignment and variant calling

We used MUSCLE V3.8.31 (Edgar, 2004) for sequence alignment and reversed both query sequence and reference to achieve right-alignment of INDELs for backwards compatibility. Default parameters for *-gapopen*, *-gapextend*, and *-maxiters* were used. We locally realigned regions such as chrM:54–71, chrM:455–459, and chrM:16183–16184 which would otherwise give inconsistent alignments due to homopolymers or multiple compensatory INDEL mutations (Table **S3**). For example, insertions following the poly-C region positions chrM:455 and chrM:16183 were normalized to be right-shifted. The complex region between positions chrM:54–71 was locally realigned based on evaluating a preliminary tree without those mutations, then creating some rules to support consistency of homologous INDELs/TRs across samples. Haplogroups W5b, R0a, H15, and A2 alignments were particularly affected by multiple INDELs interacting with SNPs.

Homopolymers and Short Tandem Repeats (both hereafter referred to by the umbrella term “tandem repeats”; TRs) with clear phylogenetic signal were scored from INDELs. We used a set of merge rules to normalize the observed sequence into an ordinal value of repeats (Table **S4**). The following TRs were scored using the listed RSRS value: 71G[6], 291A[6], 299C[2], 455T[4], 459C[4], 463C[3], 498C[5], 502C[3], 524AC[4], 2232A[6], 4322C[5], 5752A[6], 6425C[6], 7471C[6], 8289CCCCCTCTA[2], 12241C[5], 12310A[5], 15944T[5], 16186C[3], 16296C[3]. The following INDELs were not scored as TRs, but rather listed as “XC” (binary) INDELs: 309.XC, 315.XC, 573.XC, 960.XC, 965.XC, 5899.XC, 8276.XC, 8278.XC, 8287.XC, 16193.XC. Table **S5** shows all INDELs and TRs used in our analysis. The following positions were entirely excluded due to well-documented alignment ambiguity or extreme mutational instability rendering them phylogenetically uninformative (Bandelt et al., 2002; Yao et al., 2009): 309–310, 523–524, 573–574, 576, 3107, 16182–16183, whereas 16093 was only excluded for SNPs, and the following were only excluded for INDELs/TRs: 311–315, 502, 514–522, 960, 965, 3105–3106, 5899, 16184, 16187, 16189, 16193 (Table **S6**). In addition to excluding some markers, other hypervariable yet informative markers were “second-tier,” to be considered only after a preliminary tree was built with primary (“first-tier”) markers.

Although marker weights (maximum parsimony) or models of evolution (maximum likelihood) can account for molecular rate variability, heterotachy amongst markers (local rate changes in specific lineages) present a greater difficulty. We introduced a concept of “COMBO markers” to locally increase the weights of markers whose globally defined weights would otherwise be insufficient to compete with conflicting markers (Table **S7**). Similar ideas have been proposed to bolster the weight of groups of markers (Capella-Gutierrez et al., 2014), hence the combination of coevolving markers was preferred over a one-dimensional global weight. In some cases, otherwise informative markers were unreliable in specific clades, in which case we locally excluded one or more alleles at those positions (Table **S8**).

### Constraint tree analysis

Given the nearly unprecedented size of the phylogenetic problem, yet the need for consistency, accuracy, and speed (and frequent re-analyses), we used a divide-and-conquer approach (Fig. **3**). During early experiments, we tested the NJML method (Ota and Li, 2000) which partitions the tree with a neighbor joining (NJ) step, with subsequent intense searches by were larger than ideal, and ML searches using IQ-TREE2 v2.1.3 (Minh et al., 2020) were too slow for 100,000s of sequences. Instead, we used heuristic parsimony-based searches for both steps in TNT v1.6 (Goloboff et al., 2008), for smaller partitions and greater speed. First, we used TNT’s *piñón fijo* method, which is a type of cyclic perturbation based on the earlier technique of *Recursive Iterative-DCM3* (Roshan et al., 2004; Goloboff and Pol, 2007). Essentially, a starting tree (based upon one TBR replicate) is partitioned into reduced datasets, where Hypothetical Taxonomic Units (HTUs) are given the correct ancestral states with only 0.95 probability (Goloboff, 2015). By swapping this perturbed tree with several rounds of TBR and drifting, then reattaching to the larger tree, the algorithm efficiently ratchets huge trees toward a more parsimonious state. We performed 100 bootstrap support (“BS”) replicates, and 100 replicate support (“RS”) reps with different starting seeds, in addition to the main replicate. We further reduced computational load in this and subsequent analyses by de-duping sequences to “meta-haplotypes,” or identical haplotypes after ignoring markers on the exclude list (22,862 / 177,196 = 13% reduction).

**Figure 3:**
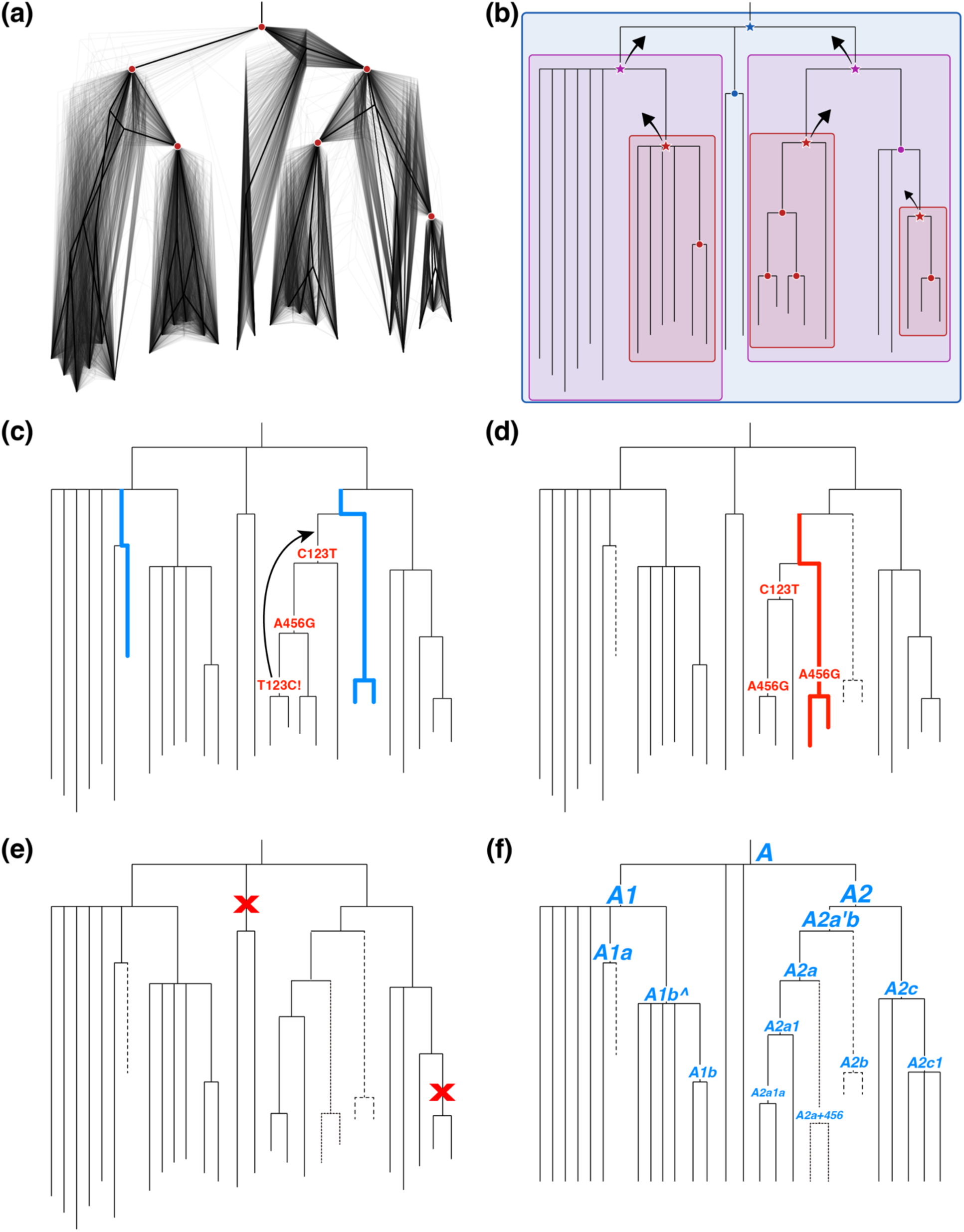
Methodology Pipeline: Major steps of Mitotree’s unique divide-and-conquer strategy: (a) a constraint tree is built using hundreds of replicates, to choose confident partitions; (b) more intensive weighted parsimony is applied recursively within each partition, using a bottom-up traversal (starred nodes each become one sequence); (c) ancient DNA samples (blue branches) are added secondarily and allowed to branch swap; (c) *also* illustrated is the pre-correction state of a reversal (red mutations) to compare with panel (d), which shows the *post*-correction state of 2-step reversals (red branches = new location of reversal-clade sequences); (e) weak branches are collapsed based on selected criteria; (f) TMRCAs are estimated (tree is now ultrametric), and branches are named.

Constraint nodes were defined as having either (1) 0.9 BS support, (2) 1.0 RS support in clades larger than 5% of the total tree, or (3) membership to certain special haplogroups, based on containing COMBO markers for subclades of H (H1, H2, H3, H5’36, H7, H10, H29, H55, H66) or HV0a (V). Criteria (1–2) ensured that only strongly supported nodes could partition the tree, and criterion (3) ensured partitions from huge clades (e.g., H1 has nearly 20K haplotypes) with strong support from previous research, but variable node support due to polytomous nature of branches and a few uncertain sequences. Constraint nodes were required to have a minimum of five sequences to ensure that intensive phylogenetic searches were of a useful size. Typically, this resulted in ≈3% of the final tree consisting of constraint nodes.

We used these 101 preliminary constraint tree RS replicates to further prepare marker weights for the more intense, recursive phylogenetic step (described below). Weights were calculated from median counts of mutation occurrences across trees using the *map* command in TNT, which uses two-pass Fitch optimization on DNA and INDEL characters, and Farris optimization on ordinal TR characters (Goloboff, 1993a; Goloboff et al., 2008). Letting 𝐜 be the vector of mutation counts with elements 𝑐*_i_*, then we calculated an intermediate vector of weights 𝐰 with elements:

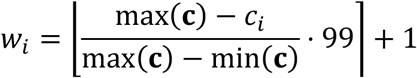

And transformed 𝐰 into a final vector of weights 𝐳 with elements:

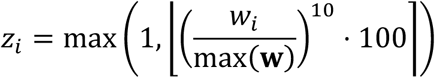

The exponential transformation effectively compressed the weights toward lower counts, to accommodate a log-normal distribution with heavy right skew. Hypervariable markers with outlier count values received weights close to 1, and strong markers spanned up to 100.

When calculating ancestral state changes for weights and subsequent pipeline steps described below, we made several adjustments for ambiguities. Biallelic ambiguities (e.g., C/T or 0/1) spanning multiple HTUs were resolved upstream, and triallelic ambiguities (e.g., C/T parallel to C/A) were resolved as two downstream biallelic ambiguities. Sequences with heteroplasmies (HETs; e.g., C16184M) were sometimes excluded from clades immediately downstream (e.g., C16184a) given that TNT’s correct parsing of IUPAC ambiguity codes can dissolve otherwise well-supported clades when even a single sequence carries a HET at a clade-defining position. We automatically moved these sequences downstream in unambiguous situations where the HET was clearly compatible with one downstream clade. All pipeline operations were written in R v4.5.0 (R Core Team, 2026), primarily using the phylogenetic packages APE v5.8.1 (Paradis and Schliep, 2019), PHYTOOLS v2.4.4 (Revell, 2024), PHANGORN v2.12.1 (Schliep, 2011), and TREEIO v1.32 (Wang et al., 2020).

### Recursive phylogenetics

Within each constraint clade, we ran moderately intensive phylogenetic searches using TNT’s weighted parsimony, with the function *xmult* set at the maximum search intensity of 10. This heuristic search strategy combines sectorial searches, drifting, ratcheting, and fusing to perturb the tree while performing TBR branch swapping, to optimize its parsimony score and escape local optima (Goloboff, 1999). We employed a moderately conservative but thorough search strategy for sectorial searches (*sectsch: selfact 50 maxsize 500 godrift 250 fuse 2 drift 8 slack 100;*) and drifting (*drift: rfitdiff 0.1 fitdiff 2;*). The *ccode* function activated marker weights for parsimony searches and allowed TR markers to be treated as additive (ordinal). Outgroups were selected by traversing the ancestral constraint nodes, and for each upstream sibling branch, choosing two sequences from independent clades. Including a ladder of multiple close outgroups helps to polarize ancestral from derived states (Nixon, 1993). The outgroup topology was enforced using TNT’s *force* and *constrain* functions, which have recently been optimized to allow defining some taxa as floaters (Goloboff and Maier, in review). Embedded constraint clades within the focal constraint clade were removed and replaced by a single HTU sequence (Fig. **3b**). We used all downstream taxa (i.e., no HTUs informed by other HTUs), and the same ancestral sequence reconstruction (ASR) method described above except that ambiguities were kept.

We performed 20 BS and 20 RS replicates in addition to the main replicate. Out of the 21 identically run replicates (20 RS + 1 main), we chose the best tree based upon maximum parsimony (MP) score, but used maximum clade credibility (MCC; APE’S *maxCladeCred* function) score to break ties. Although MCC is typically applied to Bayesian posterior distributions, we use it here as a tiebreaker among equally optimal parsimony trees, selecting the tree whose bipartitions are most consistently recovered across replicates. We considered ties to include cases where polytomous (collapsed) MP scores were closer to each other than the maximum difference between polytomous and dichotomous MP scores, suggesting uncertainty in how tree optimization of the latter corresponds to quality in the former. We assigned BS and RS values to the best tree by counting bipartitions across replicate trees using APE’s *prop.part* and *prop.clades* functions. We designed an additional measure of node support termed “mutational support” (MS), defined as having the same mutational signature (including upstream first-tier mutations), or just having the same branch-specific mutations, plus sharing 95% similar taxa. This measure is less sensitive to several rogue or uncertain taxa flipping between clades. We used the *digest* function from the DIGEST v0.6.37 package to create an md5 hash for efficient branch comparisons.

Rapid reversals (i.e., mutations occurring and then reverting two steps downstream) are often caused by inter-marker conflict, and a flat parsimony landscape for optimizing tree length. Although TNT does not discourage rapid reversals compared to parallel mutations, intuitively, the former usually make less sense, when a weaker character is sandwiched between two (forward/reverse) occurrences of a stronger character. We further develop this argument below, but here describe the first (“pre-correction”) of two steps for minimizing rapid reversals. Rapid reversals were identified on the best tree and all RS trees, then prioritized by (1) weight, (2) size of clade, (3) number of occurrences. For each reversal, a section of tree was excised and replaced by a donor RS tree section wherever possible to eliminate or reduce the issue. Once replaced, these tree sections were flagged and removed from further modification.

Once all steps described were performed for first-tier markers, the resulting topology was enforced as a tree constraint using TNT’s *force* and *constrain* functions, while second-tier mutations were introduced to potentially add additional branches. This allowed more hypervariable (but still informative) markers to refine the topology, but only after giving priority to more stable mutations, and avoiding some phylogenetic conflict. The second-tier step added another 41 replicates (20 BS, 20 RS, 1 primary) for a total of 82 replicates per constraint clade. Constraint clades were recursively run from tips to root. After all recursive levels had been finished, the combined tree was annotated with their original BS/RS/MS support values (since constraint nodes would have received 1.0/1.0/1.0 values in the last step). Constraints were quality-checked for adherence.

### aDNA (second-priority samples)

Due to the potential for aDNA sequences to contain artifacts, excess HETs, and low or variable sequencing depth, we deprioritized their influence on the overall phylogeny. We added 2,579 aDNA haplotypes to be branch swapped across the existing topology reconstructed from first-priority samples (Fig. **3c**). These excluded the 18 previously mentioned aDNA samples vetted to represent the original PTv17 structure and given first-priority status, and also 559 duplicates. We used TNT’s *hybrid!* function (Goloboff and Morales, 2023), which treats the existing modern-DNA topology as a fixed subtree, sequentially adding aDNA via Wagner addition, and then TBR-swapping these new taxa as “floaters” across the fixed modern structure. This was performed for 100 BS, 100 RS, and one main replicate again, with newly generated MS scores, and finally the previously calculated BS/RS/MS scores were carried forward.

### Reversal correction

When two explanations of phylogenetic conflict (i.e., reversals vs. parallel mutations) have nearly equal weighted parsimony scores, the tiebreaker should not be arbitrary but rather should reflect an underlying probability behind each explanation. Rapid reversals (which revert two steps downstream) often appear as weaker characters sandwiched between forward/reverse occurrences of a stronger character, yet the optimality criterion should systematically prefer the explanation that invokes homoplasy in the faster-evolving (weaker) character (i.e., Camin-Sokal parsimony; Camin and Sokal, 1965). Although weights, COMBO markers, and local exclusions help mitigate this problem, heterotachy and imperfectly understood weight-rate relationships can still produce poorly supported rapid reversal structures.

We used the following criteria for reversal correction: (1) weight >10 for reversing mutation, (2) exactly one first-tier mutation in the middle, (3) either (a) exactly one middle mutation (ignoring hypervariable mutations of weight 1), or (b) multiple such mutations, with a combined weight ≤ the reversing weight. We also ignored reversals spanning two constraint clades. These criteria identify and correct up to 6-step reversals with flimsy branches in between. To minimize the tree size computational burden, we again partitioned the tree into chunks containing a mutation and all its rapid downstream reversals. To ensure accurate reconstruction of HTUs, we included nodes up to three steps further downstream if they contained any of the relevant mutations. Chunks that shared adjacent nodes were merged, then outgroups were selected as above, and HTUs were reconstructed using the same ASR method as above.

Reversal correction was performed by positively constraining the entire topology, except that the sandwiching clades (forward/reversal branches) were left unconstrained (Fig. **3c–d**). The middle clade was negatively constrained to disallow testing of the reversal clade(s). Similar sandwich structures could still bypass these constraints (if tips varied slightly) so this algorithm was run iteratively across each chunk until stability was achieved. Stability was defined as one of: (1) zero qualifying reversals, (2) no reduction in such reversals after trying new negative constraints 3×, (3) no reduction in such reversals, and no new negative constraints to try, or (4) an increase in reversals (then reverting to the previous iteration and stopping). Given that heuristic tree searches are non-deterministic and may find different local optima across runs, we repeated this entire process 10× and selected the tree with fewest qualifying reversals for each chunk. Using the final set of positive and negative constraints, we followed this with 20 BS and 20 RS replicates and MS score calculation, then aggregated BS/PP/MP scores for newly created nodes, and finally plugged chunks back into the main tree. This reduced qualifying reversals from n=315 to n=11 (96.5% reduction) across the tree. This n=11 reflects only the immediate post-correction state; “weak branch collapse” (below) can re-introduce sandwich structures during mutation re-optimization, yielding higher final numbers (n=117; see Results).

### Sibling merge

One final artifact to potentially correct is the occurrence of identical sibling mutations under the same ancestral node, either due to separation of constraint analyses, or the reversal correction algorithm. We corrected three scenarios: (1) two sibling branches with one identical mutation only were merged, (2) if sibling branches X and Y shared one mutation, but Y had other mutations, then XY formed a new branch with Y as subclade, and (3) if sibling branches X and Y shared one mutation, but both X and Y had other mutations, then XY formed a new branch with both X and Y as subclades. We only considered first-tier mutations, and in the rare case that multiple pairs of sibling mutations affected one or more of the same branches, the mutation with highest weight was chosen for correction. A total of n=6 sibling branch pairs were affected by these rules across the tree.

### Meta-haplotype re-integration

No further branch formation or heuristic tree searches were allowed after the sibling merge step, and hence tree size was no longer a limitation. We attached missing meta-haplotypes as terminal branches onto their corresponding parent branch and then used another round of ASR to identify and collapse nodes supported by zero or ambiguous mutations. Out of 22,862 branch splits, 3,930 were retained for further consideration. We categorized all remaining nodes into five categories, according to which step created them: (1) constraint nodes, (2) first-tier nodes, (3) second-tier nodes, (4) aDNA nodes, or (5) meta-haplotype nodes.

### Weak branch collapse

We iterated through several cycles of checking branch mutations, collapsing weak or otherwise unsupported branches, then rechecking until all criteria were fulfilled (Fig. 3e; Table **S9**). First, we performed ASR with our defined set of mutations on the PTv17 topology for the purpose of comparing between trees, and then we ran ASR on pre-collapsed Mitotree. Several categories of mutation instability were defined based on occurrence across the tree: global occurrence (GO) quantile; local occurrence (LO) quantile, i.e., occurrences downstream of the focal node; and (3) local recurrence (LR) quantile, i.e., local occurrences followed by one or more reversals. Collapse criteria included the following types of nodes: (1) zero mutations, or only uncertain mutations, (2) only second-tier mutations with GO >90th percentile (*P*_90_), and best node score max(BS,RS) ≤0.5, and either LO >*P*_90_ or LR >*P*_0.1_, (3) second-tier nodes with max(BS,RS) of 0.0, (4) any nodes with max(BS,RS,MS) of 0.0, (5) only COMBO markers (ignoring uncertain mutations), or (6) only hypervariable (weight of one) SNP transitions on second-tier or aDNA branches, in older clades. Since no ages had yet been computed, we calculated the mean path length (MPL) as a proxy (i.e., the average number of first-tier mutations from the focal node to each modern leaf), and applied a minimum threshold MPL of 2.0 ≈ 10,000 ybp.

We added several exceptions to these collapse rules. Unique SNP matches to PTv17 branches were saved from collapse, ignoring uncertain PTv17 branches (denoted by SNP-based names). For example, haplogroups L1b1a10, M13a’b, M7c1c3, U3a2a, U5a1a1d, U5a1b1, U5b2b4a, and U6a’b’d were saved from collapse by this rule, but haplogroups such as U4b1-T146C! were collapsed by rule (6). Another exception was for branches with only COMBO markers, rule (5), which occasionally form if several samples lack the COMBO marker for their branch (due to missing upstream requirements). Usually, rule (5) automatically rectified this situation by merging a downstream COMBO-only branch with its upstream (correct) branch. Rarely, such a collapse would instead destroy the desired branch, due to upstream homoplasy. For example, by automatically collapsing the branch containing only the H1 COMBO marker <H1:2706A+3010A>, H1 would be destroyed, and its subclades merged with a false upstream branch defined by 3010A (which evolved in parallel several times, in H30b, H41, H65a, H79a, and H84). We rectified this by detecting which of the two branches had more leaves in common with the desired branch (H1), and collapsing the false one.

### Branch length estimation and age estimates

Branch lengths were estimated as the sum of linear mutation weights on each branch. We benchmarked this method against using IQ-TREE2 (evolutionary model for SNPs: TIM3+F+I+G4; INDELs: GTR2+F+ASC+G4; TRs: MK+ASC+G4) and found nearly identical results (*r* = 0.99), but IQ-TREE2 was unable to complete on larger datasets. We counted HETs as ½ weights given their incomplete state as substitutions. Branch lengths were normalized to a unitless value of 1.0 for the longest branch. We used the relaxed clock model of LSD v1.9.9 (To et al., 2016) to estimate times to most recent common ancestor (TMRCAs) of each branch (Fig. **3f**). The LSD model uses weighted least squares to simultaneously minimize errors on every branch, while penalizing longer branches the most, because they tend to reflect the biggest rate changes.

The root date was estimated by BEAST v2.7.7 (Bouckaert et al., 2014) to be 143,000 ybp, and we used a minimum of one generation (30 years) between branches. Tip dates were set to birth years (if known) or else a default of 1950 CE. Branches were constrained to a minimum age of 1800 CE. Using the *v=2* option, LSD was run twice: once to estimate branch lengths, and again to calculate variances. Variance of branch length 𝑏*_i_* used to distribute relaxed clock error is 𝑣*_i_*:

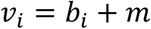

where 𝑚 is the median branch length. The confidence intervals of the branch lengths (𝑐*_i_*) were estimated from 100 simulated trees:

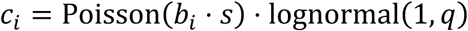

where 𝑞 (the SD of the relaxed clock) was estimated from BEAST (ucld.sdev = 0.208; coefficient of variation = 0.21; Maier et al., 2022), and 𝑠 was the sequence length 16,569. We performed these age estimates initially on modern samples only, to prioritize higher quality sequences.

Using the modern TMRCA estimates as constraints, we reinserted aDNA samples for a second round of LSD estimates. Due to the potentially artifactual nature of aDNA private variants we pruned aDNA samples down to their inner nodes, which served as terminal branches with minimum age constraints based on the C_14_ dates of the archaeological samples. We checked for any paradoxes between modern and ancient constraints and iteratively removed any modern node constraints younger than known ancient sample dates. We approximated the final set of 95% confidence intervals using Gamma distributions:

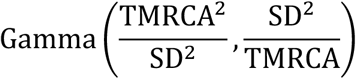

### Variant annotation

Most variant types were annotated identically to previous conventions for backwards compatibility (Table **1**). For example, SNPs were given ancestral and derived states, in addition to position number, lower case letter to denote transversions, and IUPAC codes to denote uncertainty in ancestral state (e.g., for a complex triallelic change), or represented a HET for observed sequences. Typically, branch uncertainties were biallelic and simply represented by parentheses, denoting uncertainty in which branch contains the synapomorphy, e.g., (A56G). Deletions were given an ancestral state and position, or range positions, followed by a “d”, whereas insertions followed the original position with the added base(s). Complex insertions included homopolymers too difficult to normalize into TRs and hence scored as binary “XC” mutations (e.g., 8287.XC), and long insertions (e.g., 480.1TCCCATACTACCAAT). TRs were mostly homopolymers and encoded with the ancestral and derived counts in square brackets, e.g., [6]71G[5], where “71” refers to the last position of the ancestral repeat. The INDEL at position 8289, originally described by three variants 8281-8289d, 8289.9bpINS, and 8289.18bpINS, was encoded as a TR, with corresponding values [2]8289CCCCCTCTA[1], [2]8289CCCCCTCTA[3], and [2]8289CCCCCTCTA[4]. Reversals were denoted by one “!” per reversal event between root and branch. In rare cases where uncertain SNPs occurred at the same position as confident deletions of the same position, the SNP was ignored.

**Table 1.** Variant annotation.

| Class | Type | Example | Example Reversal |
| --- | --- | --- | --- |
| SNP | Transition | G15930A | A15930G! |
| SNP | Transversion | A16265c | C16265a |
| SNP | Reversal | T152C! | C152T!! |
| SNP | Ambiguous Branch | (A56G) | (G56A!) |
| SNP | Ambiguous State | A16265V | V16265A! |
| INDEL | Deletion 1 bp | A249d | d249A! |
| INDEL | Deletion 2+ bp | 59-60d | d59-60TT! |
| INDEL | Insertion 1 bp | 60.1T | T60.1d! |
| INDEL | Insertion 2+ bp | 57.1CT | CT57.1d! |
| INDEL | Insertion Homopolymer | 8287.XC | C8287.Xd! |
| TR | TR 1 bp Motif | [6]71G[5] | [5]71G[6]! |
| TR | TR 2+ bp Motif * | [2]8289CCCCCTCTA[1] | [1]8289CCCCCTCTA[2]! |

\* Only 1 bp and 9 bp TRs were actually used (others were excluded)

### Branch naming

Preserving continuity with decades of literature discussing mtDNA haplogroups was a priority for the Mitotree naming system. Haplogroups were named to maximize backwards compatibility with PTv17 naming conventions and minimize the need for future renaming. PTv17 “root” names (A–Z, L0–L7, CZ, HV, and JT) were assigned to the best-matching Mitotree haplogroups based on the samples they contain, and if no unambiguous mapping could be found, by best matching variant signature. Root haplogroup B, which was polyphyletic in PTv17 but monophyletic in Mitotree, was mapped to the most recent common ancestor (MRCA) of B4, B5, and B6. Each remaining Mitotree haplogroup was assigned to a root based on its placement relative to the root haplogroups.

Four name types were used in addition to the root names (Fig. **3f**). (1) “Primary” names consist only of alphanumeric characters (e.g., A2, B2b2) and form the core of the naming structure. “Non-primary” names (2–4) are used for haplogroups surrounding primary names to preserve the name structure and avoid reassigning names to entire clades when the tree is refined over time. (2) “Inner” names are used for joining multiple descendants with primary names, (e.g., L5’7). A single apostrophe may join up to five name units and up to a total length of ten characters for the joined string (e.g., 1’2’3’4’5 is five units and nine characters). For brevity, longer inner names use double quotation marks between the lowest and highest name unit (e.g., M13”108, L0a”m). (3) Variant names follow the shorthand format used by HAPLOGREP v3 (Schönherr et al., 2023): the parent haplogroup name, a plus sign, and the defining variant position (e.g., H2a1+16354). For haplogroups defined by multiple variants, the lowest position is selected, with preference for stable variant positions over second-tier positions. Variant names are also given to any haplogroup joining more than ten names to avoid instability as new lineages are discovered. (4) “Interim” names denote a split of a haplogroup with a primary name and use one or more caret symbols (^) following the downstream primary name (e.g., B2^, B2^^). When a future child haplogroup is formed and assigned a primary name, the interim haplogroup is renamed with an inner name (e.g., B2’9). Name type combinations (e.g., A2ad”ei^, X2l^+152, V57’64+150) are used when applicable.

Within each root haplogroup, haplogroups were named recursively, top-to-bottom, relative to the PTv17 names. Primary names were mapped to the Mitotree haplogroup with the most similar variant signature, and in ambiguous cases, by identical samples. PTv17 haplogroups that were absent from Mitotree were ignored, but their names were reserved to prevent reuse. Second-tier haplogroups and homoplastic haplogroups (defined by a reversal or convergence within 3 branch steps), which are more likely to change in future versions, were assigned non-primary names (e.g., J1b+16261), unless they had previously been assigned a primary name (e.g., CZ or K1c). Previously assigned names in published mtDNA trees (PhyloTree, Mitotree, YFull MTree) or the literature were avoided when assigning new haplogroup names, and preferred when assigning names to haplogroups that matched the published variant signatures, to avoid confusion and maintain continuity.

The root of the modern human tree, encapsulating all modern human clades L0–L7 is named “L.” In the PTv17 convention, after the “z” letter is reached in an alphabetic series, “aa” through “az” are used, and so on. Applying this same convention to the root letters, we reserved root haplogroup names AA and AB for the Neanderthal and Denisovan archaic human clades (Fig. **4**; Fu et al., 2025; Meyer et al., 2014; Sawyer et al., 2015).

**Figure 4:**
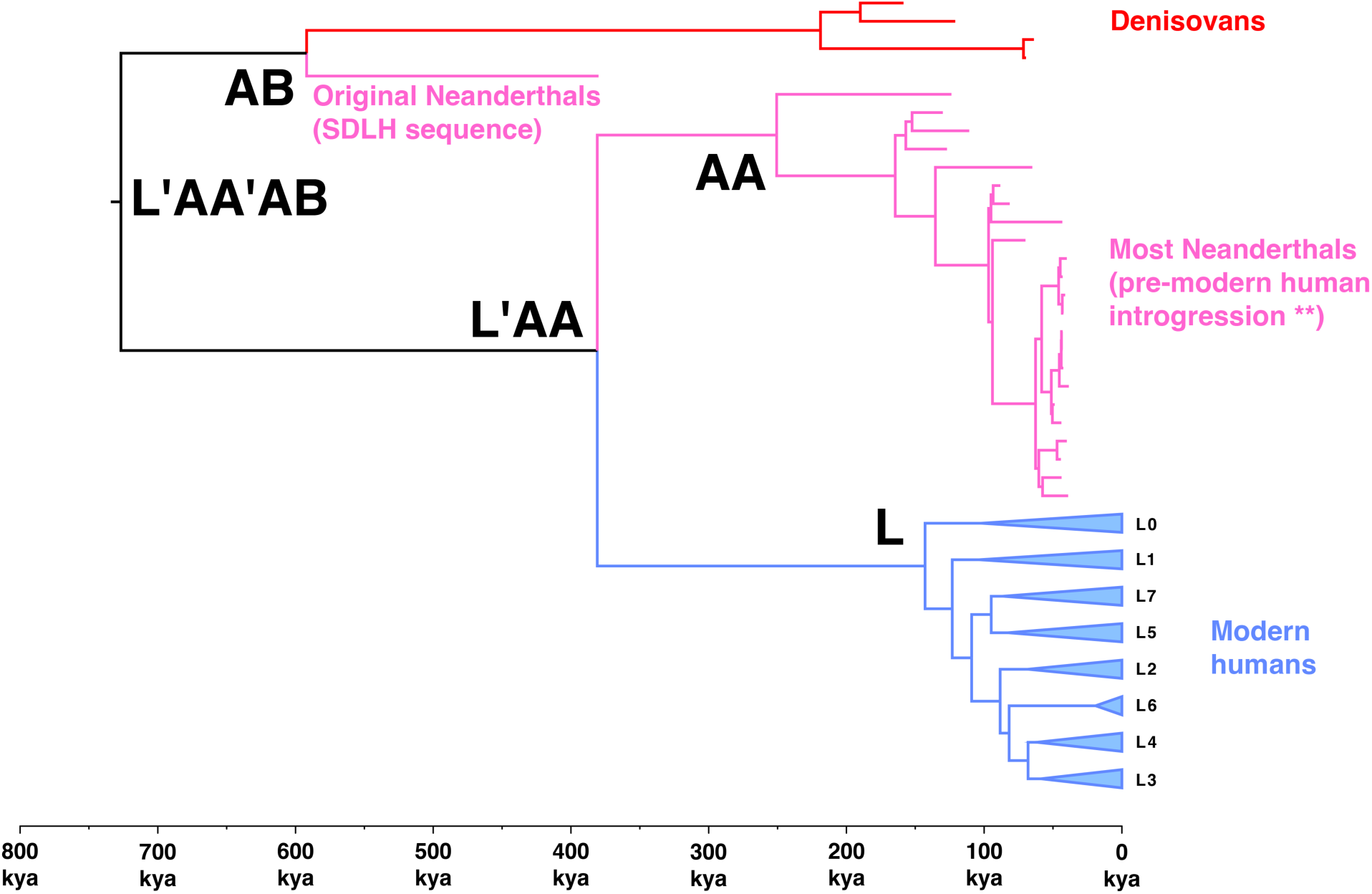
Neanderthal and Denisovan Reserved Haplogroups: Following PTv17 convention (“aa” follows “z” in an alphabetic series), root haplogroup names AA and AB are reserved for Neanderthal and Denisovan clades. Although AA is closer to modern humans (L) than Denisovans (AB), this is likely because most Neanderthals carried pre-modern human mtDNA via ancient introgression (Petr et al., 2020). The Sima de los Huesos (SDLH) sequence may represent an “original” Neanderthal mtDNA motif.

### Validation approach

To assess false-positive (FPR) and false-negative (FNR) branch recovery rates, we simulated phylogenetic datasets modeled on the empirical PTv17 (plus L7) requisite sequence set (∼7,000 sequences; Table **S2**). This minimum dataset for reconstructing PTv17 provides a tractable unit for multiple validation replicates given the multi-day runtime of a full Mitotree analysis. Simulated datasets were generated using mutation rates and branch lengths drawn from the empirical dataset: variants were drawn from a Poisson distribution parameterized by the empirically observed per-marker weights and per-branch expected counts, preserving the site-specific rate heterogeneity and empirical homoplasy distribution of the real data. Heterotachy was not explicitly modeled, meaning simulation conditions were marginally more favorable than real mtDNA data in this respect. Sequencing noise was set to zero, consistent with standard practice in phylogenetic benchmarking (Park et al., 2021; Zaharias and Warnow, 2022) and to avoid introducing arbitrary noise parameters uncalibrated to empirical error rates. Each simulated alignment was run through the full Mitotree pipeline, and the resulting tree was compared against the known ground-truth topology. FNR was defined as the proportion of true branches absent from the reconstructed tree, and FPR as the proportion of reconstructed branches absent from the true tree. We additionally report accuracies excluding second-tier nodes, which are inherently less stable. We performed n=10 replicates and repeated this using FASTTREE2 (Price et al., 2010) as a comparative benchmark, given its status as the only alternative method capable of scaling to 100,000s of sequences.

## Results

### Tree size and stats

We found a total of 53,588 haplogroups, ≈10× larger compared to 5,438 in PTv17 and 5,470 after the L7 discovery (Table **S10**). Broken down by pipeline step: 1,726 haplogroups (3%) were constraint nodes, 41,189 (77%) were first-tier nodes, 5,517 (10%) were second-tier nodes, 633 (1%) were aDNA nodes, 587 (1%) were reversal-correction nodes, and 3,936 (7%) were meta-haplotype nodes (Table **2**). Broken down by branch name, 36 (0.1%) received root names, 1,865 (3%) received inner names, 45,415 (85%) received primary names, 3,497 (7%) received variant names, and 2,775 (5%) received interim names. A total of 335,283 variants were distributed across Mitotree, with 89,653 shared variants, 245,630 private variants (PVs), and 46,679 uncertain ones (either shared or private). Haplogroup ages were consistently younger than PTv17 ages within each root (Fig. **5**), with a median reduction of 2,003 years (PTv17 = 1262 BCE, Mitotree = 741 CE), and a mean reduction of 2,793 years (PTv17 = 2656 BCE, Mitotree = 137 CE), reflecting the increased resolution of terminal haplogroups.

**Figure 5:**
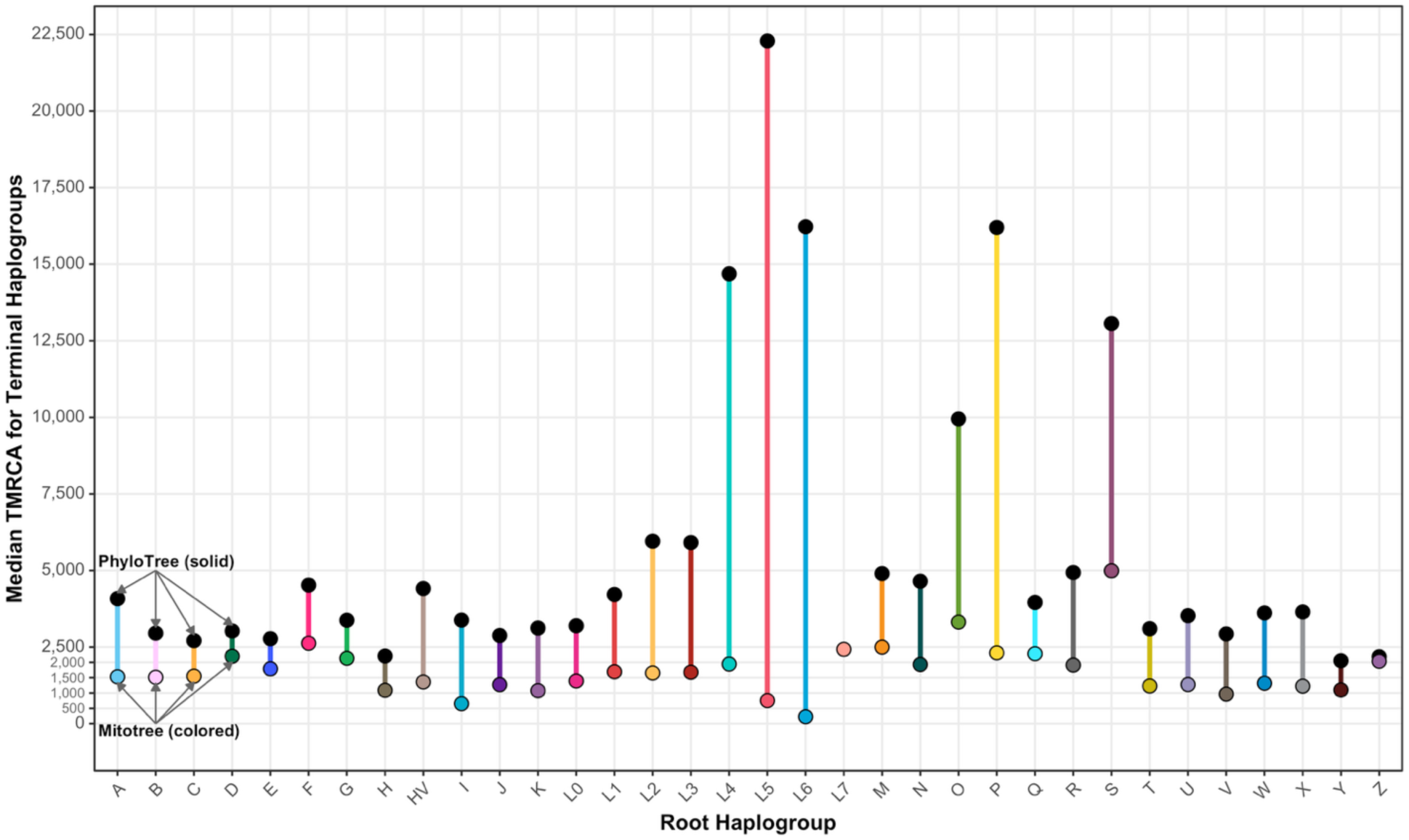
Mitotree’s Resolution of Younger Haplogroups. Median reduction in the age of terminal haplogroups (those without any subclades), shown for each root haplogroup. Comparison of Mitotree to PhyloTree (PTv17).

**Table 2.**
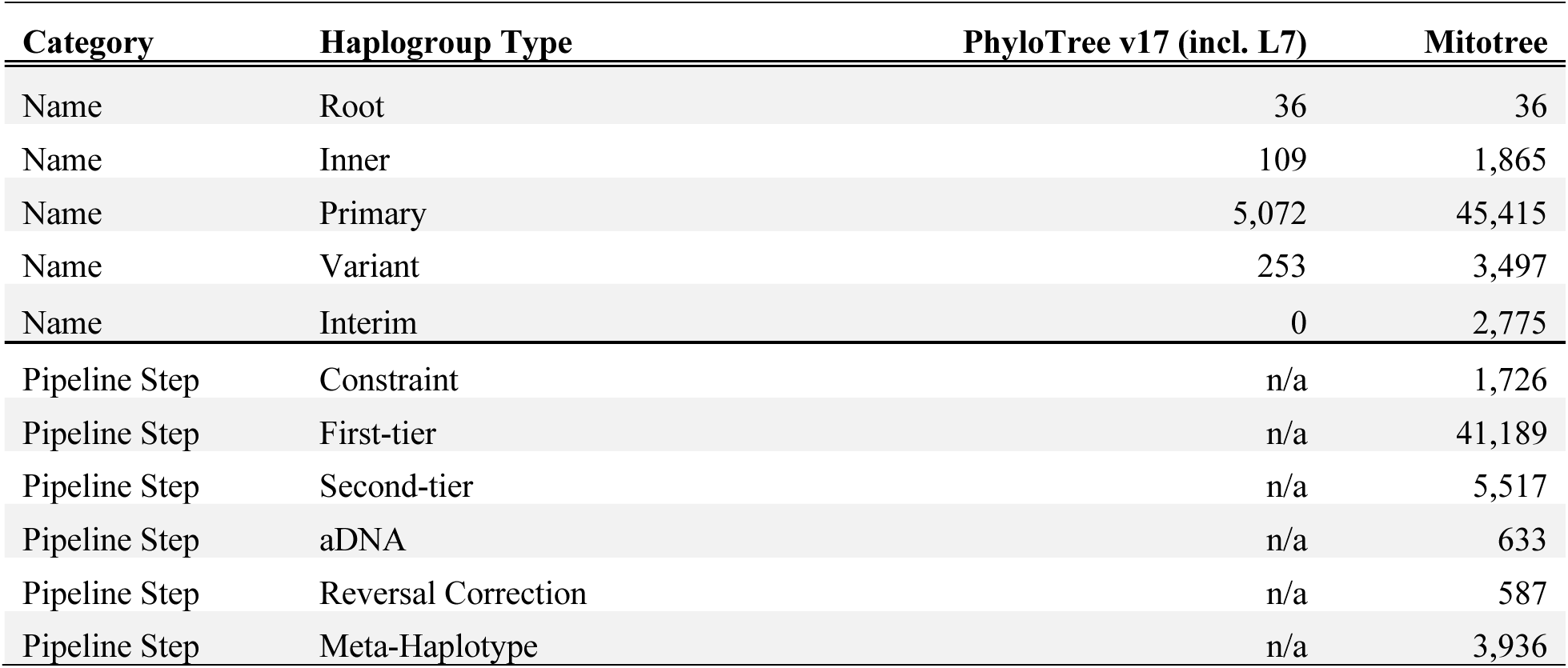
Haplogroup breakdown by name and pipeline step.

| Category | Haplogroup Type | PhyloTree v17 (incl. L7) | Mitotree |
| --- | --- | --- | --- |
| Name | Root | 36 | 36 |
| Name | Inner | 109 | 1,865 |
| Name | Primary | 5,072 | 45,415 |
| Name | Variant | 253 | 3,497 |
| Name | Interim | 0 | 2,775 |
| Pipeline Step | Constraint | n/a | 1,726 |
| Pipeline Step | First-tier | n/a | 41,189 |
| Pipeline Step | Second-tier | n/a | 5,517 |
| Pipeline Step | aDNA | n/a | 633 |
| Pipeline Step | Reversal Correction | n/a | 587 |
| Pipeline Step | Meta-Haplotype | n/a | 3,936 |

We used some QC statistics to assess the quality of the tree and look for any red flags (Table **S11**). We found zero paths of low support (max of BS/RS/MS≤0.1 for two or more adjacent nodes), and only eight individual haplogroups with values ≤0.1. Sibling branches with identical parallel mutations occasionally happen but are usually associated with hypervariable markers that cannot successfully form a confident subclade. When considering moderate levels (n≥5 siblings), 259 haplogroups contained sibling mutations, and only 8 haplogroups when narrowing to first-tier mutations only. Prior to the final tally of 53,588 haplogroups, 1,715 branches were collapsed based on the following criteria (not mutually exclusive): (1) zero mutations (n=286) or only uncertain mutations (n=109), (2) only second-tier mutations with poor confidence of max(BS,RS)≤0.5 and mutation rate as previously defined (n=132), (3) only second-tier mutations with zero confidence for max(BS,RS) (n=105), (4) zero total confidence with max(BS,RS,MS) of 0.0 (n=105), (5) only COMBO markers (n=18), and/or (6) only hypervariable (weight of one) SNP transitions on ≈10,000 ybp branches (n=1,178). Despite applying reversal correction, a small number of qualifying reversals remained for 1-step (n=30) and 2-step (n=87) types.

### Extinct haplogroups

Out of the 5,217 PTv17 (plus L7) haplogroups that ignore unstable variant names (e.g., H1-T152C!), 100 (1.9%) were not recovered (Table **S2**); their names were reserved and will not be reused in future versions (see “Branch naming”). Nineteen of these could not be recovered because at least one of the requisite full or high-quality samples were missing: A9, B4d2, B4j, C7a1a1, D2c, D4h3a7, F1b, H88, M31b2, M38d, M47, M49a2, N14, Q2b, R12, S5, U6a3d, U8c, W5b1. Another three haplogroups also lacked samples, yet a comparable branch was still added: L3d, M80, R0a2e. Other haplogroups were missing due to excluded mutations (B2r: C5899d; B4a1a1a16: A16182t; U4a1a1: 965.XC; U4a2: T310C; U5a2b1: 960.XC), locally excluded mutations (HV4a: C16221T; R5a, R5a2a, R5a2b2: C16266T), samples lacking requisite mutations (HV4a1a3, I5a1c, J1b1b1b, M42’74, M55’77, M5b’c, M73’79, Q1’2, Q1f, R31, U2c’d), downstream missing samples (R12’21: no R12 samples), or restructuring based on mutational conflict (most cases).

A further 79 (1.5%) PTv17 haplogroups were nearly identical yet named differently (Table **S2**). Many of these cases (n=48) contained identical samples, yet were named differently due to novel downstream combinations (e.g., L1c1a’b’d became L1c1a’b’d’g with the addition of new haplogroup L1c1g), downstream restructuring (e.g., M66b now contains all M66a samples, making M66b the closest haplogroup to M66), or collapse of two branches into one (e.g., H1j2 and H1j2a are now a single branch, making H1j2a the closest haplogroup for previous H1j2). Twelve of these had nearly (≥90%) identical samples but differed for the same reasons, and another 19 were <90% identical, but matched the mutational signature of a previous PTv17 haplogroup.

### Notable discoveries

With nearly 50,000 new branches, we choose to highlight notable discoveries in several ways: (1) ancient splits (i.e., new branches bisecting stems of existing branches older than 30kya; Table **S12**), (2) new ancient clades (>30 kya) that are not splits (Table **S13**), (3) former terminal haplogroups (>30kya) that now have subclades (Table **S14**), (4) large polytomy expansions (>30 child haplogroups, and either an increase of 50 or proportional increase of 10 compared to PTv17; Table **S15**), (5) sporadically chosen younger new clades of anthropological interest, and (6) the average reduction in age of terminal haplogroups with the new resolution (Fig. **5**).

#### L: African phylogeography

Mitotree resolves seven deep-rooted L superclades (L0–L7) plus the Out-of-Africa (OOA) descendants M and N(R) (Tables **S17**, **S18**, **S19**), spanning the full demographic history of modern humans within Africa (Table **S16**). African L lineages document distinct foraging populations (rainforest hunter-gatherers, Khoisan-speaking groups, Tanzanian click-language groups), the West African origin and pan-African Bantu expansion, deep Rift Valley continuity, and the OOA staging population. We discuss each L superclade in turn below.

#### L0: deep African haplogroups

L0a is broadly distributed across sub-Saharan Africa with peak abundance in D.R.C. and with L0a2b reaching peak frequency in Mbuti rainforest hunter-gatherers (RFHGs; Batini et al., 2011; Gonder et al., 2007). These autochthonous Mbuti foragers may represent the founders of the original L0a clade, yet lineage extinction and ancient introgression could explain why the surrounding clades are largely comprised of Bantu and Afro-Asiatic populations (Maier et al., 2022). The novel L0a2b structure is surrounded by many haplogroups in L0a2b”l (30 kya), including new haplogroups such as L0a2e, L0a2h, L0a2i, L0a2k, L0a2l from east Africa, Comoros, and Madagascar. The number of L0a subclades is doubled from three to six (up to L0a8 now), with major diversification leading up to the Last Glacial Maximum (LGM; 22–31 kya) during expansion of arid savannah in Africa. The ancient (33 kya) and novel haplogroup L0a5 is distributed across east Africa (Ethiopia and Kenya) and Madagascar. One previously terminal haplogroup, L0a1a2, now contains a polytomy with 30 child clades coalescing to about 6.3 kya, possibly a Neolithic expansion during the tail end of the Green Sahara period (Tierney et al., 2017).

L0b is an extremely rare and old (60 kya) haplogroup found in Ethiopia and Kenya and is a vivid example of demographic bottlenecks in east Africa; previously a terminal haplogroup, we add the first L0b subclades, and they coalesce to around 20 kya during the LGM. L0d and L0k are contained almost entirely within Khoisan-speakers in Botswana and South Africa; we add a 49 kya split (L0d2c^) to L0d2c. L0f is a rare east African haplogroup found in Afro-Asiatic, Bantu, and Malagasy populations (Batai et al., 2013; Pierron et al., 2017). The Paleolithic L0f2 (64 kya) now contains an extremely old split (L0f2’4; 72 kya) based on the novel L0f4 samples from Madagascar (Fig. **6**), and L0f2a contains new ancient subclades L0f2a1^^ and L0f2a2. The 69 kya previously terminal haplogroup L0f1 now contains subclades with a mean age around the LGM (20 kya), and up to 61 kya (three >30 kya: L0f1b’c, L0f1b^^, L0f1c). L0f itself gained two new major subclades (L0f3 and L0f4), with L0f3 (49 kya) containing new ancient clades 31–39 kya: L0f3b, L0f3c. Lastly, the mysterious and extremely rare L0g is known from a few Khoisan and Khoekhoe-speaking individuals (Damara, !Xuun, Nama) of Namibia and surrounding areas (Barbieri et al., 2014a; Chan et al., 2015). This small and recent (2 kya) haplogroup mirrors L0a2b by gaining some context with new surrounding haplogroups L0l and L0m, coalescing in the Later Stone Age haplogroup L0g’l’m^ (41 kya), from a mixture of Bantu and Khoekhoe-speaking groups of southern Africa.

**Figure 6:**
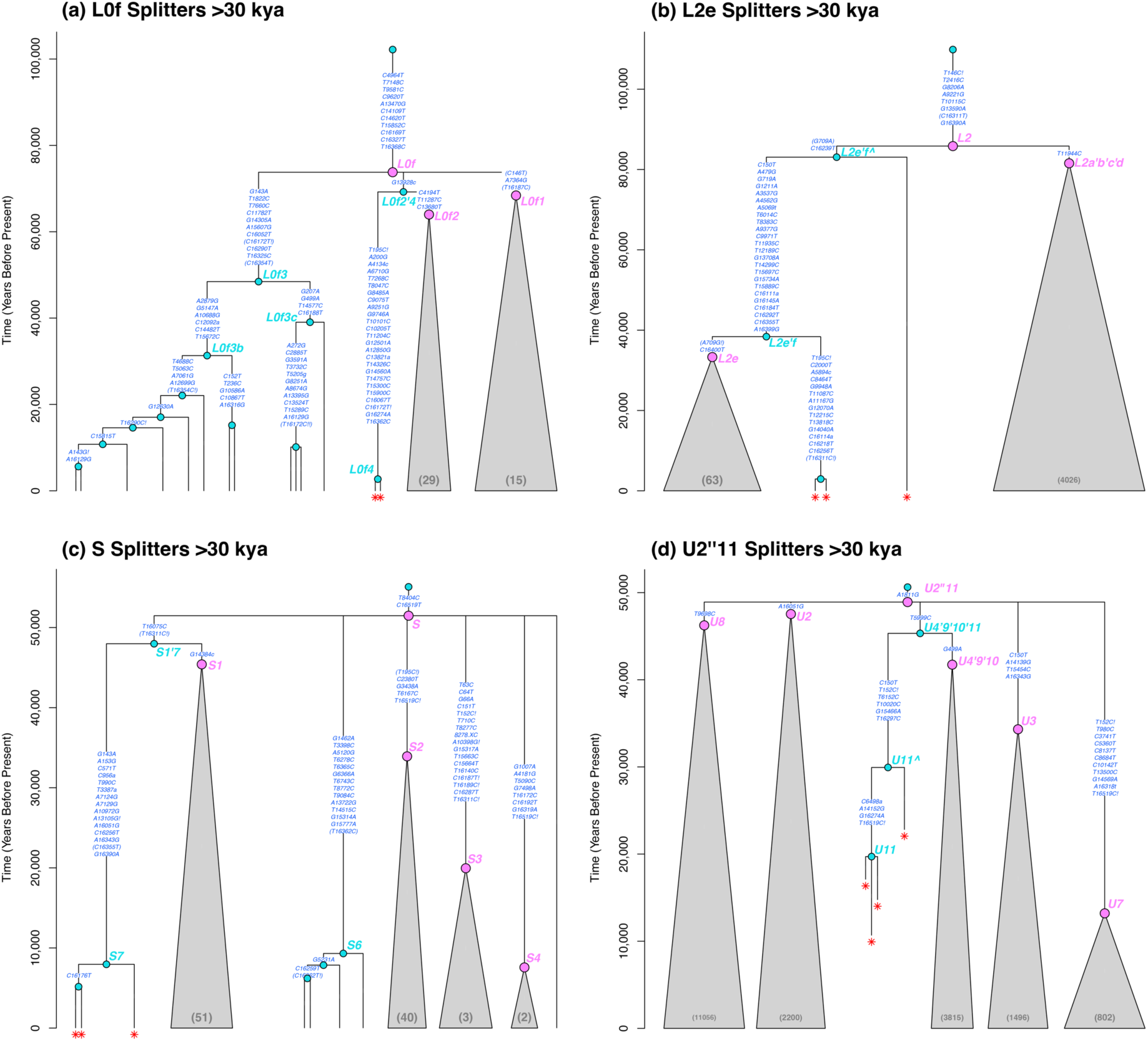
Example Haplogroup Breakers >30 kya: Breakers are defined as one or more novel samples (red asterisks) that create novel haplogroup(s) (cyan) between existing haplogroups (pink). Other novel (non-breaker) haplogroups are also shown in cyan. Shown are: (a) a 72 kya split of L0f2, (b) a double split (83 kya and 38 kya) of L2e, (c) a 48 kya split of S1, (d) a 45 kya split of U4’9’10 by ancient DNA clade U11.

#### L1: western RFHGs (Mbenga and Bedzan)

L1 contains subclades with west (L1b; Senegal, Guinea, Sierra Leone) and central (L1c; Cameroon, Gabon, C.A.R.) African epicenters, carried by primarily Niger-Congo and Bantu populations. L1c reaches its peak diversity on the west coast of Gabon, Cameroon, and Nigeria, and may be prehistorically associated with western RFHGs in the Congo Basin (Quintana-Murci et al., 2008; Batini et al., 2011). It was probably absorbed by Bantu farmers 3–5 kya, as they spread from the Nigeria-Cameroon cradle of agriculture across sub-Saharan Africa. However, the western RFHGs in Mbenga and Bedzan groups (Baka, Bakola, Biaka, Babongo, Mbenzele, Bakoya, Babinga, and Tikar) have relictually high frequencies of L1c1a and L1c4 (Batini et al., 2007). We discovered four major (>30 kya) splits amongst these RFHG lineages. In order of age: L1c4 was split near the root of L1c4’7 at 67 kya, by the novel 59 kya L1c7^; L1c1a1 was split near the root of L1c1a at 46 kya, and gained 36 kya subclade L1c1a1a’c’d; L1c1a1a was split under L1c1a1 at 33 kya; and L1c1a2 was split under L1c1a at 31 kya. L1c1a itself gained 33 kya subclade L1c1a2’4’6’7.

Non-RFHG haplogroups in L1 also became substantially more diverse. Four major splits (>30 kya) were found: the diverse L1c1d, associated with the autochthonous Khoisan (Behar et al., 2008b) and island Malagasy, and mainland Bantu and Niger-Congo groups, was split 40 kya by novel L1c1g; along with L1c1a and L1c1b this forms L1c1a’b’d’g, which was split 65 kya near the root of L1c1 by novel L1c1e; previously terminal haplogroup L1c1c (primarily Niger-Congo) was doubly split near the root of L1c1 (by L1c1f’i) at 45 kya, and again 58 kya by L1c1h; this group gained five haplogroups older than 30 kya (L1c1c4, L1c1c8, L1c1c8a, L1c1c8a^, L1c1c8b; ranging 30–42 kya). Numerous other ancient haplogroups were discovered. Tiny terminal haplogroup L1c6, dated to 56 kya and known previously from two samples (Bantu: Galoa, and Niger-Congo: Kassena), gained a subclade L1c6d at 52 kya. L1c1’2’4’5’6 (now L1c1”9; which unites RFHG, Bantu, and Niger-Congo groups) gained 52 kya haplogroup L1c5’8’9^. L1c3b (Bantu, Nilo-Saharan, Niger-Congo, Malagasy) gained 39 kya haplogroup L1c3b3. L1c2b (Khoisan, Malagasy, Bantu) gained 30 kya subclade L1c2b3. Haplogroup L1c1b, represented by both RFHGs and Gabonese Bantu, gained 30 kya subclade L1c1b1’2^. Finally, we detected a major expansion event during the Green Sahara period in L1b1a3, known primarily from Niger-Congo and Bantu groups spanning Nigeria to Zambia, which increased from 2 subclades into a polytomy of 33 subclades.

#### L7: an aDNA split

After the publication of L7’s discovery, an aDNA split of the new L7a1 branch was found in a 7 kya individual from the Kondoa region of Tanzania, near the peak frequency of modern L7 (Lipson et al., 2022; Maier et al., 2022). This new haplogroup, L7a2, is shared by the ancient Tanzanian, and one modern sample with east African ancestry, 34 kya. L7a1’2 is the new split estimated at 41 kya. The ancient sample’s ancestry was similar to modern Sandawe, who harbor the greatest modern L7 frequency.

#### L2: pre-Bantu deep structure in West Africa

L2 is an ancient haplogroup that probably arose in West Africa ∼90 kya after the end of a western superdrought, when East African hydrological and volcanic conditions were deteriorating (Cohen et al., 2007; Scholz et al., 2007; Blome et al., 2012; Silva et al., 2015). Cyclical pulses of climatic and tectonic instability during that time may have driven human migration back and forth along an east-west gradient, forging the diversity observed in African prehistorical mtDNA (Maier et al., 2022). We discovered the oldest such diaspora in our dataset within L2e (primarily west African), which was split twice: once at 83 kya (Middle Stone Age), and again at 38 kya (Later Stone Age; Fig. **6**). The oldest split dates nearly to the origin of L2 itself (86 kya). We found another ancient split within L2a5 (55 kya), which is centered in southern Africa (Zambia), but also found in eastern Africa (Kenya, Rwanda, Madagascar), suggesting an early Paleolithic southward migration. The L2a5^ sample is from the Kuvale people of Angola, semi-nomadic cattle-herding pastoralists similar to the Bantu Herero, but with significant Khoisan admixture (Mitchell, 2010; Barbieri et al., 2014b). Three new subclades >30 kya (oldest 39 kya) were also added to L2a5, previously a terminal haplogroup, now with mean age of 11 kya. L2a1 is the most ubiquitous haplogroup in sub-Saharan Africa, owing to the recent Bantu migration. Two large Bantu-associated polytomies were expanded: L2a1f (7 kya) increased from 3 to 52 subclades, and L2a1a2 (6 kya) increased from 3 to 37 subclades.

#### L4: Horn of Africa diversity

East Africa harbors some of the most highly distinct and rare African lineages, including L4, L5, L6, and L7. Terminal haplogroup L4b2b1, known from a few samples of the autochthonous Hadza hunter-gatherers in Tanzania, was split at 34 kya by the novel L4b2b2, creating L4b2b1’2. The even lesser known L4b2a2 gained a new 30 kya L4b2a2f. Terminal haplogroup L4a2 (32 kya) now has subclades as old as 26 kya (mean 8 kya).

#### L3: Out of Africa implications

Although L2a1 is the most ubiquitous African subclade (13% Sub-Saharan Africa, compared to 5% for second highest L3e1; Maier et al., 2022), collectively, L3’s lineage diversity dominates Africa (L3: 33%; L2: 23%). In its polytomy, L3 also contains M and N, the only two OOA clades (although M1 and N-descendant U6 are considered African back-migrations). Contrary to the traditional groupings of L3b’f based on [5]15944T[4] and L3c’d based on A13105G!, we found support for L3b’d (T16124C) within a larger L3b’c’d (A13105G!), with [5]15944T[4] occurring in parallel twice, on L3b and L3f, and reducing overall homoplasy. Geographically, this groups L3b (strong West African signal, with signal of eastward migration to Uganda, Kenya, and Madagascar) with L3d (faint West African origin, with strong signals of migration south and eastward to Namibia and Madagascar). The rare L3c lineage also hints at a west-east migration, with roots in Nilo-Saharan western groups (Songhai, Hassaniya, Fulbe in Nigeria and Niger), but spanning east to Sudan, Ethiopia, and Kenya (Nilotic Turkana, Bantu Kikuyu, and Yemenite Jews). Temporally, this west-to-east migration coincides with the OOA event (L3b’c’d: 59 kya, L3b’d: 54 kya).

Except for the major Bantu expansion marker L3e found across Africa and the globe, the remaining L3 haplogroups have epicenters in East Africa (L3a, L3f, L3h, L3i, L3x), or a bimodal east-west distribution (L3k; however, a paucity of samples exists). This latter haplogroup L3k was found to contain a 34 kya split L3k’o, by the novel L3o of unknown geographic origin. Within L3h (common in Nilotic populations of Uganda, Kenya, and Sudan), we found a novel haplogroup L3h1a2a’b at 41 kya. Previously terminal haplogroup L3h1a2b now has subclades (oldest 19 kya, mean 9 kya). L3f (common in eastern Afro-Asiatic populations around Chad, but also southern Bantu populations) gained a new 47 kya haplogroup L3f1’3’4’6’2a. The recent (6 kya) and previously terminal haplogroup L3f1b1a1, associated with southern Bantu expansion around Mozambique, increased from zero to 33 subclades. Ethiopian-centered L3a1 gained a new 30 kya haplogroup L3a1b^^, and its previously terminal Tanzanian subclade L3a1a gained new subclades (oldest 25 kya, mean 16 kya). One of the major Bantu-expansion polytomies L3e2b (i.e., L3e2+16320; 8 kya) increased from 5 to 68 subclades.

None of these L3 lineages directly participated in the OOA event, but as sibling lineages that document Middle Stone Age movement eastward to the Rift Valley, they depict the population background from which OOA clades M and N emerged. Temporally, L3’s crown age is estimated at 65 kya, and its subclades all coalesce between 49–61 kya (M: 52 kya; N: 57 kya), which is consistent with previous OOA estimates (Soares et al., 2009). Additionally, multiple Neolithic aDNA samples in Kenya and Tanzania containing various subclades of the 45 kya L3h1a (Prendergast et al., 2019), suggest a deep continuity of Rift Valley populations, during and after the exodus.

#### M: Out-of-Africa radiation in South Asia

The OOA exodus (50–70 kya) left its legacy in two non-African superclades (M and N descendants of L3) from which all living non-Africans trace their matrilineal history. M is virtually absent from the Levant and Western Eurasia today (although found in 35–47 kya ancient European burials; Hublin et al., 2020; Posth et al., 2016), yet its extreme multifurcation in South and Southeast Asia, and into Melanesia and Australia, trace humanity’s rapid arc across the southern Asian coast, from the Horn of Africa into the Pacific (Macaulay et al., 2005). Haplogroups such as M2 (represented today at high frequencies in Dravidian tribes; e.g., 40% in the Betta Kuruba) represent the massive Indian radiation that occurred immediately post-Africa (Kumar et al., 2008), with only haplogroups M1 and U6 (in R) back-migrating into North and East Africa millennia later from the Levant (Olivieri et al., 2006). Indigenous tribes of the Andaman and Nicobar Islands (subclades of haplogroups M31 and M32) are living descendants of this 50 kya OOA seacombing expansion.

M’s polytomy included 80 deep-rooted lineages in PTv17 (Table **S17**), and Mitotree adds 28 new primary haplogroups. The majority of M lineages have an Indian epicenter: M2, M3, M4, M5, M6, M14 (in part), M18, M30, M33 (in part: everything but M33c), M34, M35, M36, M37, M38, M39, M40, M41, M42 (in part: M42b), M43, M44, M45, M47, M48, M49, M52, M53, M54, M55, M56, M57, M58, M62, M63, M64, M65, M66, M67, M68, M69, M70, M76, M77, M81, which span Dravidian, Austro-Asiatic, Sino-Tibetan, and Indo-European tribes across the subcontinent. New Indian haplogroups include: M78, M86, M97, M99, M100, M101, M104, M105, M106, M107, M115, M117, M118, M119, M120, and M121. Numerous ancient clades (30–50 kya) were added to the M structure: M2a’b’c’d’e, M2b’d’e, M5b^, M5c2^, M6’105, M14b (in previously terminal haplogroup M14), M33b^, M34’115, M36a’b’d’e, M36a’d’e, M36a’e, M38’121, M39’97^, M39’97, M42b2’3 (in previously terminal haplogroup M42b2), M44^, M52b1’3, M58a’b (in previously terminal haplogroup M58), M63’119’120, M63’119, M66b^, M76^, M117’118, and M118’5c1. M47 (sequences unavailable) and M66 (restructured) are different in Mitotree. M33 (a, b, and d) which are South Asian are no longer united with M33c (East Asian and Ashkenazi Jewish in Europe).

Southeast Asia (spanning Myanmar to Indonesia) is the second largest stronghold of M, continuing the southern trajectory toward Oceania, including M13, M17, M19, M20, M21, M22, M24, M26, M46, M50, M51, M59, M60, M61, M71, M72, M73, M74, M75, M79, M80, M91 (Hill et al., 2006; Kutanan et al., 2017), to which we added new primary haplogroups M92, M95, M96, M103, M108, M109, M110, M111, M112, M114, M116. Numerous ancient clades (30–50 kya) were added: M13c’d (in previously terminal haplogroup M13c), M17b, M17d, M17d1’c1a, M20’114, M20^^, M21b’e, M21e, M22a and M22b (both previously terminal haplogroups now with subclades), M26^, M50^, M50b, M59 (previously terminal haplogroup now with subclades), M60b’c, M73b”h, and M75’110. Several M branches extend northward into Northeast Asia (China, Japan, Korea), including M7, M8 (parent of CZ), M9 (parent of E), M10, M11, M12, G (sibling of M12), M33 (in part: M33c excluding novel Ashkenazi M33c9b+8251), and D (in part: excluding Native American branches). Haplogroups C (excluding Native American branches) and Z extend into Siberia. We found two splits >30 kya: M8a’b and M7b1^. Haplogroup E diversified through Island Southeast Asia, becoming common in Indonesia, the Philippines, and amongst Taiwanese aboriginals, and traced the Austronesian expansion along with markers such as M23 (endemic to Madagascar), further discussed below.

#### M: India-Sahul connection

We found deep connections between India and Australia, relics from the rapid coastal OOA expansion. M15, M42a and novel M42c and M42d are endemic to Australian Aboriginals, while M14 spans both India and Australia. The Pleistocene Sahul continent (during which Australia was contiguous with Papua New Guinea; PNG) showed a separation of endemic PNG lineages M25, M27, M28, M29, and Q (sibling of M29). We discovered that novel Indian M86 splits Australian M25 (M25’86) at 46 kya, and Indian M5 appears to form a 50 kya grouping with Melanesian M28 (however, this is tentative based on G16129A! and we do not yet unite these branches under M). Within Sahulian haplogroups, we also found novel ancient (30–50 kya) structure: M15^, M28^, M42a (previously terminal haplogroup with ancient subclades such as M42a2), M42c, and M42d. Aside from Sahul-connections, many backbone M groups found novel groupings older than 30 kya, including India-specific (M53’100^, M40’62, M78^, M52’58), East Asian-specific (M26’73, M19’109, M46’79), and crossovers between the two (M2’46, M77”104, M77’92’95, M13’61, M13”108, M11’81, M55”108). Homoplasy is rampant amongst M’s short-stemmed children, where single-mutation groupings are difficult to distinguish from convergence; several of these novel structures should be treated as provisional pending corroboration from additional samples.

#### N: Out-of-Africa radiation in West Eurasia

The other megaclade to leave Africa (N) radiated most extensively in East Asia, yet several of its branches (particularly the mega-diverse R) dominate 80-90% of Europe, the Near East, and over 60% of Central Asia today. Haplogroup H comprises about 40% of West Europeans, U (including East European K) covers another 25% from early hunter-gatherer, while Near Eastern JT is third most frequent at approximately 20%. This crossroads between west and east revealed a previously unknown split early in the history of this OOA clade: Two ancient samples from the 7 kya Neolithic Sahara (Vai et al., 2019; novel N25), along with existing Southeast Asian haplogroup N21 frequent in aboriginal Malays (Hill et al., 2006), add a confident split of N (N+8701) by lacking the signature G8701A. Such a three-way divide (Europe, E Asia, and N Africa) dated to 57 kya coincides perfectly with hyper-arid conditions alleviating, underscoring the Gulf Oasis of the Arabian Peninsula as a hypothesized staging ground for expansion (Fernandes et al., 2012).

The eastward trajectory of N contains the majority of basal N haplogroups (Table **S18**): N5 (India), N7 (Semang Negritos in Malaysia), N8 (China, Korea, Japan), N9 (East Asia and Siberia), Y (child of N9, carried by indigenous groups in the Philippines), N10 and N11 (China), N21 and N22 (pre-Austronesian groups in Southeast Asia), A (East Asia and Siberia, except for Native American A2). We add two new primary haplogroups (N23, N28) and several splits and novel groupings between 30–50 kya: N7’28, N10a’c’d, and N8’23. Further into Oceania, several aboriginal Australian haplogroups exist in N: N13, N14 (sequences unavailable), S, and O, to which we add Australian N27, N31, N32, and PNG haplogroup N29. N13”31 is found to be a four-way split of N13, N27, N29, and N31 from 35 kya, which itself forms a novel split with N32 (N13”32) 38 kya. S gains three clades 30–50 kya (S1’7, S1b’c, S2e; Fig. **6**), and Haplogroup O is also given its second subclade O2 (Gandini et al., 2025).

Basal N lineages in Western Eurasia and the Near East are typically thought to include N1 (and I, child of N1a1b), N2 (especially its child W), N3, and X2/X4 (Native American X2a excluded). We introduce two remarkably ancient novel N* lineages from 34–46 kya samples, some of the oldest Paleolithic humans ever found: 50 kya haplogroup N26 from remains in modern-day Germany (n=4) and Czechia (n=1; Mylopotamitaki et al., 2024; Sümer et al., 2025), and 48 kya haplogroup N6, from remains in Bacho Kiro Cave in Bulgaria (n=2), and also disparate Mongolia (n=1; Hublin et al., 2020; Massilani et al., 2020). The Czech woman’s (Zlatý Kůň) autosomal profile shows her lineage is basal to all known Eurasians (Vallini et al., 2022). Four more samples, including three German and one Neanderthal-admixed Romanian (Oase1; Fu et al., 2015), support N3’10’26: an ancestral grouping that includes European N26, Near East N3, and East Asian N10. These now extinct lineages were equally distinct from modern East and West Eurasians (Sümer et al., 2025), reinforcing the N+8701 pattern of basal OOA N* lineages sharing western geographic epicenter.

#### R: World dominance and the final OOA bifurcation

Despite the nesting of N(R), these two radiations are probably a single population event reflecting a post-bottleneck diffusion across Eurasia, with R’s stem mutations T16223C T12705C reflecting preexisting OOA diversity that accumulated over a short time frame (estimated <500 years). Although R’s dominance today exists in Western Eurasia (H, V, U, and JT), it mirrors M and N with its trail of R* haplogroups that are most diverse across South and East Asia (Table **S19**). Interestingly, we found early evidence of R’s westward migration into Europe with a 46 kya sample (from the same Bulgarian cave as the N6 sample) placed directly on R* without any PVs (Hublin et al., 2020). On the opposite (eastward) trajectory, we resurrected Asian haplogroup B (which only existed in PhyloTree v1–4) based on shared [2]8289CCCCCTCTA[1] and T16189C!, and supported by the 40 kya Tianyuan Man in Beijing, relative of ancestral Asians and Native Americans (Fu et al., 2013a). Between the two, 45 kya Ust’Ishim from Western Siberia is R* for mtDNA and just below K2 for Y-DNA, meaning that in both haploid inheritance systems he predates the moment of diversification between several later West Eurasian, East Eurasian, Native American, and Oceanic lineages (Fu et al., 2014).

The southern coastal arc hypothesis is strengthened by the phylogeographic similarity between M, N, and R migrations. For example, both M and R independently left “bookend” lineages along their course: M31/M32 (Andaman Islands, midpoint relics), R12/R14 (Australia/PNG, endpoint relics), each extremely rare today. Similarly, all three also had at least one major lineage diversification in Australia/PNG: M42a/c/d and Q (M), S (N), and P (R), with P having the most extant subclades (PTv17: n=28; Mitotree: n=306) of any Sahul lineage today. We significantly expand haplogroup P by adding four new subclades (P12, P19–21) to the traditional P1–P10, with many novel splits/haplogroups up to 55 kya: P1’19’20’21, P1a”j, P1c, P2 (first subclades introduced), P3’10, P3b1^, P3b2, P3b2’3, P5 (first subclades introduced), P6b (in previously terminal P6), P7a (in previously terminal P7), P8’12, P8^, P9a (first subclades introduced), P12, P12a, and P19’20’21. Colonization of the Sahul (Australia and PNG) may have occurred across distinct northern (Sulawesi and Moluccas toward modern PNG) and southern (Lesser Sundas and Timor toward Australia) routes based on the distinctly bimodal distribution of Oceanian M/N/R today (Gandini et al., 2025).

Even the diversity of R* haplogroups in India (R1, R2, R5, R6, R7, R8, R30, R31, R32; with newly added R33, R35, R34, R37, R38, R40) shows evidence of an early European connection. The same cave in Ranis, Germany containing the ancient N26 samples also hosted a 45 kya individual (Mylopotamitaki et al., 2024) who supports R5’35, a split between two solidly Indian haplogroups. The aDNA sample fits perfectly at the crux of those lineages, whereas several modern-day Taiwanese individuals are R5’35 with multiple PVs. We found several novel clades between 30–50 kya: R6d, R6’7, R7a’b’i’j, R8’40, R30a1^, R32”1a, R32”38, R35^^, R35^^^. East Asia is known to have R haplogroups R9, F (child of R9), R11, R21, R22, R23, R24, and B (except Native American B2), we also describe multiple new haplogroups in this age range: R9b’c, R21’39^, R22’1b, R22c’d’f, R22b (under previously terminal R22), and one spanning both India and Southeast Asia (R24’34). We also add the oldest described B haplogroup (B8: 47 kya) to the existing 50 kya B (B4, B5, B6), known from just a few modern Chinese individuals. Other novel clades >30 kya from East Asia include B4l, and F1’4.

#### U: The Paleolithic European flagship

Cooling conditions and abundant megafauna across Europe’s steppe-tundra zone brought the first European colonists more than 45 kya, during the Initial Upper Paleolithic (IUP; extinct haplogroups N6, N26). Ghost populations now, these haplogroups were soon replaced by the Gravettian Upper Paleolithic culture 33 kya, erasing N* signal entirely and introducing basal U* lineages, particularly U4 and U5, and this Gravettian–Mesolithic continuum eventually shifted into the so-called Western Hunter Gatherers (WHG) in post-LGM Mesolithic populations (∼14 kya onward). Neolithic farmers expanded from the fertile crescent 9–11 kya with domestication and technologies that largely wiped out WHG U* signal, replacing with N1a (early on but disappeared), T2, K, J1c, H, and W. Finally, the bronze-age Yamnaya (themselves a mixture of partly Eastern Hunter Gatherers with U*), reintroduced U4 and U5a, but also elevated T1a and I.

Today, haplogroup U is distributed in and around Western Eurasia: Europe (U4, U5, U2e, U8 and its main U8b descendant K, novel U11), Anatolia and Near East (U1, U2d, U3, novel U10), Arabian Peninsula (U9), South Asia (U2a/b/c, U7), North Africa (U6). We introduce the following splits or novel clades 30–50 kya: U2c’g, U2a2’5’6’7^, U2c2, U2c1d’h, U4^, U8b^, U9’10, and U4’9’10’11. One ancient U* sample (only one PV) gives population context to the founders of haplogroup U (50 kya): 37 kya Buran-Kaya III from Crimea, and likely progenitor of the Gravettian people who followed (Bennett et al., 2023). The now-extinct Ancient North Eurasians (ANE) in Siberia, who contributed to Native American ancestry, represented several offshoots of extant U haplogroups: 24 kya Mal’ta boy with U* (10 PVs) from Lake Baikal (Raghavan et al., 2014), and 32 kya Yana1/Yana2 with U2”11* (6–7 PVs) from the Sakha Republic near Beringia, a haplogroup ancestral to all U except U1, U5, and U6. We found similar aDNA representation for 43 kya U5* (29 kya samples with no PVs in Austria and Moravia; Fu, Mittnik, et al., 2013; Teschler-Nicola et al., 2020), 46 kya U8* (37 kya sample with 4 PVs in Bacho Kiro Cave in Bulgaria; Hublin et al., 2020), and a 44 kya split of U8b (31 kya sample with 3 PVs in Moravia; Fu, Mittnik, et al., 2013). We introduce a novel 30 kya haplogroup U11 solely composed of ancient (11–23 kya) Italian, French, and Iberian samples which splits U4’9’10, a possible relic of the LGM and the ending Pleistocene Epoch (Fig. **6**; Posth et al., 2016; Higgins et al., 2024; Modi et al., 2020, 2021; Villalba-Mouco et al., 2023).

#### Massive polytomy increases

Hard polytomies are common within mtDNA due to the limited resolution and rapid population explosions at various prehistoric moments, particularly the Neolithic and Bronze Age revolutions. One measure of phylogenetic expansion is how many immediate children were gained in such polytomies. The following haplogroups contain >100 Mitotree children, and either 50 new children or 10× more children in Mitotree (n_2_) than PTv17 (n_1_): H (n_1_=86; n_2_=358), H1 (n_1_=65; n_2_=346), T2b (n_1_=31; n_2_=265), H3 (n_1_=38; n_2_=195), T1a1 (n_1_=17; n_2_=187), V (n_1_=27; n_2_=177), A2 (n_1_=36, ignoring C64T; n_2_=175), B2 (n_1_=25; n_2_=132), K1a (n_1_=19; n_2_=129), H5a1 (n_1_=14; n_2_=125), H1c (n_1_=22; n_2_=116), H6a1a (n_1_=10; n_2_=116), U5a1a1 (i.e., U5a1a1+16192; n_1_=6; n_2_=114), J1c2 (n_1_=20; n_2_=110), and H2a2a1 (n_1_=8; n_2_=108) (Table **S15**).

#### Austronesian, Polynesian, and Malagasy updates

Austronesian-speaking peoples (Austronesians) emerged as a seafaring and agricultural culture in Taiwan, where aboriginals today (Ami and Yami) carry mainly five matrilines: B (B4a, B4c, B5a), E, F (F1a, F3b), M7 (M7b3, M7c1c), and R9 (Soares et al., 2016). The Philippines was subsequently settled by Austronesians 5 kya with a reduced diversity of haplogroups B4a1a, B4b1, B4c1b, E1a1a, M7b3, and M7c3c, and by 3.5–4 kya people speaking Malayo-Polynesian languages spread across Island Southeast Asia (from Sumatra to the Bismarck Archipelago). The Bismarcks became a staging point for Lapita culture (e.g., intricate dentate-stamped pottery; Chambers and Edinur, 2020) by 3.2 kya, where incoming Austronesian haplogroups (e.g., B4a) mixed with New Guinean lineages (M, P, Q). Here, the Polynesian motif B4a1a1a likely emerged, and rose to 70–95% in far-reaching populations, spanning Fiji, Hawaii, Easter Island, and New Zealand by 1300 CE (Duggan et al., 2014). Western Austronesians in Borneo sailed westward across the Indian Ocean and eventually mixed with Bantu people in Madagascar, forging the Malagasy population of East Africa 500–800 CE.

We drastically expand subclade counts for each of the ancestral steps of this journey: B4a1a (Taiwan and Philippines; n_1_=7; n_2_=26), B4a1a1 (Near Oceania, Bismarcks; n_1_=31; n_2_=77), B4a1a1a (Bismarcks, Polynesia; n_1_=23; n_2_=53). Our TMRCAs also compress the Polynesian pre-motif B4a1a1 from a previous estimate of 6.8 (95% CI: 5–8.7) kya (Soares et al., 2011) to 4.5 (95% CI: 4–5.2) kya, more consistent with known chronology of Lapita cultural staging in the Bismarcks, and bolstering the “slow boat to Melanesia” hypothesis (Oppenheimer and Richards, 2001; Kayser, 2010; Chambers and Edinur, 2015). Although M23 and M32c are known endemics to Madagascar (Capredon et al., 2013; Pierron et al., 2017), we restructure M23 to include a split M23^ (lacking C10142T G12618A), which now forms a 20 kya polytomy with novel Vietnamese and Cambodian clades M96 and M111 under M23’96’111. The Malagasy motif (B4a1a1b; sister to the Polynesian motif) grew from zero to 46 subclades, and likewise M23 grew from zero to 17 subclades, and M32c grew from zero to 2 subclades, with the downstream polytomy M32c2a”j containing 11 subclades. The recent mean TMRCA estimates of these three Malagasy clades (3.5–4.5 kya) are inconsistent with the settlement of Madagascar by hypothesized pre-Bantu/pre-Austronesian “Vazimba” (Ricaut et al., 2009).

#### Isolates from OOA (Andaman Islands)

Indigenous tribes of the Andaman and Nicobar Islands have been semi-isolated from other Eurasians since humans left Africa 50–70 kya (Thangaraj et al., 2005). The Onge carry haplogroup M31a1b (we introduce two new subclades) and the haplotype M32a* with A8108G (previously called M32a1, but single haplotypes cannot be haplogroups in Mitotree; Endicott et al., 2006), to which we add one new subclade M32a2. The Jarawa, who share close cultural and linguistic kinship with Onge, and initiated contact with outsiders in 1998, share M31a1b, M31a1b* (a convergent A8108G haplotype, previously called M31a1b1) and M32a1. The Great Andamanese carry M31a1*, M31a1a, M32*, and M32a* (with only ∼50 individuals, it is the most endangered lineage). Finally, the Shompen (Nicobar Islands) carry B5a and R22 (previously “R12”), which have more overlap with mainland populations (Trivedi et al., 2006). Partial mitogenomes in that study make specific comparisons impossible, but R22 increased from zero to 24 subclades, and B5a increased from 25 to 154 subclades. Interestingly, two rare mutations (A8108G and 2156.1A) independently occur in M31 and M32; 2156.1A was once thought to unite the two groups (Barik et al., 2008), but at least seven other mutations now confidently separate them.

#### Native American lineages

The first wave of Amerindian dispersal into the New World consisted of haplogroups A2, B2, C1 (C1b, C1c, C1d), and D (D1, D4h3a) along the Pacific coast, followed by a likely second wave of C4c and X2a interiorly (Tamm et al., 2007; Achilli et al., 2008, 2013; Perego et al., 2009). This reportedly 15–18 kya initial wave fits well with our mean TMRCA estimates, which span 16.9–18.5 kya, followed by a younger wave of C4c (15.8 kya) and X2a (10.0 kya). Out of these founding Amerindian lineages, D4h3a has an almost exclusively Pacific distribution (with the notable exception of Anzick Child of Montana from 12.7 kya), contrasting sharply to the X2a (represented by Kennewick Man) and C4c which traced the interior ice-free corridor east of the Rocky Mountains when it eventually became habitable. B2a radiated out of B2 inside North America 11–13 kya, later following the spread of Uto-Aztecan maize farmers (Yuman, Pima, Papago) from Mesoamerica into the Southwest 4 kya. Paleo-Eskimo and Na-Dene speakers were the third push 3–7 kya, from Alaska into circumpolar Canada and Greenland, carrying mainly A2a, but also D2a (Na-Dene speakers such as Tlingit), D2b (Aleut), D2c/D3 (Eskimo), and D4b1a2a1a2 (Birnirk/Thule Inuit ancestors). The Southern Athabaskan subgroup of Na-Dene speakers (Apache and Navajo) was part of this mixture, injecting A2a (A2a4/A2a5) and B2a from Alaska into the Southwest by 1300–1500 CE. The Thule expansion from Beringia to Greenland in 1000 CE was the final push and brought Neo-Eskimos (carrying similar haplogroups as Paleo-Eskimos) into the Arctic, where they became modern Inuit-Yupik.

We found a Beringian split of B2 (B2^; lacking A3547G) based on an 11.5 kya sample buried in central Alaska (Tackney et al., 2015), and a split of D1 (D1^; lacking C2092T) based on a modern Brazilian sample (Avila et al., 2019). Parts of C1b and C1d were split into a sibling haplogroup C1h (Ecuador, Mexico), and some C1* samples were grouped into C1k (Brazil and Paraguay). Within the large Pan-American A2, we increase the polytomy from 36 to 175 immediate children—ignoring the unstable A2-(C64T) which we found to be somewhat heteroplasmic. Hundreds of lineages that were previously buried as singletons under “A2” with no resolved relationships now occupy named branches, including new combinations of old branches such as A2ac’ai (G16213A), based on Colombian/Ecuadorian (A2ac) and A2ai (Mexican). Novel ancient-modern pairings were discovered such as A2bw’hr (C13989T): consisting of modern A2bw, and A2hr only known from ancient Mayans in Belize/Yucatán (Verdugo et al., 2020; Barquera et al., 2024). Similarly, the following haplogroups expanded their immediate children counts substantially in Mitotree (n_2_) compared to PTv17 (n_1_): B2 (n_1_=25; n_2_=132), C1b (n_1_=13; n_2_=95), C1c (n_1_=7; n_2_=45), C1d (n_1_=3; n_2_=14), D1 (n_1_=13; n_2_=85), and D4h3a (n_1_=9; n_2_=18), after ignoring weaker temporary branches (e.g., C194T or C152T!!). We placed Paleo-Eskimo samples spanning from Siberia to Greenland, including a late Neolithic Ogonyok warrior in Yakutia, Russia (D3*), and the Greenland Saqqaq sample (D2a1), both from approximately 4 kya (Gilbert et al., 2008; Zeng et al., 2025).

#### Ötzi gains family

A 46-year-old Copper Age man dubbed “Ötzi the Iceman” was discovered in 1991, preserved by glacier in the Eastern Italian Alps, and is the best-preserved natural mummy from this early era (∼3300–3100 BCE). Society has since been fascinated by his extensive tattoos, copper axe, many apparent health conditions, and death by arrow to the back. Ötzi’s matriline was first localized to K using HVR1 data T16224C/T16311C (Handt et al., 1994), predicted as a novel K1* branch using coding data from higher quality intestinal samples (Rollo et al., 2006), then finally placed as a unique branch K1ö, standardized later as K1f (Ermini et al., 2008). After exhaustively filtering and sequencing K1 mitogenomes, Coia et al. (2016) estimated that K1f today must have a frequency of <0.3% with 95% confidence, and without any intermediate haplotypes they concluded that it probably had gone extinct.

Despite this impasse, we discovered an exact K1f match to Ötzi with a modern individual of Algerian Shawiya Berber maternal ancestry. This speaks to previous undersampling of African data, and a Berber connection is broadly consistent with Ötzi’s broader genomic profile: ∼90% Early Neolithic Anatolian-farmer ancestry (Wang et al., 2023). This is the maximum of any Copper Age European, a pattern now shown to characterize most alpine individuals from his territory (Croze et al., 2025). The isolated Ötztal Alps seem to have preserved incoming farmer endogamy from incumbent Western Hunter Gatherer ancestry for thousands of years. Ötzi’s K1f gained a K1f* sibling from 1150 CE medieval Germany (Gretzinger et al., 2025), and a K1f1 subclade from 650 BCE in southwestern Europe (Akbari et al., 2026). We also introduce a split to K1f (K1f^) based on a 10.8 kya Mesolithic hunter-gatherer associated with the Iron Gates region between modern Serbia/Romania (Mathieson et al., 2018). Interestingly, the pre-Neolithic samples show more affinity to Neolithic Anatolian-farmers than to Western Hunter Gatherers, suggesting that Iron Gates hunters and later Anatolian farmers are both offshoots of a common LGM refugial population. Ötzi’s K1f ancestry may represent a pre-existing Balkan Mesolithic K1 substrate that got absorbed into and carried westward by the farming expansion (which carried mostly K1a and K1b). We also expand PTv17’s K1-T16362C (i.e., K1d’e’f) into a new haplogroup K1d”k (i.e., K1d’e’f’i’k), based on the newly discovered K1i (Iron Gates Mesolithic/Neolithic; González-Fortes et al., 2017; Mathieson et al., 2018; Mattila et al., 2023) and K1k (modern Sardinians; Olivieri et al., 2017). Sardinians contain the highest fraction (62.5%) of genome-wide similarity with Neolithic Anatolian-farmers (Fernandes et al., 2020), further associating K1d”k with Near East expansion. Finally, we add K1g (ancient), K1l (ancient/modern), and K1m (ancient) directly to K1, recasting Ötzi’s matriline not as an isolated alpine oddity but as a member of a southeastern-European K1 sub-branch with Mesolithic depth in the Balkans.

#### Sardinian-specific haplogroups

Sardinians are genetic outliers in the European landscape, with 78% of modern Sardinian mtDNAs falling into 89 population-specific haplogroups (Olivieri et al., 2017). The principal Sardinian founder lineage U5b3 (Pala et al., 2009) and its subclade U5b3a1a have gained additional structure in Mitotree, but the more dramatic refinement is the formalization of major Sardinian-specific clades that did not exist in PTv17 at all: K1a2f (Near Eastern Late Glacial; under PTv17 K1a2), U5b1i1a (Western European Late Glacial; under PTv17 U5b1i), N1b1a22 (under PTv17 N1b1a), and the Bell Beaker-era HV104 (under PTv17 HV).

#### Counting the newest Jewish founders

Behar et al. (2008a) identified ∼53 founding maternal lineages across Jewish communities but explicitly marked many as “prospective candidates of clades to be defined properly in the future.” Mitotree formalizes a substantial fraction of these: for example, 42% of Ashkenazi Jews fall into five founding lineages that previously had no subclades, and have been substantially resolved: K1a1b1a (20.6%—54 subclades, ignoring C16223T!), N1b1b1 (5.9%—6 subclades; previously “N1b2”), K1a9 (5.7%—14 subclades), K2a2a1 (5.6%—10 subclades; previously “K2a2”), HV1b2 (4.1%—2 subclades; previously “HV1b”). We also found a 4.3 kya split of the primary Ashkenazi K1a1b1a (2.8 kya) based on a Romanian sample, likely from its non-Jewish European roots in the late Neolithic (Costa et al., 2013). We found a similar 5 kya Romanian split of 2.5 kya K2a2a1 in a private sample.

Aside from the Ashkenazi, several Jewish populations experienced strong founder effects, elevating frequencies of several haplogroups (we focus on those with >20% frequency in at least one haplogroup). Within the Belmonte crypto-Jewish community of Portugal, which experienced centuries of inbreeding under the Inquisition, 93.3% of samples were found to be HV0b, which we were able to pinpoint as novel HV0b2 (A8520G) amongst 10 other named subclades. The Mountain Jews who migrated from Persia to the Caucasus include the Azerbaijan community enriched with 58.6% J2b1 (refined into J2b1f2 by C10223T), and the Georgian community enriched with 58.1% HV1a1a1 (refined into HV1a1a11 by A4257G). Two separate Indian Jewish diasporas exist: the Bene (B’nei) Israel of Mumbai, and Cochin Jews of Kochi in southwest India. The Bene Israel have 41.2% M39a1 founder ancestry (refined into M39a1c by A9284G), 11.8% M30c1a1 (no refinement), 8.8% R30b2 (misclassified “R30a”; refined into R30b2a9 by T3335C T14000a), and 5.9% H13a2a1 (reclassified into H13a2a15 based on A183G C11151T). The Cochin Jews have 26.7% M5a1 (reclassified into M5a6b1 based on T4373C G10589A), 17.8% M64 (unknown at the time so called “M50”; reclassified into M64b2a based on T5201C G15355A A15196G C16527T), 11.1% R5a1a (refined into R5a1a4a by G5147A), and 11.1% U1b (reclassified into U1a2f1 based on G8790A G11440A C16222T). The Libyan Jewish community have 39.8% X2e1a1a (refined into X2e1a1a1 by T13789C), and H30 (refined into H30b* by A8638G A14241G, however due to restructuring in Mitotree this becomes H579). Finally, the Silk Road facilitated cultural diffusion and intermarriage between Ashkenazi Jews and northwest Chinese merchants during the Tang Dynasty 1.4 kya, resulting in East Asian haplogroup M33c being introduced into Europe (Tian et al., 2015). We refine this rare introgressed lineage to M33c9b, to standardize this additional Jewish-Chinese diaspora.

#### Saami sampling improved

Saami populations are characterized by high consolidation into two maternal lineages: U5b1b1 (the ’Saami motif,’ defined by 16144/16189/16270) and V, together encompassing ∼90% of Saami mtDNAs (Tambets et al., 2004). Both lineages trace to the Franco-Cantabrian glacial refuge and post-LGM dispersal; U5b1b is shared with Moroccan Berbers at a coalescence of 10 kya, one of the most striking long-distance maternal connections in European population genetics (Achilli et al., 2005). Mitotree dramatically resolves the internal structure of both of these lineages: U5b1b1 expands from 17 to 198 named sub-clades and V1a from 5 to 138, reflecting the dense recent diversification expected in a small, long-isolated population with deep founder concentration.

#### Ainu / Jōmon remnant (Hokkaido, Japan)

The Ainu and their Jōmon predecessors retain maternal lineages that distinguish them as descendants of Paleolithic northeast Asian/Siberian populations. The Funadomari Jōmon individuals from Rebun Island, Hokkaido (Adachi et al., 2009) carried N9b, M7a, and D1a, with N9b strongly predominant; the broader Hokkaido Jōmon sample (Adachi et al., 2011) added D4h2 and G1b to this founder set. Modern Ainu populations show this Jōmon substrate overlaid with two later admixture layers: an Okhotsk-culture Siberian input (G1b, Y1, C5a) and a more recent mainland Japanese input (M7a1, D4, A5). Mitotree resolves the relevant Jōmon lineages substantially: N9b’s previously flat polytomy of N9b1/N9b2/N9b3/N9b4 is reorganized under the compound inner-named clade N9b1’2’3’5’6, joining the original four sub-branches (N9b4 renamed as N9b5) with newly discovered N9b6, and novel N9b8 as the new outgroup; M7a1a doubles from 9 to 16 children including the compound M7a1a4’16, and D4 expands from 18 to 30 immediate children.

### Validation results

Across n=10 simulation replicates modeled on the PTv17 (plus L7) dataset, Mitotree achieved a FNR of 3.5% (95% CI: 3.3–3.7%) and a FPR of 1.0% (95% CI: 0.9–1.1%). When excluding branches supported exclusively by second-tier markers (which are inherently more volatile and subject to change across pipeline runs) these rates improved to a FNR of 2.3% (95% CI: 2.2–2.4%) and FPR of 0.7% (95% CI: 0.6–0.8%). For comparison, FASTTREE2 produced a FNR of 4.1% (95% CI: 4.1–4.2%) and FPR of 1.0% (95% CI: 1.0–1.1%), which also improved to a FNR of 3.0% (95% CI: 2.9–3.0%) and FPR of 0.7% (95% CI: 0.7–0.7%) when ignoring weaker branches. Mitotree’s methodology thus recovers true branches more reliably than the only other method capable of operating at comparable scales. These simulation-based estimates are consistent with the qualitative PTv17 branch recovery rate (92–96%) reported in Table **S2** and described in “Extinct haplogroups”; this larger (4–8%) discrepancy between PhyloTree and Mitotree reflects genuine biological and methodological differences between the two trees (i.e., topology changes, collapsed branches, and haplogroup renaming) rather than merely pipeline error, and is therefore not directly comparable to the simulation FNR.

## Discussion

### Implications for phylogenetics with 100,000s sequences

The choice of weighted maximum parsimony (WMP) over maximum likelihood (ML) for Mitotree had both practical and theoretical motivations, but the theoretical advantages of WMP on deeply homoplastic sequence data deserve elaboration. Although ML promises statistical consistency, this guarantee assumes both (1) infinite data and (2) correct model specification—the estimated topology and branch lengths asymptotically converge on the correct answer only if the substitution model matches the true evolutionary process. While our dataset is large, mtDNA’s short length (∼16 kb) yields few phylogenetically informative characters, a situation under which WMP appears to have advantages over methods assuming greater uniformity (Goloboff, 2022). Compounding this, human mtDNA violates several key assumptions of standard nucleotide models. Endicott and Ho (2008) observed extreme rate heterogeneity across data partitions, with clear evidence of substitutional saturation, natural selection, and significant heterotachy. ML methods (e.g., the common GTR+Γ model) accommodate rate heterogeneity by fitting characters onto a discrete gamma distribution, which imposes a parametric shape on what is, in human mtDNA, a heavily right-skewed and irregular distribution of homoplasy counts. Mitotree’s empirically derived weights are instead case-by-case reflections of homoplasy observed at each mutation, without any assumption about distributional shape. Thus, WMP offers a meaningful advantage given the true rate distribution (heavy-tailed and irregular) departs substantially from gamma. Our exponential compression formula—the two-step transformation from raw counts to final weights 𝐳—was specifically designed to handle a log-normal count distribution with heavy right skew, which no standard gamma parameterization would capture well. Simulations under models appropriate for discrete characters with heterogeneous rates, including morphological data, showed that implied weighting not only matches ML but clearly outperforms it across a range of evolutionary scenarios (Goloboff et al., 2018).

Heterotachy presents a distinct challenge that neither global weights nor GTR+Γ alone can address, since a character that is hypervariable across the full tree may be phylogenetically reliable within a particular subclade (or vice versa). Mitotree handles this through COMBO markers, which locally boost the weight of characters whose globally-derived weights are insufficient to compete with conflicting signals in specific clades (extending the concept of combining correlated characters proposed by Capella-Gutierrez et al., 2014; Goloboff and De Laet, 2024, to a local rather than global context), and through local exclusions, which suppress markers that are globally informative but unreliable in specific lineages. These mechanisms operate independently of the choice of optimality criterion and represent a purpose-built solution to heterotachy that neither standard ML nor standard WMP provides. Heterotachy appears to be the norm rather than the exception in molecular sequence data: Lopez et al. (2002) demonstrated that 95% of variable positions in mitochondrial cytochrome b are heterotachous across vertebrate lineages, even in a protein whose function is conserved across all groups, and similar patterns have been documented in bacterial and plastid genes (Lockhart et al., 2006) and chloroplast phylogenomics of gymnosperms (Zhong et al., 2011). Any such dataset where globally-derived character weights misrepresent local phylogenetic signal in specific clades could in principle benefit from a COMBO-marker approach. A related mechanism withholds second-tier (hypervariable) markers from the primary tree search, introducing them only after the stable-marker topology is fixed. While these mechanisms effectively address heterotachy in practice, they remain ad hoc solutions rather than a principled global criterion; ideally, future optimality criteria would handle clade-specific rate shifts without requiring such local adjustments.

Mitotree’s novel divide-and-conquer architecture addresses a scale problem that existing frameworks have not solved. Zaharias and Warnow (2022) noted that no divide-and-conquer method (i.e., NJMERGE, TREEMERGE, GTM) had been benchmarked beyond 50,000 taxa, and that the most scalable approach at the time (GTM) approached RAXML-NG in topological accuracy, but could not keep pace with runtimes of FASTTREE2 and its successors (Park et al., 2021; Zaharias and Warnow, 2022). FASTTREE2 and VERYFASTTREE4 themselves produce much lower accuracy when scaled up (Sayyari et al., 2017; Lees et al., 2018; Park et al., 2021; Smirnov and Warnow, 2021). Mitotree’s pipeline operates on approximately 175,000 unique haplotypes in full heuristic mode, 3.5× the previous benchmark, with sufficient accuracy, demonstrating feasibility at a scale that was computationally uncharted in any published phylogenetic framework. The innovation that enables this is also the key methodological distinction from GTM, NJMERGE and TREEMERGE: rather than computing ML subtrees on disjoint partitions and merging them via a guide-tree topology, Mitotree uses a two-phase parsimony approach in which a preliminary constraint analysis defines strongly supported nodes, and recursive WMP searches are then conducted independently within each constraint clade. This strategy is made computationally feasible by the exceptional speed of TNT (Goloboff et al., 2008; Goloboff and Catalano, 2016), which would be intractable with slower parsimony tools. Node support is assessed through a three-pronged metric (BS, RS, MS) that separates character sampling variance (BS), heuristic search variance (RS), and mutational consistency (MS)—distinct uncertainties that bootstrap alone conflates.

This design avoids a known accuracy ceiling of guide-tree merging methods, where the quality of the merged tree is bounded by the quality of the starting topology, a limitation Park et al. (2021) identified as particularly consequential when FASTTREE2 is used as the guide. By contrast, Mitotree’s constraint nodes are defined by BS and RS thresholds rather than inherited from a single guide tree, making the recursive searches genuinely independent and the merged result less susceptible to errors propagated from an initial approximation. The closest predecessor in scale and scope for automated mtDNA phylogenetics is Blanco et al. (2011), who developed a scalable ML pipeline on approximately 3,000 sequences; Mitotree extends this roughly 60-fold in sequence count while replacing ML with empirically calibrated WMP, and adding recursive constraint partitioning, reversal correction, second-tier marker integration, and aDNA deprioritization. This unique combination addresses sources of error that were either absent at smaller scales or simply left unresolved in that earlier framework. Taken together, the theoretical properties of empirical weighting under realistic models of mtDNA evolution, the novel treatment of heterotachy through COMBO markers, and known failures of ML approaches at extreme scales argue that Mitotree’s recursive WMP is not a methodological compromise, but rather an appropriate and well-grounded choice for this class of problem.

### Limitations to Mitotree

Several limitations of our methodology should be clearly acknowledged. Approximately 74% of haplotypes are proprietary FTDNA mitogenomes unavailable in public repositories, meaning the full tree cannot be independently reconstructed from public data alone; Mitotree, its branch annotations, and TMRCA estimates are nevertheless freely accessible. A related consequence of FTDNA’s predominantly European customer base is sampling bias: haplogroups common in Europe such as H, J, T, K, and U are substantially overrepresented, while African L lineages, Oceanic haplogroups such as P, S, and Q, and East and Southeast Asian clades remain undersampled relative to their true global diversity. Undersampling produces longer stems feeding into affected clades, which attract disproportionately large relaxed-clock corrections under LSD’s weighted least squares framework. This can potentially inflate TMRCA estimates for those haplogroups, which should be interpreted with greater caution. Finally, 117 reversal structures (∼0.22% of branches) remain unresolved in the final tree—30 one-step and 87 two-step types—representing cases where no alternative topology provided a better explanation of the data, whether due to genuine reversals or data artifacts.

### Future applications and improvements

Mitotree is designed for periodic full de novo re-estimation rather than incremental placement of new sequences onto a fixed backbone, a philosophical distinction with practical consequences. Placement-only approaches, including USHER-style tools (Turakhia et al., 2021; Ye et al., 2022) and community-curated resources such as mitoLEAF (Huber et al., 2025) and the EMPOP fine-tuning approach (Dür et al., 2021), efficiently expand an existing topology but cannot detect errors in it. The consequences of an unchallenged backbone are not merely theoretical: haplogroup L7, a lineage approximately 100,000 years old, went entirely undetected in PTv17 because its defining samples were misplaced under existing branches and no de novo reconstruction was ever attempted after the original manual curation (Maier et al., 2022). As sequencing data continue to accumulate—particularly from underrepresented populations and ancient samples—topological errors of this kind are likely to persist or multiply in any resource that does not periodically reconstruct the tree from scratch. We regard mitoLEAF as a valuable and complementary resource: its collaborative, community-driven curation model and forensic orientation occupy a distinct niche from Mitotree’s fully computational approach, and its expansion from 5,435 to 6,409 haplogroups reflects meaningful incremental progress within the PTv17 framework. YFull’s MTree represents another actively maintained resource in the genetic genealogy community, with 27,672 haplogroups as of November 2025, but differs fundamentally from Mitotree in relying on manual curation of customer NGS data rather than de novo computational reconstruction, a method that scales less readily and currently yields roughly one-third the haplogroup resolution of Mitotree. These parallel trees (MTree and mitoLEAF) pursue different but compatible goals to Mitotree’s aim of becoming the de facto reference phylogeny against which all other human mtDNA trees are compared, given that Mitotree’s 53,588 haplogroups (and growing) effectively subsume the haplogroup content of any currently published resource.

The tenfold increase in resolution unlocks practical advances across several fields. For genealogical research, finer haplogroup assignment dramatically narrows the geographic and temporal inference available from a single mtDNA test. The trajectory of Ötzi’s lineage K1f illustrates this concretely: previously a mysterious singleton branch with no known living relatives and presumed extinct (Coia et al., 2016), it now groups a Mesolithic Balkan subclade and a living North African representative, transforming an apparent evolutionary dead end into a traceable matrilineal connection spanning continents and millennia. This tendency will continue: formerly terminal haplogroups will resolve into geotemporally meaningful clades across countless lineages as population sampling continues to improve. For population genetics, the systematic reduction in median TMRCA of approximately 2,000 years (mean ∼2,800 years) relative to PTv17 has direct implications for any calibrated ancient DNA study that relied on PhyloTree ages. Inferred dates for events such as Neolithic expansions, Polynesian settlement chronology, or Bronze Age dispersals may require recalibration where they depended on PTv17 terminal haplogroup ages as proxies for population divergence times. For medical genetics, haplogroup-disease associations—spanning mitochondrial disorders, metabolic risk factors, and pharmacogenomic sensitivities—are currently tested at the coarse resolution that PTv17 afforded; Mitotree’s much higher granularity could enable substantially more precise association testing in large biobank datasets such as UK Biobank (Bycroft et al., 2018) and gnomAD (Chen et al., 2024), where mtDNA haplogroup assignments are already routinely computed. Finally, the formalization of population-specific founder haplogroups documented here (e.g., Sardinian, Ashkenazi, Belmonte, and Indian Jewish lineages) has direct bioarcheological relevance, enabling more precise biogeographic inference from mtDNA evidence where population-of-origin is a question of interest.

Mitotree’s validation on simulated phylogenies confirmed the pipeline’s accuracy at tractable scales, with false-negative and false-positive branch recovery rates of 3.5% and 1.0% respectively. This is consistent with qualitative indicators observed between trees: 92–96% recovery of PTv17 branches, depending upon how “equivalent” branches are defined. Several directions remain for future development. Geographic coverage is the most pressing: African, Oceanian, and indigenous populations of South and Southeast Asia remain underrepresented, and Mitotree’s future updates may be substantially more informative for these regions as sequencing costs decrease, and controlled-access resources from initiatives such as AGenDA (Fortes-Lima et al., 2026) become more accessible to the broader research community, including through application-based data sharing with industry. Ancient DNA integration could be further improved in future versions: while aDNA currently contributes to both topology and TMRCA estimation at second priority, incorporating raw variant call statistics (e.g., allele depth, strand bias, base quality) directly into the phylogenetic weighting scheme could enable more accurate placement of damaged or low-coverage samples without relying on explicit post-mortem damage models. Looking further ahead, the reservation of root names AA and AB for Neanderthal and Denisovan archaic haplogroups positions Mitotree as a framework for deep phylogenetics across the genus *Homo* (Fig. **4**); the recent recovery of Denisovan mtDNA from the >146,000-year-old Harbin cranium (Fu et al., 2025) demonstrates that such data will continue to emerge, and Mitotree’s naming architecture is designed to absorb them without disruption.

Two broader contributions of this work merit explicit note. First, the four-type naming system (i.e., primary, inner, variant, and interim names), with explicit rules for backwards compatibility and stability under future topological revision, represents a contribution to phylogenetic nomenclature that extends beyond Mitotree itself. The underlying logic prioritizes continuity with existing literature while accommodating new discoveries without cascading renaming. This parallels the stability principles of formal nomenclatural codes and provides a reusable framework for any large haplotype phylogeny that must balance scientific accuracy with community continuity. Second, Mitotree is immediately deployable through HAPLOGREP v3 (Schönherr et al., 2023) plugins in both RSRS and rCRS formats, making it accessible to the large existing community of researchers who already use HAPLOGREP for haplogroup assignment without any change to their analysis pipelines.

Finally, two sources of noise that are not fully resolved by the pipeline warrant acknowledgment. Nuclear mitochondrial DNA segments (NUMTs) represent a genuine if minor source of noise, as nuclear copies of mtDNA segments can mimic real mitochondrial variants in sequencing data, particularly in low-coverage or short-read datasets (Marshall and Parson, 2021). Our quality filters mitigate but cannot eliminate these in third-party sequences—FTDNA’s high-depth (∼1000×) sequencing renders NUMT interference negligible, but external FASTA data cannot be fully validated without raw genotyping scores. Although we have stringently filtered out third-party sequences for accuracy, residual artifacts may still contribute to anomalous branching in a small number of lineages—mtDNA inclusivity and perfect QC must be balanced. Heteroplasmies (HETs), treated here as half-weight substitutions and excluded from several downstream clades, similarly represent an unresolved approximation, one that is standard in the field but that a future pipeline incorporating full HET-aware models could improve upon.

## Conclusions

Mitotree represents a methodological and empirical advance in human mtDNA phylogenetics, combining recursive weighted maximum parsimony, purpose-built solutions to heterotachy, and a principled naming system to produce a tree of 53,588 haplogroups. Available to researchers through HAPLOGREP v3, this tree is an order of magnitude beyond any previously published resource, from the largest mitogenome dataset ever assembled. The resulting resolution transforms haplogroup assignment from a coarse geographic indicator into a precise genealogical and historical tool. Among the most striking findings, Mitotree recovers L2e’f^, a novel African haplogroup splitting ∼83 kya during the Middle Stone Age, resolves Ötzi’s lineage K1f from an apparently extinct terminal branch into a structured clade with living descendants, and identifies population-specific founder haplogroups across dozens of ethnic communities (e.g., distinct Ashkenazi and Sephardic Jewish subclades within haplogroups previously assigned only at coarse resolution). Mitotree also places hundreds of historically documented individuals into refined lineages, including Holy Roman Emperors Otto I and Henry II (Ringbauer et al., 2026), Tsar Nicholas II (Rogaev et al., 2009), President Abraham Lincoln, and First Lady Dolley Madison (Cavagnino et al., 2024). These contributions span population genetics, medical genetics, forensics, history, and the genomic study of the ancient world. Mitotree will be maintained as a living resource, updated periodically (at least once/year) through full de novo reconstruction as new sequences accumulate, with the aim of becoming the de facto reference phylogeny for human mitochondrial DNA.

## Ethics statement

All experiments in this study were conducted in adherence with the set of ethical principles of the Declaration of Helsinki. Ethical clearance for the study was obtained from the Pearl Institutional Review Board (protocol number: 21-GBYG-101). Informed consent was obtained from all participants.

## Data availability

Mitotree can be publicly accessed in two formats: (1) “classic” branch annotations, including mutations, confidence, and countries (https://discover.familytreedna.com/mtdna/L/classic), and (2) “time” tree with TMRCA information (https://discover.familytreedna.com/mtdna/L/tree). HAPLOGREP v3 plugins for assigning new samples to Mitotree can be found in RSRS format (the preferred unbiased reference) or rCRS format (https://github.com/genebygene/mitotree). Approximately 60,000 sequences used during the current study were previously published (Table S1); the FamilyTreeDNA sequences are not publicly available due to privacy terms, but upon reasonable request to the corresponding author, permission can be sought from participants on a case-by-case basis. Mitotree is freely available for academic (non-commercial, non-derivative) use under a CC-BY-NC-ND license.

## List of Abbreviations

aDNA: Ancient DNA
ASR: Ancestral Sequence Reconstruction
BS: Bootstrap Support
FTDNA: FamilyTreeDNA
HET: Heteroplasmy
HTU: Hypothetical Taxonomic Unit
INDEL: Insertion/Deletion
IUPAC: code Int’l Union of Pure & Applied Chemistry: ambiguity code (e.g., Y = C or T)
MCC: Maximum Clade Credibility
ML: Maximum Likelihood
MP: Maximum Parsimony
MPL: Mean Path Length
MS: Mutational Support
mtDNA: Mitochondrial DNA
NGS: Next-generation Sequencing
NJML: Neighbor Joining Maximum Likelihood
PTv17: PhyloTree version 17
rCRS: Revised Cambridge Reference Sequence
RS: Replicate Support
RSRS: Reconstructed Sapiens Reference Sequence
SNP: Single Nucleotide Polymorphism
TBR: Tree Bisection and Reconnection
TMRCA: Time to Most Recent Common Ancestor
TR: Tandem Repeat
WMP: Weighted Maximum Parsimony

## Acknowledgements

We are grateful to all sample donors for participating in this study, either through one of the many public mtDNA studies or FamilyTreeDNA. We thank the Gene by Gene, Ltd. lab for their help sequencing novel samples, and Doron M. Behar for his encouragement. We are grateful to FamilyTreeDNA for sponsoring the effort, and otherwise, the research did not receive any specific grant from funding agencies in the public, commercial, or not-for-profit sectors. We would like to thank Bennett Greenspan (retired after founding FamilyTreeDNA) for his two decades of dedication to scientific discovery and genealogy, and for his commitment to scientific research.

## Author contributions

P.A.M., G.R., R.J.E., and M.G.V. contributed equally to the conception of the study. P.A.M. designed, programmed, and tested the phylogenetic methodology with conceptual input from G.R., including the divide-and-conquer, recursive HTU, aDNA, reversal correction, branch collapse, and TMRCA strategies. P.A.M. aligned and variant called FASTA data, while G.R. designed a database to store and query it, developed both the NGS-calling pipeline, and haplogroup naming strategy. J.D. developed the front-end website interface. P.A.M., G.R., R.J.E., M.G.V., M.T.S., and A.M.B. helped with data validation. P.A.G. contributed major improvements to the TNT software package to allow partial constraints and speed improvements. P.A.M. wrote the manuscript, and all authors provided important feedback in editing the manuscript.

## Additional information

Supplemental information includes one Excel file containing Supplementary Tables S1–S19, and one PDF file containing Supplementary Figures (all novel splits older than 30 kya).

## References

1. Achilli, A., Perego, U.A., Bravi, C.M., Coble, M.D., Kong, Q.-P., Woodward, S.R., Salas, A., Torroni, A. and Bandelt, H.-J., 2008. The phylogeny of the four pan-American MtDNA haplogroups: Implications for evolutionary and disease studies. PLoS One 3(3), e1764. 10.1371/journal.pone.0001764

2. Achilli, A., Perego, U.A., Lancioni, H., Olivieri, A., Gandini, F., Hooshiar Kashani, B., Battaglia, V., Grugni, V., Angerhofer, N., Rogers, M.P., Herrera, R.J., Woodward, S.R., Labuda, D., Smith, D.G., Cybulski, J.S., Semino, O., Malhi, R.S. and Torroni, A., 2013. Reconciling migration models to the Americas with the variation of North American native mitogenomes. Proc. Natl. Acad. Sci. U. S. A. 110(35), 14308–14313. 10.1073/pnas.1306290110

3. Achilli, A., Rengo, C., Battaglia, V., Pala, M., Olivieri, A., Fornarino, S., Magri, C., Scozzari, R., Babudri, N., Santachiara-Benerecetti, A.S., Bandelt, H.-J., Semino, O. and Torroni, A., 2005. Saami and Berbers—an unexpected mitochondrial DNA link. Am. J. Hum. Genet. 76(5), 883–886. 10.1086/430073

4. Adachi, N., Shinoda, K., Umetsu, K., Kitano, T., Matsumura, H., Fujiyama, R., Sawada, J. and Tanaka, M., 2011. Mitochondrial DNA analysis of Hokkaido Jomon skeletons: Remnants of archaic maternal lineages at the southwestern edge of former Beringia. Am. J. Phys. Anthropol. 146(3), 346–360. 10.1002/ajpa.21561

6. Adachi, N., Shinoda, K., Umetsu, K. and Matsumura, H., 2009. Mitochondrial DNA analysis of Jomon skeletons from the Funadomari site, Hokkaido, and its implication for the origins of Native American. Am. J. Phys. Anthropol. 138(3), 255–265. 10.1002/ajpa.20923

7. Akbari, A., Perry, A., Barton, A.R., Kariminejad, M., Gazal, S., Li, Z., Zeng, Y., Mittnik, A., Patterson, N., Mah, M., Zhou, X., Price, A.L., Lander, E.S., Pinhasi, R., Rohland, N., Mallick, S. and Reich, D., 2026. Ancient DNA reveals pervasive directional selection across West Eurasia. Nature. 10.1038/s41586-026-10358-1

8. Anderson, S., Bankier, A.T., Barrell, B.G., de Bruijn, M.H.L., Coulson, A.R., Drouin, J., Eperon, I.C., Nierlich, D.P., Roe, B.A., Sanger, F., Schreier, P.H., Smith, A.J.H., Staden, R. and Young, I.G., 1981. Sequence and organization of the human mitochondrial genome. Nature 290, 457–465. 10.1038/290457a0

9. Arenas, M. and Posada, D., 2010. The effect of recombination on the reconstruction of ancestral sequences. Genetics 184(4), 1133–1139. 10.1534/genetics.109.113423

10. Attimonelli, M., Accetturo, M., Santamaria, M., Lascaro, D., Scioscia, G., Pappadà, G., Russo, L., Zanchetta, L. and Tommaseo-Ponzetta, M., 2005. HmtDB, a human mitochondrial genomic resource based on variability studies supporting population genetics and biomedical research. BMC Bioinformatics 6(S4), S4. 10.1186/1471-2105-6-S4-S4

11. Avila, E., Graebin, P., Chemale, G., Freitas, J., Kahmann, A. and Alho, C.S., 2019. Full mtDNA genome sequencing of Brazilian admixed populations: A forensic-focused evaluation of a MPS application as an alternative to Sanger sequencing methods. Forensic Sci. Int. Genet. 42, 154–164. 10.1016/j.fsigen.2019.07.004

12. Band, G., Le, Q.S., Clarke, G.M., Kivinen, K., Hubbart, C., Jeffreys, A.E., Rowlands, K., Leffler, E.M., Jallow, M., Conway, D.J., Sisay-Joof, F., Sirugo, G., d’Alessandro, U., Toure, O.B., Thera, M.A., Konate, S., Sissoko, S., Mangano, V.D., Bougouma, E.C., Sirima, S.B., Amenga-Etego, L.N., Ghansah, A.K., Hodgson, A.V.O., Wilson, M.D., Enimil, A., Ansong, D., Evans, J., Ademola, S.A., Apinjoh, T.O., Ndila, C.M., Manjurano, A., Drakeley, C., Reyburn, H., Phu, N.H., Quyen, N.T.N., Thai, C.Q., Hien, T.T., Teo, Y.Y., Manning, L., Laman, M., Michon, P., Karunajeewa, H., Siba, P., Allen, S., Allen, A., Bandelt, H.-J., Quintana-Murci, L., Salas, A. and Macaulay, V., 2002. The fingerprint of phantom mutations in mitochondrial DNA data. Am. J. Hum. Genet. 71(5), 1150–1160. 10.1086/344397

13. Bahlo, M., Davis, T.M.E., Simpson, V., Shelton, J., Spencer, C.C.A., Busby, G.B.J., Kerasidou, A., Drury, E., Stalker, J., Dilthey, A., Mentzer, A.J., McVean, G., Bojang, K.A., Doumbo, O., Modiano, D., Koram, K.A., Agbenyega, T., Amodu, O.K., Achidi, E., Williams, T.N., Marsh, K., Riley, E.M., Molyneux, M., Taylor, T., Dunstan, S.J., Farrar, J., Mueller, I., Rockett, K.A. and Kwiatkowski, D.P., 2019. Insights into malaria susceptibility using genome-wide data on 17,000 individuals from Africa, Asia and Oceania. Nat. Commun. 10(1). 10.1038/s41467-019-13480-z

14. Barbieri, C., Güldemann, T., Naumann, C., Gerlach, L., Berthold, F., Nakagawa, H., Mpoloka, S.W., Stoneking, M. and Pakendorf, B., 2014a. Unraveling the complex maternal history of Southern African Khoisan populations. Am. J. Phys. Anthropol. 153(3), 435–448. 10.1002/ajpa.22441

15. Barbieri, C., Vicente, M., Oliveira, S., Bostoen, K., Rocha, J., Stoneking, M. and Pakendorf, B., 2014b. Migration and interaction in a contact zone: mtDNA variation among Bantu-speakers in Southern Africa. PLoS One 9(6), e99117. 10.1371/journal.pone.0099117

16. Barik, S.S., Sahani, R., Prasad, B.V.R., Endicott, P., Metspalu, M., Sarkar, B.N., Bhattacharya, S., Annapoorna, P.C.H., Sreenath, J., Sun, D., Sanchez, J.J., Ho, S.Y.W., Chandrasekar, A. and Rao, V.R., 2008. Detailed mtDNA genotypes permit a reassessment of the settlement and population structure of the Andaman Islands. Am. J. Phys. Anthropol. 136(1), 19–27. 10.1002/ajpa.20773

17. Barquera, R., Del Castillo-Chávez, O., Nägele, K., Pérez-Ramallo, P., Hernández-Zaragoza, D.I., Szolek, A., Rohrlach, A.B., Librado, P., Childebayeva, A., Bianco, R.A., Penman, B.S., Acuña-Alonzo, V., Lucas, M., Lara-Riegos, J.C., Moo-Mezeta, M.E., Torres-Romero, J.C., Roberts, P., Kohlbacher, O., Warinner, C. and Krause, J., 2024. Ancient genomes reveal insights into ritual life at Chichén Itzá. Nature 630(8018), 912–919. 10.1038/s41586-024-07509-7

18. Batai, K., Babrowski, K.B., Arroyo, J.P., Kusimba, C.M. and Williams, S.R., 2013. Mitochondrial DNA diversity in two ethnic groups in Southeastern Kenya: Perspectives from the northeastern periphery of the Bantu expansion. Am. J. Phys. Anthropol. 150(3), 482–491. 10.1002/ajpa.22227

19. Batini, C., Coia, V., Battaggia, C., Rocha, J., Pilkington, M.M., Spedini, G., Comas, D., Destro-Bisol, G. and Calafell, F., 2007. Phylogeography of the human mitochondrial L1c haplogroup: Genetic signatures of the prehistory of Central Africa. Mol. Phylogenet. Evol. 43(2), 635–644. 10.1016/j.ympev.2006.09.014

20. Batini, C., Lopes, J., Behar, D.M., Calafell, F., Jorde, L.B., Van Der Veen, L., Quintana-Murci, L., Spedini, G., Destro-Bisol, G. and Comas, D., 2011. Insights into the demographic history of African Pygmies from complete mitochondrial genomes. Mol. Biol. Evol. 28(2), 1099–1110. 10.1093/molbev/msq294

21. Behar, D.M., Metspalu, E., Kivisild, T., Rosset, S., Tzur, S., Hadid, Y., Yudkovsky, G., Rosengarten, D., Pereira, L., Amorim, A., Kutuev, I., Gurwitz, D., Bonne-Tamir, B., Villems, R. and Skorecki, K., 2008a. Counting the founders: The matrilineal genetic ancestry of the Jewish Diaspora. PLoS One 3(4), e2062. 10.1371/journal.pone.0002062

22. Behar, D.M., Van Oven, M., Rosset, S., Metspalu, M., Loogväli, E.L., Silva, N.M., Kivisild, T., Torroni, A. and Villems, R., 2012. A “Copernican” reassessment of the human mitochondrial DNA tree from its root. Am. J. Hum. Genet. 90(4), 675–684. 10.1016/j.ajhg.2012.03.002

23. Behar, D.M., Villems, R., Soodyall, H., Blue-Smith, J., Pereira, L., Metspalu, E., Scozzari, R., Makkan, H., Tzur, S., Comas, D., Bertranpetit, J., Quintana-Murci, L., Tyler-Smith, C., Wells, R.S. and Rosset, S., 2008b. The dawn of human matrilineal diversity. Am. J. Hum. Genet. 82(5), 1130–1140. 10.1016/j.ajhg.2008.04.002

24. Bennett, E.A., Parasayan, O., Prat, S., Péan, S., Crépin, L., Yanevich, A., Grange, T. and Geigl, E.-M., 2023. Genome sequences of 36,000–to 37,000-year-old modern humans at Buran-Kaya III in Crimea. Nat. Ecol. Evol. 7(12), 2160–2172. 10.1038/s41559-023-02211-9

25. Bergström, A., McCarthy, S.A., Hui, R., Almarri, M.A., Ayub, Q., Danecek, P., Chen, Y., Felkel, S., Hallast, P., Kamm, J., Blanché, H., Deleuze, J.F., Cann, H., Mallick, S., Reich, D., Sandhu, M.S., Skoglund, P., Scally, A., Xue, Y., Durbin, R. and Tyler-Smith, C., 2020. Insights into human genetic variation and population history from 929 diverse genomes. Science 367(6484). 10.1126/science.aay5012

26. Blanco, R., Mayordomo, E., Montoya, J. and Ruiz-Pesini, E., 2011. Rebooting the human mitochondrial phylogeny: An automated and scalable methodology with expert knowledge. BMC Bioinformatics 12(1), 174. 10.1186/1471-2105-12-174

27. Blome, M.W., Cohen, A.S., Tryon, C.A., Brooks, A.S. and Russell, J., 2012. The environmental context for the origins of modern human diversity: a synthesis of regional variability in African climate 150,000-30,000 years ago. J. Hum. Evol. 62(5), 563–592. 10.1016/j.jhevol.2012.01.011

28. Bouckaert, R., Heled, J., Kuhnert, D., Vaughan, T., Wu, C.-H., Xie, D., Suchard, M.A., Rambaut, A. and Drummond, A.J., 2014. BEAST 2: A software platform for Bayesian evolutionary analysis. PLoS Comput. Biol. 10(4), e1003537. 10.1371/journal.pcbi.1003537

29. Bycroft, C., Freeman, C., Petkova, D., Band, G., Elliott, L.T., Sharp, K., Motyer, A., Vukcevic, D., Delaneau, O., O’Connell, J., Cortes, A., Welsh, S., Young, A., Effingham, M., McVean, G., Leslie, S., Allen, N., Donnelly, P. and Marchini, J., 2018. The UK Biobank resource with deep phenotyping and genomic data. Nature 562(7726), 203–209. 10.1038/s41586-018-0579-z

30. Camin, J.H. and Sokal, R.R., 1965. A method for deducing branching sequences in phylogeny. Evolution (N. Y). 19(3), 311. 10.2307/2406441

31. Cann, R.L., Stoneking, M. and Wilson, A.C., 1987. Mitochondrial DNA and human evolution. Nature 325, 31–36. 10.1038/325031a0

32. Capella-Gutierrez, S., Kauff, F. and Gabaldón, T., 2014. A phylogenomics approach for selecting robust sets of phylogenetic markers. Nucleic Acids Res. 42(7), e54. 10.1093/nar/gku071

33. Capredon, M., Brucato, N., Tonasso, L., Choesmel-Cadamuro, V., Ricaut, F.-X., Razafindrazaka, H., Rakotondrabe, A.B., Ratolojanahary, M.A., Randriamarolaza, L.-P., Champion, B. and Dugoujon, J.-M., 2013. Tracing Arab-Islamic inheritance in Madagascar: Study of the Y-chromosome and mitochondrial DNA in the Antemoro. PLoS One 8(11), e80932. 10.1371/journal.pone.0080932

34. Cavagnino, C., Runfeldt, G., Sager, M., Estes, R., Tillmar, A., Greytak, E.M., Thomas, J.T., Anderson, E., Daniels-Higginbotham, J., Kjelland, K., Sturk-Andreaggi, K., Parsons, T.J., McMahon, T.P. and Marshall, C., 2024. Unearthing who and Y at Harewood Cemetery and inference of George Washington’s Y-chromosomal haplotype. iScience 27(4), 109353. 10.1016/j.isci.2024.109353

35. Chambers, G.K. and Edinur, H.A., 2020. Reconstruction of the Austronesian diaspora in the era of genomics. Hum. Biol. 92(4), 247. 10.13110/humanbiology.92.4.04

36. Chambers, G.K. and Edinur, H.A., 2015. The Austronesian diaspora: A synthetic total evidence model. G. J. Anthropol. Res. 72, 53–65. 10.26686/nzsr.v72i4.8582

37. Chan, E.K.F., Hardie, R.-A., Petersen, D.C., Beeson, K., Bornman, R.M.S., Smith, A.B. and Hayes, V.M., 2015. Revised timeline and distribution of the earliest diverged human maternal lineages in southern Africa. PLoS One 10(3), e0121223. 10.1371/journal.pone.0121223

38. Chen, S., Francioli, L.C., Goodrich, J.K., Collins, R.L., Kanai, M., Wang, Q., Alföldi, J., Watts, N.A., Vittal, C., Gauthier, L.D., Poterba, T., Wilson, M.W., Tarasova, Y., Phu, W., Grant, R., Yohannes, M.T., Koenig, Z., Farjoun, Y., Banks, E., Donnelly, S., Gabriel, S., Gupta, N., Ferriera, S., Tolonen, C., Novod, S., Bergelson, L., Roazen, D., Ruano-Rubio, V., Covarrubias, M., Llanwarne, C., Petrillo, N., Wade, G., Jeandet, T., Munshi, R., Tibbetts, K., Abreu, M., Aguilar Salinas, C.A., Ahmad, T., Albert, C.M., Ardissino, D., Armean, I.M., Atkinson, E.G., Atzmon, G., Barnard, J., Baxter, S.M., Beaugerie, L., Benjamin, E.J., Benjamin, D., Boehnke, M., Bonnycastle, L.L., Bottinger, E.P., Bowden, D.W., Bown, M.J., Brand, H., Brant, S., Brookings, T., Bryant, S., Calvo, S.E., Campos, H., Chambers, J.C., Chan, J.C., Chao, K.R., Chapman, S., Chasman, D.I., Chisholm, R., Cho, J., Chowdhury, R., Chung, M.K., Chung, W.K., Cibulskis, K., Cohen, B., Connolly, K.M., Correa, A., Cummings, B.B., Dabelea, D., Danesh, J., Darbar, D., Darnowsky, P., Denny, J., Duggirala, R., Dupuis, J., Ellinor, P.T., Elosua, R., Emery, J., England, E., Erdmann, J., Esko, T., Evangelista, E., Fatkin, D., Florez, J., Franke, A., Fu, J., Färkkilä, M., Garimella, K., Gentry, J., Getz, G., Glahn, D.C., Glaser, B., Glatt, S.J., Goldstein, D., Gonzalez, C., Groop, L., Gudmundsson, S., Haessly, A., Haiman, C., Hall, I., Hanis, C.L., Harms, M., Hiltunen, M., Holi, M.M., Hultman, C.M., Jalas, C., Kallela, M., Kaplan, D., Kaprio, J., Kathiresan, S., Kenny, E.E., Kim, B.-J., Kim, Y.J., King, D., Kirov, G., Kooner, J., Koskinen, S., Krumholz, H.M., Kugathasan, S., Kwak, S.H., Laakso, M., Lake, N., Langsford, T., Laricchia, K.M., Lehtimäki, T., Lek, M., Lipscomb, E., Loos, R.J.F., Lu, W., Lubitz, S.A., Luna, T.T., Ma, R.C.W., Marcus, G.M., Marrugat, J., Mattila, K.M., McCarroll, S., McCarthy, M.I., McCauley, J.L., McGovern, D., McPherson, R., Meigs, J.B., Melander, O., Metspalu, A., Meyers, D., Minikel, E. V., Mitchell, B.D., Mootha, V.K., Naheed, A., Nazarian, S., Nilsson, P.M., O’Donovan, M.C., Okada, Y., Ongur, D., Orozco, L., Owen, M.J., Palmer, C., Palmer, N.D., Palotie, A., Park, K.S., Pato, C., Pulver, A.E., Rader, D., Rahman, N., Reiner, A., Remes, A.M., Rhodes, D., Rich, S., Rioux, J.D., Ripatti, S., Roden, D.M., Rotter, J.I., Sahakian, N., Saleheen, D., Salomaa, V., Saltzman, A., Samani, N.J., Samocha, K.E., Sanchis-Juan, A., Scharf, J., Schleicher, M., Schunkert, H., Schönherr, S., Seaby, E.G., Shah, S.H., Shand, M., Sharpe, T., Shoemaker, M.B., Shyong, T., Silverman, E.K., Singer-Berk, M., Sklar, P., Smith, J.T., Smith, J.G., Soininen, H., Sokol, H., Son, R.G., Soto, J., Spector, T., Stevens, C., Stitziel, N.O., Sullivan, P.F., Suvisaari, J., Tai, E.S., Taylor, K.D., Teo, Y.Y., Tsuang, M., Tuomi, T., Turner, D., Tusie-Luna, T., Vartiainen, E., Vawter, M., Wang, L., Wang, A., Ware, J.S., Watkins, H., Weersma, R.K., Weisburd, B., Wessman, M., Whiffin, N., Wilson, J.G., Xavier, R.J., O’Donnell-Luria, A., Solomonson, M., Seed, C., Martin, A.R., Talkowski, M.E., Rehm, H.L., Daly, M.J., Tiao, G., Neale, B.M., MacArthur, D.G. and Karczewski, K.J., 2024. A genomic mutational constraint map using variation in 76,156 human genomes. Nature 625(7993), 92–100. 10.1038/s41586-023-06045-0

39. Chen, Y.S., Torroni, A., Excoffier, L., Santachiara-Benerecetti, A.S. and Wallace, D.C., 1995. Analysis of mtDNA variation in African populations reveals the most ancient of all human continent-specific haplogroups. Am. J. Hum. Genet. 57(1), 133–49.

40. Cohen, A.S., Stone, J.R., Beuning, K.R.M., Park, L.E., Reinthal, P.N., Dettman, D., Scholz, C.A., Johnson, T.C., King, J.W., Talbot, M.R., Brown, E.T. and Ivory, S.J., 2007. Ecological consequences of early Late Pleistocene megadroughts in tropical Africa. Proc. Natl. Acad. Sci. U. S. A. 104(42), 16422–16427. 10.1073/pnas.0703873104

41. Coia, V., Cipollini, G., Anagnostou, P., Maixner, F., Battaggia, C., Brisighelli, F., Gómez-Carballa, A., Destro Bisol, G., Salas, A. and Zink, A., 2016. Whole mitochondrial DNA sequencing in Alpine populations and the genetic history of the Neolithic Tyrolean Iceman. Sci. Rep. 6(1), 18932. 10.1038/srep18932

42. Costa, M.D., Pereira, J.B., Pala, M., Fernandes, V., Olivieri, A., Achilli, A., Perego, U.A., Rychkov, S., Naumova, O., Hatina, J., Woodward, S.R., Eng, K.K., MacAulay, V., Carr, M., Soares, P., Pereira, L. and Richards, M.B., 2013. A substantial prehistoric European ancestry amongst Ashkenazi maternal lineages. Nat. Commun. 4, 1–10. 10.1038/ncomms3543

43. Croze, M., Paladin, A., Zingale, S., Alemanno, S., Nicolis, F., Mottes, E., Maixner, F., Pedrotti, A., Günther, T., Zink, A. and Coia, V., 2025. Genomic diversity and structure of prehistoric alpine individuals from the Tyrolean Iceman’s territory. Nat. Commun. 16(1), 6431. 10.1038/s41467-025-61601-8

44. De Maio, N., Kalaghatgi, P., Turakhia, Y., Corbett-Detig, R., Minh, B.Q. and Goldman, N., 2023. Maximum likelihood pandemic-scale phylogenetics. Nat. Genet. 55(5), 746–752. 10.1038/s41588-023-01368-0

45. Duggan, A.T., Evans, B., Friedlaender, F.R., Friedlaender, J.S., Koki, G., Merriwether, D.A., Kayser, M. and Stoneking, M., 2014. Maternal history of Oceania from complete mtDNA genomes: Contrasting ancient diversity with recent homogenization due to the Austronesian expansion. Am. J. Hum. Genet. 94(5), 721–733. 10.1016/j.ajhg.2014.03.014

46. Dür, A., Huber, N. and Parson, W., 2021. Fine-tuning phylogenetic alignment and haplogrouping of mtDNA sequences. Int. J. Mol. Sci. 22(11), 5747. 10.3390/ijms22115747

47. Edgar, R.C., 2004. MUSCLE: A multiple sequence alignment method with reduced time and space complexity. BMC Bioinformatics 5, 1–119. 10.1186/1471-2105-5-113

48. Ehler, E., Novotńy, J., Juras, A., Chylénski, M., Moravčik, O. and Pačes, J., 2019. AmtDB: A database of ancient human mitochondrial genomes. Nucleic Acids Res. 47(D1), D29–D432. 10.1093/nar/gky843

49. Endicott, P. and Ho, S.Y.W., 2008. A Bayesian evaluation of human mitochondrial substitution rates. Am. J. Hum. Genet. 82(4), 895–902. 10.1016/j.ajhg.2008.01.019

50. Endicott, P., Metspalu, M., Stringer, C., Macaulay, V., Cooper, A. and Sanchez, J.J., 2006. Multiplexed SNP typing of ancient DNA clarifies the origin of Andaman mtDNA haplogroups amongst South Asian tribal populations. PLoS One 1(1), e81. 10.1371/journal.pone.0000081

51. Ermini, L., Olivieri, C., Rizzi, E., Corti, G., Bonnal, R., Soares, P., Luciani, S., Marota, I., De Bellis, G., Richards, M.B. and Rollo, F., 2008. Complete mitochondrial genome sequence of the Tyrolean Iceman. Curr. Biol. 18(21), 1687–1693. 10.1016/j.cub.2008.09.028

52. Fernandes, D.M., Mittnik, A., Olalde, I., Lazaridis, I., Cheronet, O., Rohland, N., Mallick, S., Bernardos, R., Broomandkhoshbacht, N., Carlsson, J., Culleton, B.J., Ferry, M., Gamarra, B., Lari, M., Mah, M., Michel, M., Modi, A., Novak, M., Oppenheimer, J., Sirak, K.A., Stewardson, K., Mandl, K., Schattke, C., Özdoğan, K.T., Lucci, M., Gasperetti, G., Candilio, F., Salis, G., Vai, S., Camarós, E., Calò, C., Catalano, G., Cueto, M., Forgia, V., Lozano, M., Marini, E., Micheletti, M., Miccichè, R.M., Palombo, M.R., Ramis, D., Schimmenti, V., Sureda, P., Teira, L., Teschler-Nicola, M., Kennett, D.J., Lalueza-Fox, C., Patterson, N., Sineo, L., Coppa, A., Caramelli, D., Pinhasi, R. and Reich, D., 2020. The spread of steppe and Iranian-related ancestry in the islands of the western Mediterranean. Nat. Ecol. Evol. 4(3), 334–345. 10.1038/s41559-020-1102-0

53. Fernandes, V., Alshamali, F., Alves, M., Costa, M.D., Pereira, J.B., Silva, N.M., Cherni, L., Harich, N., Cerny, V., Soares, P., Richards, M.B. and Pereira, L., 2012. The Arabian cradle: Mitochondrial relicts of the first steps along the Southern route out of Africa. Am. J. Hum. Genet. 90(2), 347–355. 10.1016/j.ajhg.2011.12.010

54. Fu, Q., Cao, P., Dai, Q., Bennett, E.A., Feng, X., Yang, M.A., Ping, W., Pääbo, S. and Ji, Q., 2025. Denisovan mitochondrial DNA from dental calculus of the >146,000-year-old Harbin cranium. Cell 188(15), 3919–3926. 10.1016/j.cell.2025.05.040

55. Fu, Q., Hajdinjak, M., Moldovan, O.T., Constantin, S., Mallick, S., Skoglund, P., Patterson, N., Rohland, N., Lazaridis, I., Nickel, B., Viola, B., Prüfer, K., Meyer, M., Kelso, J., Reich, D. and Pääbo, S., 2015. An early modern human from Romania with a recent Neanderthal ancestor. Nature 524(7564), 216–219. 10.1038/nature14558

56. Fu, Q., Li, H., Moorjani, P., Jay, F., Slepchenko, S.M., Bondarev, A.A., Johnson, P.L.F., Aximu-Petri, A., Prüfer, K., de Filippo, C., Meyer, M., Zwyns, N., Salazar-García, D.C., Kuzmin, Y. V., Keates, S.G., Kosintsev, P.A., Razhev, D.I., Richards, M.P., Peristov, N. V., Lachmann, M., Douka, K., Higham, T.F.G., Slatkin, M., Hublin, J.-J., Reich, D., Kelso, J., Viola, T.B. and Pääbo, S., 2014. Genome sequence of a 45,000-year-old modern human from western Siberia. Nature 514(7523), 445–449. 10.1038/nature13810

58. Fu, Q., Meyer, M., Gao, X., Stenzel, U., Burbano, H.A., Kelso, J. and Pääbo, S., 2013a. DNA analysis of an early modern human from Tianyuan Cave, China. Proc. Natl. Acad. Sci. U. S. A. 110(6), 2223–2227. 10.1073/pnas.1221359110

59. Fu, Q., Mittnik, A., Johnson, P.L.F., Bos, K., Lari, M., Bollongino, R., Sun, C., Giemsch, L., Schmitz, R., Burger, J., Ronchitelli, A.M., Martini, F., Cremonesi, R.G., Svoboda, J., Bauer, P., Caramelli, D., Castellano, S., Reich, D., Pääbo, S. and Krause, J., 2013b. A revised timescale for human evolution based on ancient mitochondrial genomes. Curr. Biol. 23(7), 553–559. 10.1016/j.cub.2013.02.044

60. Gandini, F., Almeida, M., Foody, M.G.B., Nagle, N., Bergström, A., Olivieri, A., Rodrigues, S., Fichera, A., Oteo-Garcia, G., Torroni, A., Achilli, A., Pomat, W., Zainuddin, Z., Eng, K.K., Shoeib, T., Rito, T., Bulbeck, D., O’Connor, S., Bryk, J., Pala, M., Grant, M.J., Edwards, C.J., Oppenheimer, S.J., Mitchell, R.J., Soares, P.A., Farr, H. and Richards, M.B., 2025. Genomic evidence supports the “long chronology” for the peopling of Sahul. Sci. Adv. 11(48), 9493. 10.1126/sciadv.ady9493

61. GATK Team, 2025. Mitochondrial short variant discovery (SNVs + Indels) – GATK Best Practices Workflows. Broad Institute. Available at: https://gatk.broadinstitute.org/hc/en-us/articles/4403870837275-Mitochondrial-short-variant-discovery-SNVs-Indels (accessed 26 May 2026).

62. Gilbert, M.T.P., Kivisild, T., Grønnow, B., Andersen, P.K., Metspalu, E., Reidla, M., Tamm, E., Axelsson, E., Götherström, A., Campos, P.F., Rasmussen, M., Metspalu, M., Higham, T.F.G., Schwenninger, J.-L., Nathan, R., De Hoog, C.-J., Koch, A., Møller, L.N., Andreasen, C., Meldgaard, M., Villems, R., Bendixen, C. and Willerslev, E., 2008. Paleo-Eskimo mtDNA genome reveals matrilineal discontinuity in Greenland. Science. 320(5884), 1787–1789. 10.1126/science.1159750

63. Goloboff, P.A., 2022. Refining phylogenetic analyses: Phylogenetic analysis of morphological data: Volume 2. CRC Press, Boca Raton. 10.1201/9780367823412

64. Goloboff, P.A., 2015. Computer science and parsimony: A reappraisal, with discussion of methods for poorly structured datasets. Cladistics 31(2), 210–225. 10.1111/cla.12082

65. Goloboff, P.A., 1999. Analyzing large data sets in reasonable times: Solutions for composite optima. Cladistics 15(4), 415–428. 10.1006/clad.1999.0122

66. Goloboff, P.A., 1993a. Estimating character weights during tree search. Cladistics 9(1), 83–91. 10.1111/j.1096-0031.1993.tb00209.x

67. Goloboff, P.A., 1993b. Character optimization and calculation of tree lengths. Cladistics 9(4), 433–436. 10.1111/j.1096-0031.1993.tb00236.x

68. Goloboff, P.A. and Catalano, S.A., 2016. TNT version 1.5, including a full implementation of phylogenetic morphometrics. Cladistics 32(3), 221–238. 10.1111/cla.12160

69. Goloboff, P.A. and De Laet, J., 2024. Farewell to the requirement for character independence: Phylogenetic methods to incorporate different types of dependence between characters. Cladistics 40(3), 209–241. 10.1111/cla.12564

70. Goloboff, P.A., Farris, J.S. and Nixon, K.C., 2008. TNT, a free program for phylogenetic analysis. Cladistics 24(5), 774–786. 10.1111/j.1096-0031.2008.00217.x

71. Goloboff, P.A. and Maier, P.A., in review. Phylogenetic tree searches under topological constraints. Cladistics.

72. Goloboff, P.A. and Morales, M.E., 2023. TNT version 1.6, with a graphical interface for MacOS and Linux, including new routines in parallel. Cladistics 39(2), 144–153. 10.1111/cla.12524

73. Goloboff, P.A. and Pol, D., 2007. On divide-and-conquer strategies for parsimony analysis of large data sets: Rec-I-DCM3 versus TNT. Syst. Biol. 56(3), 485–495. 10.1080/10635150701431905

74. Goloboff, P.A., Torres, A. and Arias, J.S., 2018. Weighted parsimony outperforms other methods of phylogenetic inference under models appropriate for morphology. Cladistics 34(4), 407–437. 10.1111/cla.12205

75. Gonder, M.K., Mortensen, H.M., Reed, F.A., De Sousa, A. and Tishkoff, S.A., 2007. Whole-mtDNA genome sequence analysis of ancient African lineages. Mol. Biol. Evol. 24(3), 757–768. 10.1093/molbev/msl209

76. González-Fortes, G., Jones, E.R., Lightfoot, E., Bonsall, C., Lazar, C., Grandal-d’Anglade, A., Garralda, M.D., Drak, L., Siska, V., Simalcsik, A., Boroneanţ, A., Vidal Romaní, J.R., Vaqueiro Rodríguez, M., Arias, P., Pinhasi, R., Manica, A. and Hofreiter, M., 2017. Paleogenomic evidence for multi-generational mixing between Neolithic farmers and Mesolithic hunter-gatherers in the Lower Danube Basin. Curr. Biol. 27(12), 1801–1810.e10. 10.1016/j.cub.2017.05.023

77. Gretzinger, J., Biermann, F., Mager, H., King, B., Zlámalová, D., Traverso, L., Gnecchi Ruscone, G.A., Peltola, S., Salmela, E., Neumann, G.U., Radzeviciute, R., Ingrová, P., Liwoch, R., Wronka, I., Jurić, R., Hyrchała, A., Niezabitowska-Wiśniewska, B., Bartecki, B., Borowska, B., Dzieńkowski, T., Wołoszyn, M., Wojenka, M., Wilczyński, J., Kot, M., Müller, E., Orschiedt, J., Zariņa, G., Onkamo, P., Daim, F., Muhl, A., Schwarz, R., Majer, M., McCormick, M., Květina, J., Vida, T., Geary, P.J., Macháček, J., Šlaus, M., Meller, H., Pohl, W., Hofmanová, Z. and Krause, J., 2025. Ancient DNA connects large-scale migration with the spread of Slavs. Nature 646(8084), 384–393. 10.1038/s41586-025-09437-6

78. Handt, O., Richards, M., Trommsdorff, M., Kilger, C., Simanainen, J., Georgiev, O., Bauer, K., Stone, A., Hedges, R., Schaffner, W., Utermann, G., Sykes, B. and Pääbo, S., 1994. Molecular genetic analyses of the Tyrolean Ice Man. Science. 264(5166), 1775–1778. 10.1126/science.8209259

79. Harris, D.J., 2003. Can you bank on GenBank?. Trends Ecol. Evol. 18(7), 317–319. 10.1016/S0169-5347(03)00128-9

80. Henn, B.M., Hon, L., Macpherson, J.M., Eriksson, N., Saxonov, S., Pe’er, I. and Mountain, J.L., 2012. Cryptic distant relatives are common in both isolated and cosmopolitan genetic samples. PLoS One 7(4). 10.1371/journal.pone.0034267

81. Higgins, O.A., Modi, A., Cannariato, C., Diroma, M.A., Lugli, F., Ricci, S., Zaro, V., Vai, S., Vazzana, A., Romandini, M., Yu, H., Boschin, F., Magnone, L., Rossini, M., Di Domenico, G., Baruffaldi, F., Oxilia, G., Bortolini, E., Dellù, E., Moroni, A., Ronchitelli, A., Talamo, S., Müller, W., Calattini, M., Nava, A., Posth, C., Lari, M., Bondioli, L., Benazzi, S. and Caramelli, D., 2024. Life history and ancestry of the late Upper Palaeolithic infant from Grotta delle Mura, Italy. Nat. Commun. 15(1), 8248. 10.1038/s41467-024-51150-x

82. Hill, C., Soares, P., Mormina, M., Macaulay, V., Meehan, W., Blackburn, J., Clarke, D., Raja, J.M., Ismail, P., Bulbeck, D., Oppenheimer, S. and Richards, M., 2006. Phylogeography and ethnogenesis of aboriginal Southeast Asians. Mol. Biol. Evol. 23(12), 2480–2491. 10.1093/molbev/msl124

83. Huber, N., Hurmer, N., Dür, A. and Parson, W., 2025. mitoLEAF: Mitochondrial DNA Lineage, Evolution, Annotation Framework. NAR Genom. Bioinform. 7(2). 10.1093/nargab/lqaf079

84. Hublin, J.-J., Sirakov, N., Aldeias, V., Bailey, S., Bard, E., Delvigne, V., Endarova, E., Fagault, Y., Fewlass, H., Hajdinjak, M., Kromer, B., Krumov, I., Marreiros, J., Martisius, N.L., Paskulin, L., Sinet-Mathiot, V., Meyer, M., Pääbo, S., Popov, V., Rezek, Z., Sirakova, S., Skinner, M.M., Smith, G.M., Spasov, R., Talamo, S., Tuna, T., Wacker, L., Welker, F., Wilcke, A., Zahariev, N., McPherron, S.P. and Tsanova, T., 2020. Initial Upper Palaeolithic *Homo sapiens* from Bacho Kiro Cave, Bulgaria. Nature 581(7808), 299–302. 10.1038/s41586-020-2259-z

85. Ingman, M., 2006. mtDB: Human Mitochondrial Genome Database, a resource for population genetics and medical sciences. Nucleic Acids Res. 34(90001), D749–D751. 10.1093/nar/gkj010

86. Kayser, M., 2010. The human genetic history of Oceania: Near and remote views of dispersal. Curr. Biol. 20(4), R194–R201. 10.1016/j.cub.2009.12.004

87. Khan, M.A., Elias, I., Sjölund, E., Nylander, K., Guimera, R. V., Schobesberger, R., Schmitzberger, P., Lagergren, J. and Arvestad, L., 2013. Fastphylo: Fast tools for phylogenetics. BMC Bioinformatics 14(1). 10.1186/1471-2105-14-334

88. Kogelnik, A., 1996. MITOMAP: A human mitochondrial genome database. Nucleic Acids Res. 24(1), 177–179. 10.1093/nar/24.1.177

89. Kozlov, A.M., Darriba, D., Flouri, T., Morel, B. and Stamatakis, A., 2019. RAxML-NG: A fast, scalable and user-friendly tool for maximum likelihood phylogenetic inference. Bioinformatics 35(21), 4453–4455. 10.1093/bioinformatics/btz305

90. Kumar, S., Padmanabham, P., Ravuri, R.R., Uttaravalli, K., Koneru, P., Mukherjee, P.A., Das, B., Kotal, M., Xaviour, D., Saheb, S. and Rao, V., 2008. The earliest settlers’ antiquity and evolutionary history of Indian populations: Evidence from M2 mtDNA lineage. BMC Evol. Biol. 8(1), 230. 10.1186/1471-2148-8-230

91. Kutanan, W., Kampuansai, J., Srikummool, M., Kangwanpong, D., Ghirotto, S., Brunelli, A. and Stoneking, M., 2017. Complete mitochondrial genomes of Thai and Lao populations indicate an ancient origin of Austroasiatic groups and demic diffusion in the spread of Tai–Kadai languages. Hum. Genet. 136(1), 85–98. 10.1007/s00439-016-1742-y

92. Lees, J.A., Kendall, M., Parkhill, J., Colijn, C., Bentley, S.D. and Harris, S.R., 2018. Evaluation of phylogenetic reconstruction methods using bacterial whole genomes: A simulation based study. Wellcome Open Res. 3, 33. 10.12688/wellcomeopenres.14265.1

93. Lipson, M., Sawchuk, E.A., Thompson, J.C., Oppenheimer, J., Tryon, C.A., Ranhorn, K.L., de Luna, K.M., Sirak, K.A., Olalde, I., Ambrose, S.H., Arthur, J.W., Arthur, K.J.W., Ayodo, G., Bertacchi, A., Cerezo-Román, J.I., Culleton, B.J., Curtis, M.C., Davis, J., Gidna, A.O., Hanson, A., Kaliba, P., Katongo, M., Kwekason, A., Laird, M.F., Lewis, J., Mabulla, A.Z.P., Mapemba, F., Morris, A., Mudenda, G., Mwafulirwa, R., Mwangomba, D., Ndiema, E., Ogola, C., Schilt, F., Willoughby, P.R., Wright, D.K., Zipkin, A., Pinhasi, R., Kennett, D.J., Manthi, F.K., Rohland, N., Patterson, N., Reich, D. and Prendergast, M.E., 2022. Ancient DNA and deep population structure in sub-Saharan African foragers. Nature 603(7900), 290–296. 10.1038/s41586-022-04430-9

94. Lockhart, P., Novis, P., Milligan, B.G., Riden, J., Rambaut, A. and Larkum, T., 2006. Heterotachy and tree building: A case study with plastids and Eubacteria. Mol. Biol. Evol. 23(1), 40–45. 10.1093/molbev/msj005

95. Lopez, P., Casane, D. and Philippe, H., 2002. Heterotachy, an important process of protein evolution. Mol. Biol. Evol. 19(1), 1–7. 10.1093/oxfordjournals.molbev.a003973

96. Macaulay, V., Hill, C., Achilli, A., Rengo, C., Clarke, D., Meehan, W., Blackburn, J., Semino, O., Scozzari, R., Cruciani, F., Taha, A., Shaari, N.K., Raja, J.M., Ismail, P., Zainuddin, Z., Goodwin, W., Bulbeck, D., Bandelt, H.-J., Oppenheimer, S., Torroni, A. and Richards, M., 2005. Single, rapid coastal settlement of Asia revealed by analysis of complete mitochondrial genomes. Science. 308(5724), 1034–1036. 10.1126/science.1109792

97. Maier, P.A., Hu, R., Runfeldt, G., Giniebra, D. and Frichot, E., 2021. myOrigins 3.0: combining global and local methods for determining population ancestry. Houston, TX.

98. Maier, P.A., Runfeldt, G., Estes, R.J. and Vilar, M.G., 2022. African mitochondrial haplogroup L7: a 100,000-year-old maternal human lineage discovered through reassessment and new sequencing. Sci. Rep. 12(1). 10.1038/s41598-022-13856-0

99. Mallick, S., Li, H., Lipson, M., Mathieson, I., Gymrek, M., Racimo, F., Zhao, M., Chennagiri, N., Nordenfelt, S., Tandon, A., Skoglund, P., Lazaridis, I., Sankararaman, S., Fu, Q., Rohland, N., Renaud, G., Erlich, Y., Willems, T., Gallo, C., Spence, J.P., Song, Y.S., Poletti, G., Balloux, F., Van Driem, G., De Knijff, P., Romero, I.G., Jha, A.R., Behar, D.M., Bravi, C.M., Capelli, C., Hervig, T., Moreno-Estrada, A., Posukh, O.L., Balanovska, E., Balanovsky, O., Karachanak-Yankova, S., Sahakyan, H., Toncheva, D., Yepiskoposyan, L., Tyler-Smith, C., Xue, Y., Abdullah, M.S., Ruiz-Linares, A., Beall, C.M., Di Rienzo, A., Jeong, C., Starikovskaya, E.B., Metspalu, E., Parik, J., Villems, R., Henn, B.M., Hodoglugil, U., Mahley, R., Sajantila, A., Stamatoyannopoulos, G., Wee, J.T.S., Khusainova, R., Khusnutdinova, E., Litvinov, S., Ayodo, G., Comas, D., Hammer, M.F., Kivisild, T., Klitz, W., Winkler, C.A., Labuda, D., Bamshad, M., Jorde, L.B., Tishkoff, S.A., Watkins, W.S., Metspalu, M., Dryomov, S., Sukernik, R., Singh, L., Thangaraj, K., Paäbo, S., Kelso, J., Patterson, N. and Reich, D., 2016. The Simons Genome Diversity Project: 300 genomes from 142 diverse populations. Nature 538(7624), 201–206. 10.1038/nature18964

100. Marshall, C. and Parson, W., 2021. Interpreting NUMTs in forensic genetics: Seeing the forest for the trees. Forensic Sci. Int. Genet. 53, 102497. 10.1016/j.fsigen.2021.102497

101. Massilani, D., Skov, L., Hajdinjak, M., Gunchinsuren, B., Tseveendorj, D., Yi, S., Lee, J., Nagel, S., Nickel, B., Devièse, T., Higham, T., Meyer, M., Kelso, J., Peter, B.M. and Pääbo, S., 2020. Denisovan ancestry and population history of early East Asians. Science. 370(6516), 579–583. 10.1126/science.abc1166

102. Mathieson, I., Alpaslan-Roodenberg, S., Posth, C., Szécsényi-Nagy, A., Rohland, N., Mallick, S., Olalde, I., Broomandkhoshbacht, N., Candilio, F., Cheronet, O., Fernandes, D., Ferry, M., Gamarra, B., Fortes, G.G., Haak, W., Harney, E., Jones, E., Keating, D., Krause-Kyora, B., Kucukkalipci, I., Michel, M., Mittnik, A., Nägele, K., Novak, M., Oppenheimer, J., Patterson, N., Pfrengle, S., Sirak, K., Stewardson, K., Vai, S., Alexandrov, S., Alt, K.W., Andreescu, R., Antonović, D., Ash, A., Atanassova, N., Bacvarov, K., Gusztáv, M.B., Bocherens, H., Bolus, M., Boroneanţ, A., Boyadzhiev, Y., Budnik, A., Burmaz, J., Chohadzhiev, S., Conard, N.J., Cottiaux, R., Čuka, M., Cupillard, C., Drucker, D.G., Elenski, N., Francken, M., Galabova, B., Ganetsovski, G., Gély, B., Hajdu, T., Handzhyiska, V., Harvati, K., Higham, T., Iliev, S., Janković, I., Karavanić, I., Kennett, D.J., Komšo, D., Kozak, A., Labuda, D., Lari, M., Lazar, C., Leppek, M., Leshtakov, K., Vetro, D. Lo, Los, D., Lozanov, I., Malina, M., Martini, F., McSweeney, K., Meller, H., Mentušić, M., Mirea, P., Moiseyev, V., Petrova, V., Douglas Price, T., Simalcsik, A., Sineo, L., Šlaus, M., Slavchev, V., Stanev, P., Starović, A., Szeniczey, T., Talamo, S., Teschler-Nicola, M., Thevenet, C., Valchev, I., Valentin, F., Vasilyev, S., Veljanovska, F., Venelinova, S., Veselovskaya, E., Viola, B., Virag, C., Zaninović, J., Zaüner, S., Stockhammer, P.W., Catalano, G., Krauß, R., Caramelli, D., Zariną, G., Gaydarska, B., Lillie, M., Nikitin, A.G., Potekhina, I., Papathanasiou, A., Borić, D., Bonsall, C., Krause, J., Pinhasi, R. and Reich, D., 2018. The genomic history of southeastern Europe. Nature 555(7695), 197–203. 10.1038/nature25778

103. Mattila, T.M., Svensson, E.M., Juras, A., Günther, T., Kashuba, N., Ala-Hulkko, T., Chyleński, M., McKenna, J., Pospieszny, Ł., Constantinescu, M., Rotea, M., Palincaș, N., Wilk, S., Czerniak, L., Kruk, J., Łapo, J., Makarowicz, P., Potekhina, I., Soficaru, A., Szmyt, M., Szostek, K., Götherström, A., Storå, J., Netea, M.G., Nikitin, A.G., Persson, P., Malmström, H. and Jakobsson, M., 2023. Genetic continuity, isolation, and gene flow in Stone Age Central and Eastern Europe. Commun. Biol. 6(1), 793. 10.1038/s42003-023-05131-3

104. Meyer, M., Fu, Q., Aximu-Petri, A., Glocke, I., Nickel, B., Arsuaga, J.L., Martínez, I., Gracia, A., De Castro, J.M.B., Carbonell, E. and Pääbo, S., 2014. A mitochondrial genome sequence of a hominin from Sima de los Huesos. Nature 505(7483), 403–406. 10.1038/nature12788

105. Minh, B.Q., Schmidt, H.A., Chernomor, O., Schrempf, D., Woodhams, M.D., Von Haeseler, A., Lanfear, R. and Teeling, E., 2020. IQ-TREE 2: New models and efficient methods for phylogenetic inference in the genomic era. Mol. Biol. Evol. 37(5), 1530–1534. 10.1093/molbev/msaa015

106. Mitchell, P., 2010. Genetics and southern African prehistory: An archaeological view. J. Anthropol. Sci. 88, 73–92.

107. Modi, A., Catalano, G., D’Amore, G., Di Marco, S., Lari, M., Sineo, L. and Caramelli, D., 2020. Paleogenetic and morphometric analysis of a Mesolithic individual from Grotta d’Oriente: An oldest genetic legacy for the first modern humans in Sicily. Quat. Sci. Rev. 248, 106603. 10.1016/j.quascirev.2020.106603

108. Modi, A., Vai, S., Posth, C., Vergata, C., Zaro, V., Diroma, M.A., Boschin, F., Capecchi, G., Ricci, S., Ronchitelli, A., Catalano, G., Lauria, G., D’Amore, G., Sineo, L., Caramelli, D. and Lari, M., 2021. More data on ancient human mitogenome variability in Italy: New mitochondrial genome sequences from three Upper Palaeolithic burials. Ann. Hum. Biol. 48(3), 213–222. 10.1080/03014460.2021.1942549

109. Molloy, E.K. and Warnow, T., 2019a. Statistically consistent divide-and-conquer pipelines for phylogeny estimation using NJMerge. Algorithms Mol. Biol. 14(1), 14. 10.1186/s13015-019-0151-x

110. Molloy, E.K. and Warnow, T., 2019b. TreeMerge: A new method for improving the scalability of species tree estimation methods. Bioinformatics 35(14), i417–i426. 10.1093/bioinformatics/btz344

111. Morales, M.E. and Goloboff, P.A., 2023. New TNT routines for parallel computing with MPI. Mol. Phylogenet. Evol. 178, 107643. 10.1016/j.ympev.2022.107643

112. Mylopotamitaki, D., Weiss, M., Fewlass, H., Zavala, E.I., Rougier, H., Sümer, A.P., Hajdinjak, M., Smith, G.M., Ruebens, K., Sinet-Mathiot, V., Pederzani, S., Essel, E., Harking, F.S., Xia, H., Hansen, J., Kirchner, A., Lauer, T., Stahlschmidt, M., Hein, M., Talamo, S., Wacker, L., Meller, H., Dietl, H., Orschiedt, J., Olsen, J. V., Zeberg, H., Prüfer, K., Krause, J., Meyer, M., Welker, F., McPherron, S.P., Schüler, T. and Hublin, J.-J., 2024. Homo sapiens reached the higher latitudes of Europe by 45,000 years ago. Nature 626(7998), 341–346. 10.1038/s41586-023-06923-7

113. Nixon, K., 1993. On outgroups. Cladistics 9(4), 413–426. 10.1006/clad.1993.1028

114. Olivieri, A., Achilli, A., Pala, M., Battaglia, V., Fornarino, S., Al-Zahery, N., Scozzari, R., Cruciani, F., Behar, D.M., Dugoujon, J.M., Coudray, C., Santachiara-Benerecetti, A.S., Semino, O., Bandelt, H.J. and Torroni, A., 2006. The mtDNA legacy of the Levantine early Upper Palaeolithic in Africa. Science. 314(5806), 1767–1770. 10.1126/science.1135566

115. Olivieri, A., Sidore, C., Achilli, A., Angius, A., Posth, C., Furtwängler, A., Brandini, S., Capodiferro, M.R., Gandini, F., Zoledziewska, M., Pitzalis, M., Maschio, A., Busonero, F., Lai, L., Skeates, R., Gradoli, M.G., Beckett, J., Marongiu, M., Mazzarello, V., Marongiu, P., Rubino, S., Rito, T., Macaulay, V., Semino, O., Pala, M., Abecasis, G.R., Schlessinger, D., Conde-Sousa, E., Soares, P., Richards, M.B., Cucca, F. and Torroni, A., 2017. Mitogenome diversity in Sardinians: A genetic window onto an island’s past. Mol. Biol. Evol. 34(5), 1230–1239. 10.1093/molbev/msx082

116. Oppenheimer, S.J. and Richards, M., 2001. Slow boat to Melanesia?. Nature 410(6825), 166–167. 10.1038/35065520

117. Ota, S. and Li, W.-H., 2000. NJML: A hybrid algorithm for the neighbor-joining and maximum-likelihood methods. Mol. Biol. Evol. 17(9), 1401–1409. 10.1093/oxfordjournals.molbev.a026423

118. Pala, M., Achilli, A., Olivieri, A., Kashani, B.H., Perego, U.A., Sanna, D., Metspalu, E., Tambets, K., Tamm, E., Accetturo, M., Carossa, V., Lancioni, H., Panara, F., Zimmermann, B., Huber, G., Al-Zahery, N., Brisighelli, F., Woodward, S.R., Francalacci, P., Parson, W., Salas, A., Behar, D.M., Villems, R., Semino, O., Bandelt, H.-J. and Torroni, A., 2009. Mitochondrial haplogroup U5b3: A distant echo of the epipaleolithic in Italy and the legacy of the early Sardinians. Am. J. Hum. Genet. 84(6), 814–821. 10.1016/j.ajhg.2009.05.004

119. Paradis, E. and Schliep, K., 2019. ape 5.0: An environment for modern phylogenetics and evolutionary analyses in R. Bioinformatics 35(3), 526–528. 10.1093/bioinformatics/bty633

120. Park, M., Zaharias, P. and Warnow, T., 2021. Disjoint tree mergers for large-scale maximum likelihood tree estimation. Algorithms 14(5), 148. 10.3390/a14050148

121. Perego, U.A., Achilli, A., Angerhofer, N., Accetturo, M., Pala, M., Olivieri, A., Kashani, B.H., Ritchie, K.H., Scozzari, R., Kong, Q.-P., Myres, N.M., Salas, A., Semino, O., Bandelt, H.-J., Woodward, S.R. and Torroni, A., 2009. Distinctive Paleo-Indian migration routes from Beringia marked by two rare mtDNA haplogroups. Curr. Biol. 19(1), 1–8. 10.1016/j.cub.2008.11.058

122. Petr, M., Hajdinjak, M., Fu, Q., Essel, E., Rougier, H., Crevecoeur, I., Semal, P., Golovanova, L. V., Doronichev, V.B., Lalueza-Fox, C., de la Rasilla, M., Rosas, A., Shunkov, M. V., Kozlikin, M.B., Derevianko, A.P., Vernot, B., Meyer, M. and Kelso, J., 2020. The evolutionary history of Neanderthal and Denisovan Y chromosomes. Science. 369(6511), 1653–1656. 10.1126/science.abb6460

123. Pierron, D., Heiske, M., Razafindrazaka, H., Rakoto, I., Rabetokotany, N., Ravololomanga, B., Rakotozafy, L.M.-A., Rakotomalala, M.M., Razafiarivony, M., Rasoarifetra, B., Raharijesy, M.A., Razafindralambo, L., Ramilisonina, Fanony, F., Lejamble, S., Thomas, O., Mohamed Abdallah, A., Rocher, C., Arachiche, A., Tonaso, L., Pereda-loth, V., Schiavinato, S., Brucato, N., Ricaut, F.-X., Kusuma, P., Sudoyo, H., Ni, S., Boland, A., Deleuze, J.-F., Beaujard, P., Grange, P., Adelaar, S., Stoneking, M., Rakotoarisoa, J.-A., Radimilahy, C. and Letellier, T., 2017. Genomic landscape of human diversity across Madagascar. Proc. Natl. Acad. Sci. U. S. A. 114(32), E6498–E6506. 10.1073/pnas.1704906114

124. Piñeiro, C. and Pichel, J.C., 2024. Efficient phylogenetic tree inference for massive taxonomic datasets: Harnessing the power of a server to analyze 1 million taxa. Gigascience 13. 10.1093/gigascience/giae055

125. Posth, C., Renaud, G., Mittnik, A., Drucker, D.G., Rougier, H., Cupillard, C., Valentin, F., Thevenet, C., Furtwängler, A., Wißing, C., Francken, M., Malina, M., Bolus, M., Lari, M., Gigli, E., Capecchi, G., Crevecoeur, I., Beauval, C., Flas, D., Germonpré, M., van der Plicht, J., Cottiaux, R., Gély, B., Ronchitelli, A., Wehrberger, K., Grigorescu, D., Svoboda, J., Semal, P., Caramelli, D., Bocherens, H., Harvati, K., Conard, N.J., Haak, W., Powell, A. and Krause, J., 2016. Pleistocene mitochondrial genomes suggest a single major dispersal of non-Africans and a late glacial population turnover in Europe. Curr. Biol. 26(6), 827–833. 10.1016/j.cub.2016.01.037

126. Prendergast, M.E., Lipson, M., Sawchuk, E.A., Olalde, I., Ogola, C.A., Rohland, N., Sirak, K.A., Adamski, N., Bernardos, R., Broomandkhoshbacht, N., Callan, K., Culleton, B.J., Eccles, L., Harper, T.K., Lawson, A.M., Mah, M., Oppenheimer, J., Stewardson, K., Zalzala, F., Ambrose, S.H., Ayodo, G., Gates, H.L., Gidna, A.O., Katongo, M., Kwekason, A., Mabulla, A.Z.P., Mudenda, G.S., Ndiema, E.K., Nelson, C., Robertshaw, P., Kennett, D.J., Manthi, F.K. and Reich, D., 2019. Ancient DNA reveals a multistep spread of the first herders into sub-Saharan Africa. Science. 365(6448). 10.1126/science.aaw6275

127. Price, M.N., Dehal, P.S. and Arkin, A.P., 2010. FastTree 2 – Approximately maximum-likelihood trees for large alignments. PLoS One 5(3), e9490. 10.1371/journal.pone.0009490

128. Quintana-Murci, L., Quach, H., Harmant, C., Luca, F., Massonnet, B., Patin, E., Sica, L., Mouguiama-Daouda, P., Comas, D., Tzur, S., Balanovsky, O., Kidd, K.K., Kidd, J.R., Van Der Veen, L., Hombert, J.M., Gessain, A., Verdu, P., Froment, A., Bahuchet, S., Heyer, E., Dausset, J., Salas, A. and Behar, D.M., 2008. Maternal traces of deep common ancestry and asymmetric gene flow between Pygmy hunter-gatherers and Bantu-speaking farmers. Proc. Natl. Acad. Sci. U. S. A. 105(5), 1596–1601. 10.1073/pnas.0711467105

129. R Core Team, 2026. R: A language and environment for statistical computing.

130. Raghavan, M., Skoglund, P., Graf, K.E., Metspalu, M., Albrechtsen, A., Moltke, I., Rasmussen, S., Stafford Jr, T.W., Orlando, L., Metspalu, E., Karmin, M., Tambets, K., Rootsi, S., Mägi, R., Campos, P.F., Balanovska, E., Balanovsky, O., Khusnutdinova, E., Litvinov, S., Osipova, L.P., Fedorova, S.A., Voevoda, M.I., DeGiorgio, M., Sicheritz-Ponten, T., Brunak, S., Demeshchenko, S., Kivisild, T., Villems, R., Nielsen, R., Jakobsson, M. and Willerslev, E., 2014. Upper Palaeolithic Siberian genome reveals dual ancestry of Native Americans. Nature 505(7481), 87–91. 10.1038/nature12736

131. Ramsay, M., Etheredge, H., Tluway, F., D’Amato, M.E., Chikwambi, Z., Hamdi, Y., Alhudiri, I., Fakim, Y., Ahmad, K.M., Belguith, N., Bentley, D., Boujemaa, M., Calumbuana, N., Chaouch, M., Charfeddine, C., Chinien, G., Dukuze, N., Eljilani, M., Elzagheid, A., Ferraz, N., Ghoorah, A., Goorah, S., Gribaa, M., Guidara, S., Guirat, M., Hazelhurst, S., Jallul, M., Kasu, M., Kharrat, N., Khumalo, U., Kingsbury, Z., Kisiangani, I., Lopes-Cendes, I., Lukusa, P., Makay, P., Makulo, J., Mubungu, G., Muhinda, C., Mukhongo, D.M., Murwira, A., Mustafa, A., Ndinkabandi, J., Ngole, M., Nlandu, Y., Nyathi, M., Pereira, L., Rejeb, I., Santos, L.L., Sengupta, D., Shebani, A., Smyth, N., Souissi, A., Trabelsi, M., Rebai, A., Chimpolo, M.M., Lumaka, A., Masimirembwa, C., Mohamed, S.F., Mulder, N., Mutesa, L., Hanchard, N.A. and Choudhury, A., 2026. Enriching African genome representation through the AGenDA project. Nature 649(8097), 565–573. 10.1038/s41586-025-09935-7

132. Rannala, B., 2025. Recombination and phylogenetic inference. Evolutionary Journal of the Linnean Society 4(1). 10.1093/evolinnean/kzaf016

133. Revell, L.J., 2024. phytools 2.0: An updated R ecosystem for phylogenetic comparative methods (and other things). PeerJ 12. 10.7717/peerj.16505

134. Ricaut, F.-X., Razafindrazaka, H., Cox, M.P., Dugoujon, J.-M., Guitard, E., Sambo, C., Mormina, M., Mirazon-Lahr, M., Ludes, B. and Crubézy, E., 2009. A new deep branch of Eurasian mtDNA macrohaplogroup M reveals additional complexity regarding the settlement of Madagascar. BMC Genomics 10(1), 605. 10.1186/1471-2164-10-605

135. Ringbauer, H., Wozniak, T., Feuchter, J., Runfeldt, G., Bianco, R.A., Zhang, G., Prüfer, K., Orschiedt, J., Simm, A., Maier, P., Sager, M., Dresely, V., Krause, J., Meller, H. and Wehner, D., 2026. Reconstructing their genomes confirms the historically attested genealogy of the two medieval emperors Otto I (“the Great”) and Heinrich II (“Saint Henry”). biorxiv, 1–15. 10.64898/2026.03.18.712637

136. Roch, S., 2006. A short proof that phylogenetic tree reconstruction by maximum likelihood is hard. IEEE/ACM Trans. Comput. Biol. Bioinform. 3(1), 92–94. 10.1109/TCBB.2006.4

137. Rogaev, E.I., Grigorenko, A.P., Moliaka, Y.K., Faskhutdinova, G., Goltsov, A., Lahti, A., Hildebrandt, C., Kittler, E.L.W. and Morozova, I., 2009. Genomic identification in the historical case of the Nicholas II royal family. Proc. Natl. Acad. Sci. U. S. A. 106(13), 5258–5263. 10.1073/pnas.0811190106

138. Rollo, F., Ermini, L., Luciani, S., Marota, I., Olivieri, C. and Luiselli, D., 2006. Fine characterization of the Iceman’s mtDNA haplogroup. Am. J. Phys. Anthropol. 130(4), 557–564. 10.1002/ajpa.20384

139. Roshan, U.W., Moret, B.M.E., Warnow, T. and Williams, T.L., 2004. Rec-I-DCM3: A fast algorithmic technique for reconstructing large phylogenetic trees, in: Proceedings of the 2004 IEEE Computational Systems Bioinformatics Conference. IEEE, pp. 94–105. 10.1109/CSB.2004.1332422

140. Sawyer, S., Renaud, G., Viola, B., Hublin, J.J., Gansauge, M.T., Shunkov, M. V., Derevianko, A.P., Prüfer, K., Kelso, J. and Pääbo, S., 2015. Nuclear and mitochondrial DNA sequences from two Denisovan individuals. Proc. Natl. Acad. Sci. U. S. A. 112(51), 15696–15700. 10.1073/pnas.1519905112

141. Sayers, E.W., Cavanaugh, M., Frisse, L., Pruitt, K.D., Schneider, V.A., Underwood, B.A., Yankie, L. and Karsch-Mizrachi, I., 2025. GenBank 2025 update. Nucleic Acids Res. 53(D1), D56–D61. 10.1093/nar/gkae1114

142. Sayyari, E., Whitfield, J.B. and Mirarab, S., 2017. Fragmentary gene sequences negatively impact gene tree and species tree reconstruction. Mol. Biol. Evol. 34(12), 3279–3291. 10.1093/molbev/msx261

143. Schliep, K.P., 2011. phangorn: Phylogenetic analysis in R. Bioinformatics 27(4), 592–593. 10.1093/bioinformatics/btq706

144. Scholz, C.A., Johnson, T.C., Cohen, A.S., King, J.W., Peck, J.A., Overpeck, J.T., Talbot, M.R., Brown, E.T., Kalindekafe, L., Amoako, P.Y.O., Lyons, R.P., Shanahan, T.M., Castañeda, I.S., Heil, C.W., Forman, S.L., McHargue, L.R., Beuning, K.R., Gomez, J. and Pierson, J., 2007. East African megadroughts between 135 and 75 thousand years ago and bearing on early-modern human origins. Proc. Natl. Acad. Sci. U. S. A. 104(42), 16416–16421. 10.1073/pnas.0703874104

145. Schönherr, S., Weissensteiner, H., Kronenberg, F. and Forer, L., 2023. Haplogrep 3 – An interactive haplogroup classification and analysis platform. Nucleic Acids Res. 51(W1), W263–W268. 10.1093/nar/gkad284

146. Silva, M., Alshamali, F., Silva, P., Carrilho, C., Mandlate, F., Jesus Trovoada, M., Černý, V., Pereira, L. and Soares, P., 2015. 60,000 years of interactions between Central and Eastern Africa documented by major African mitochondrial haplogroup L2. Sci. Rep. 5(1), 12526. 10.1038/srep12526

147. Siva, N., 2008. 1000 Genomes project. Nat. Biotechnol. 26(3), 256. 10.1038/nbt0308-256b

148. Smirnov, V. and Warnow, T., 2021. Phylogeny estimation given sequence length heterogeneity. Syst. Biol. 70(2), 268–282. 10.1093/sysbio/syaa058

149. Smirnov, V. and Warnow, T., 2020. Unblended disjoint tree merging using GTM improves species tree estimation. BMC Genomics 21(S2), 235. 10.1186/s12864-020-6605-1

150. Soares, P.A., Ermini, L., Thomson, N., Mormina, M., Rito, T., Röhl, A., Salas, A., Oppenheimer, S., Macaulay, V. and Richards, M.B., 2009. Correcting for purifying selection: An improved human mitochondrial molecular clock. Am. J. Hum. Genet. 84(6), 740–759. 10.1016/j.ajhg.2009.05.001

151. Soares, P.A., Rito, T., Trejaut, J., Mormina, M., Hill, C., Tinkler-Hundal, E., Braid, M., Clarke, D.J., Loo, J.-H., Thomson, N., Denham, T., Donohue, M., Macaulay, V., Lin, M., Oppenheimer, S. and Richards, M.B., 2011. Ancient voyaging and Polynesian origins. Am. J. Hum. Genet. 88(2), 239–247. 10.1016/j.ajhg.2011.01.009

152. Soares, P.A., Trejaut, J.A., Rito, T., Cavadas, B., Hill, C., Eng, K.K., Mormina, M., Brandão, A., Fraser, R.M., Wang, T.-Y., Loo, J.-H., Snell, C., Ko, T.-M., Amorim, A., Pala, M., Macaulay, V., Bulbeck, D., Wilson, J.F., Gusmão, L., Pereira, L., Oppenheimer, S., Lin, M. and Richards, M.B., 2016. Resolving the ancestry of Austronesian-speaking populations. Hum. Genet. 135(3), 309–326. 10.1007/s00439-015-1620-z

153. Sümer, A.P., Rougier, H., Villalba-Mouco, V., Huang, Y., Iasi, L.N.M., Essel, E., Bossoms Mesa, A., Furtwaengler, A., Peyrégne, S., de Filippo, C., Rohrlach, A.B., Pierini, F., Mafessoni, F., Fewlass, H., Zavala, E.I., Mylopotamitaki, D., Bianco, R.A., Schmidt, A., Zorn, J., Nickel, B., Patova, A., Posth, C., Smith, G.M., Ruebens, K., Sinet-Mathiot, V., Stoessel, A., Dietl, H., Orschiedt, J., Kelso, J., Zeberg, H., Bos, K.I., Welker, F., Weiss, M., McPherron, S.P., Schüler, T., Hublin, J.-J., Velemínský, P., Brůžek, J., Peter, B.M., Meyer, M., Meller, H., Ringbauer, H., Hajdinjak, M., Prüfer, K. and Krause, J., 2025. Earliest modern human genomes constrain timing of Neanderthal admixture. Nature 638(8051), 711–717. 10.1038/s41586-024-08420-x

154. Tackney, J.C., Potter, B.A., Raff, J., Powers, M., Watkins, W.S., Warner, D., Reuther, J.D., Irish, J.D. and O’Rourke, D.H., 2015. Two contemporaneous mitogenomes from terminal Pleistocene burials in eastern Beringia. Proc. Natl. Acad. Sci. U. S. A. 112(45), 13833–13838. 10.1073/pnas.1511903112

155. Tambets, K., Rootsi, S., Kivisild, T., Help, H., Serk, P., Loogväli, E.-L., Tolk, H.-V., Reidla, M., Metspalu, E., Pliss, L., Balanovsky, O., Pshenichnov, A., Balanovska, E., Gubina, M., Zhadanov, S., Osipova, L., Damba, L., Voevoda, M., Kutuev, I., Bermisheva, M., Khusnutdinova, E., Gusar, V., Grechanina, E., Parik, J., Pennarun, E., Richard, C., Chaventre, A., Moisan, J.-P., Barać, L., Peričić, M., Rudan, P., Terzić, R., Mikerezi, I., Krumina, A., Baumanis, V., Koziel, S., Rickards, O., De Stefano, G.F., Anagnou, N., Pappa, K.I., Michalodimitrakis, E., Ferák, V., Füredi, S., Komel, R., Beckman, L. and Villems, R., 2004. The western and eastern roots of the Saami—the story of genetic “outliers” told by mitochondrial DNA and Y chromosomes. Am. J. Hum. Genet. 74(4), 661–682. 10.1086/383203

156. Tamm, E., Kivisild, T., Reidla, M., Metspalu, M., Smith, D.G., Mulligan, C.J., Bravi, C.M., Rickards, O., Martinez-Labarga, C., Khusnutdinova, E.K., Fedorova, S.A., Golubenko, M. V., Stepanov, V.A., Gubina, M.A., Zhadanov, S.I., Ossipova, L.P., Damba, L., Voevoda, M.I., Dipierri, J.E., Villems, R. and Malhi, R.S., 2007. Beringian standstill and spread of native American founders. PLoS One 2(9), e829. 10.1371/journal.pone.0000829

157. Teschler-Nicola, M., Fernandes, D., Händel, M., Einwögerer, T., Simon, U., Neugebauer-Maresch, C., Tangl, S., Heimel, P., Dobsak, T., Retzmann, A., Prohaska, T., Irrgeher, J., Kennett, D.J., Olalde, I., Reich, D. and Pinhasi, R., 2020. Ancient DNA reveals monozygotic newborn twins from the Upper Palaeolithic. Commun. Biol. 3(1), 650. 10.1038/s42003-020-01372-8

158. Thangaraj, K., Chaubey, G., Kivisild, T., Reddy, A.G., Singh, V.K., Rasalkar, A.A. and Singh, L., 2005. Reconstructing the origin of Andaman Islanders. Science. 308(5724), 996–996. 10.1126/science.1109987

159. Tian, J.-Y., Wang, H.-W., Li, Y.-C., Zhang, W., Yao, Y.-G., van Straten, J., Richards, M.B. and Kong, Q.-P., 2015. A genetic contribution from the Far East into Ashkenazi Jews via the ancient Silk Road. Sci. Rep. 5(1), 8377. 10.1038/srep08377

160. Tierney, J.E., Pausata, F.S.R. and deMenocal, P.B., 2017. Rainfall regimes of the Green Sahara. Sci. Adv. 3(1). 10.1126/sciadv.1601503

161. To, T.H., Jung, M., Lycett, S. and Gascuel, O., 2016. Fast dating using least-squares criteria and algorithms. Syst. Biol. 65(1), 82–97. 10.1093/sysbio/syv068

162. Torres, A., Goloboff, P.A. and Catalano, S.A., 2022. Parsimony analysis of phylogenomic datasets (I): scripts and guidelines for using TNT (Tree Analysis using New Technology). Cladistics 38(1), 103–125. 10.1111/cla.12477

163. Trivedi, R., Sitalaximi, T., Banerjee, J., Singh, A., Sircar, P.K. and Kashyap, V.K., 2006. Molecular insights into the origins of the Shompen, a declining population of the Nicobar archipelago. J. Hum. Genet. 51(3), 217–226. 10.1007/s10038-005-0349-2

164. Turakhia, Y., Thornlow, B., Hinrichs, A.S., De Maio, N., Gozashti, L., Lanfear, R., Haussler, D. and Corbett-Detig, R., 2021. Ultrafast Sample placement on Existing tRees (UShER) enables real-time phylogenetics for the SARS-CoV-2 pandemic. Nat. Genet. 53(6), 809–816. 10.1038/s41588-021-00862-7

165. Vai, S., Sarno, S., Lari, M., Luiselli, D., Manzi, G., Gallinaro, M., Mataich, S., Hübner, A., Modi, A., Pilli, E., Tafuri, M.A., Caramelli, D. and di Lernia, S., 2019. Ancestral mitochondrial N lineage from the Neolithic ‘green’ Sahara. Sci. Rep. 9(1), 3530. 10.1038/s41598-019-39802-1

166. Vallini, L., Marciani, G., Aneli, S., Bortolini, E., Benazzi, S., Pievani, T. and Pagani, L., 2022. Genetics and material culture support repeated expansions into Paleolithic Eurasia from a population hub out of Africa. Genome Biol. Evol. 14(4). 10.1093/gbe/evac045

167. van Oven, M., 2015. PhyloTree Build 17: Growing the human mitochondrial DNA tree. Forensic Sci. Int. Genet. Suppl. Ser. 5, e392–e394. 10.1016/j.fsigss.2015.09.155

168. van Oven, M. and Kayser, M., 2009. Updated comprehensive phylogenetic tree of global human mitochondrial DNA variation. Hum. Mutat. 30(2), 386–394. 10.1002/humu.20921

169. Verdugo, C., Zhu, K., Kassadjikova, K., Berg, L., Forst, J., Galloway, A., Brady, J.E. and Fehren-Schmitz, L., 2020. An investigation of ancient Maya intentional dental modification practices at Midnight Terror Cave using anthroposcopic and paleogenomic methods. J. Archaeol. Sci. 115, 105096. 10.1016/j.jas.2020.105096

170. Vigilant, L., Stoneking, M., Harpending, H., Hawkes, K. and Wilson, A.C., 1991. African populations and the evolution of human mitochondrial DNA. Science. 253(5027), 1503–1507. 10.1126/science.1840702

171. Villalba-Mouco, V., van de Loosdrecht, M.S., Rohrlach, A.B., Fewlass, H., Talamo, S., Yu, H., Aron, F., Lalueza-Fox, C., Cabello, L., Cantalejo Duarte, P., Ramos-Muñoz, J., Posth, C., Krause, J., Weniger, G.-C. and Haak, W., 2023. A 23,000-year-old southern Iberian individual links human groups that lived in Western Europe before and after the Last Glacial Maximum. Nat. Ecol. Evol. 7(4), 597–609. 10.1038/s41559-023-01987-0

172. Wang, K., Prüfer, K., Krause-Kyora, B., Childebayeva, A., Schuenemann, V.J., Coia, V., Maixner, F., Zink, A., Schiffels, S. and Krause, J., 2023. High-coverage genome of the Tyrolean Iceman reveals unusually high Anatolian farmer ancestry. Cell Genomics 3(9), 100377. 10.1016/j.xgen.2023.100377

173. Wang, L., Lam, T.T., Xu, S., Dai, Z., Zhou, L., Feng, T., Guo, P., Dunn, C.W., Jones, B.R., Bradley, T., Zhu, H., Guan, Y., Jiang, Y. and Yu, G., 2020. Treeio: An R package for phylogenetic tree input and output with richly annotated and associated data. Mol. Biol. Evol. 37(2), 599–603. 10.1093/molbev/msz240

174. Wright, A.M. and Wynd, B.M., 2024. Modeling of rate heterogeneity in datasets compiled for use with parsimony. bioRxiv, 1–22. 10.1101/2024.06.26.600858

175. Yao, Y.G., Salas, A., Logan, I. and Bandelt, H.J., 2009. mtDNA data mining in GenBank needs surveying. Am. J. Hum. Genet. 85(6), 929–933. 10.1016/j.ajhg.2009.10.023

176. Ye, C., Thornlow, B., Hinrichs, A., Kramer, A., Mirchandani, C., Torvi, D., Lanfear, R., Corbett-Detig, R. and Turakhia, Y., 2022. matOptimize: A parallel tree optimization method enables online phylogenetics for SARS-CoV-2. Bioinformatics 38(15), 3734–3740. 10.1093/bioinformatics/btac401

177. Zaharias, P. and Warnow, T., 2022. Recent progress on methods for estimating and updating large phylogenies. Philos. Trans. R. Soc. B 377(1861). 10.1098/rstb.2021.0244

178. Zeng, T.C., Vyazov, L.A., Kim, A., Flegontov, P., Sirak, K., Maier, R., Lazaridis, I., Akbari, A., Frachetti, M., Tishkin, A.A., Ryabogina, N.E., Agapov, S.A., Agapov, D.S., Alekseev, A.N., Boeskorov, G.G., Derevianko, A.P., Dyakonov, V.M., Enshin, D.N., Fribus, A. V., Frolov, Y. V., Grushin, S.P., Khokhlov, A.A., Kiryushin, K.Yu., Kiryushin, Y.F., Kitov, E.P., Kosintsev, P., Kovtun, I. V., Makarov, N.P., Morozov, V. V., Nikolaev, E.N., Rykun, M.P., Savenkova, T.M., Shchelchkova, M. V., Shirokov, V., Skochina, S.N., Sherstobitova, O.S., Slepchenko, S.M., Solodovnikov, K.N., Solovyova, E.N., Stepanov, A.D., Timoshchenko, A.A., Vdovin, A.S., Vybornov, A. V., Balanovska, E. V., Dryomov, S., Hellenthal, G., Kidd, K., Krause, J., Starikovskaya, E., Sukernik, R., Tatarinova, T., Thomas, M.G., Zhabagin, M., Callan, K., Cheronet, O., Fernandes, D., Keating, D., Candilio, F., Iliev, L., Kearns, A., Özdoğan, K.T., Mah, M., Micco, A., Michel, M., Olalde, I., Zalzala, F., Mallick, S., Rohland, N., Pinhasi, R., Narasimhan, V.M. and Reich, D., 2025. Ancient DNA reveals the prehistory of the Uralic and Yeniseian peoples. Nature 644(8075), 122–132. 10.1038/s41586-025-09189-3

179. Zhong, B., Deusch, O., Goremykin, V. V., Penny, D., Biggs, P.J., Atherton, R.A., Nikiforova, S. V. and Lockhart, P.J., 2011. Systematic error in seed plant phylogenomics. Genome Biol. Evol. 3(1), 1340–1348. 10.1093/gbe/evr105

