## Supplemental Figures for "Mitotree: The Universal Human Mitochondrial Reference Phylogeny at 10× the Resolution"

### L0d2 (Mitotree 2026−05−06)

*(Splitters >30,000 Years)*

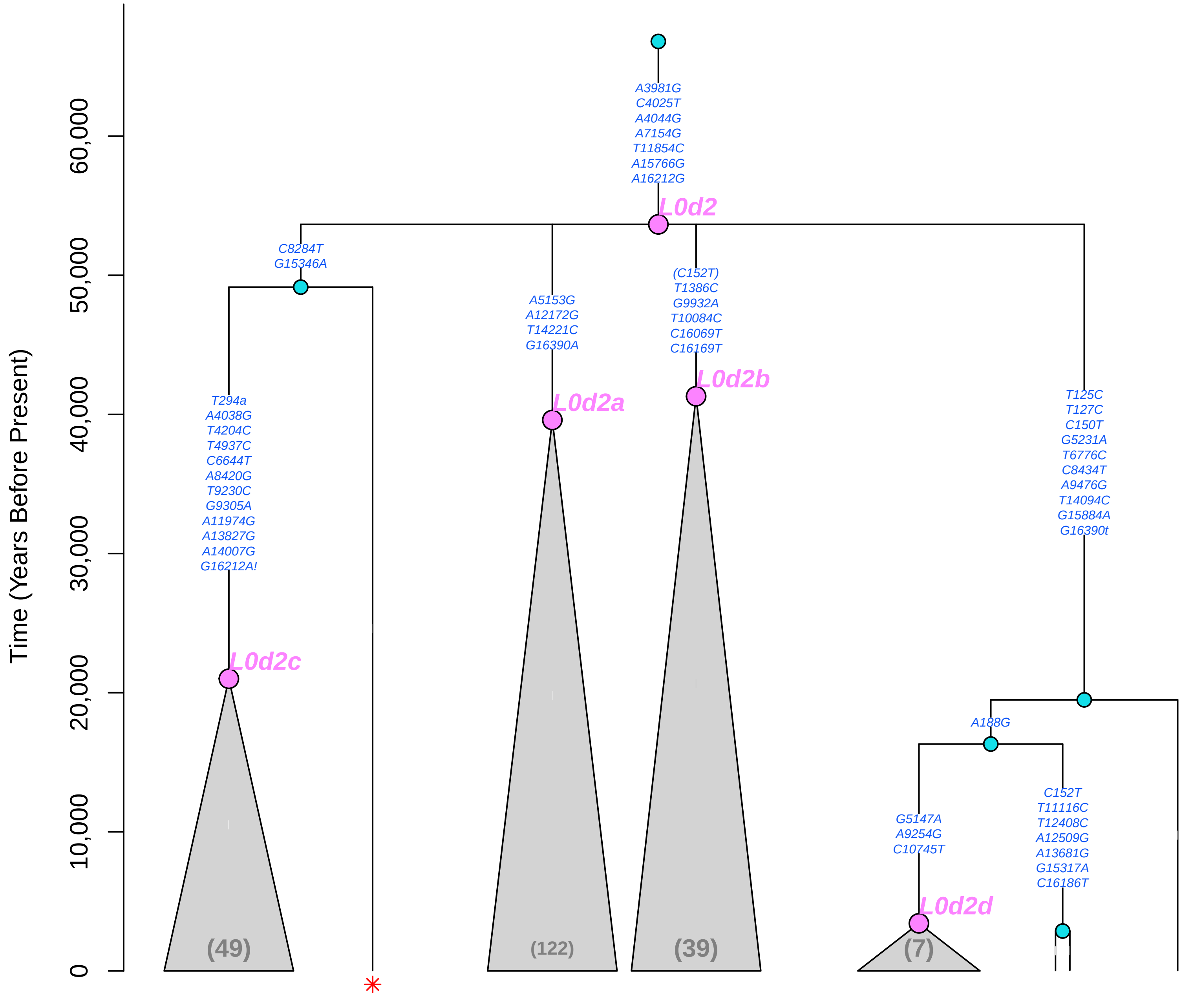

### L0f (Mitotree 2026−05−06)

*(Splitters >30,000 Years)*

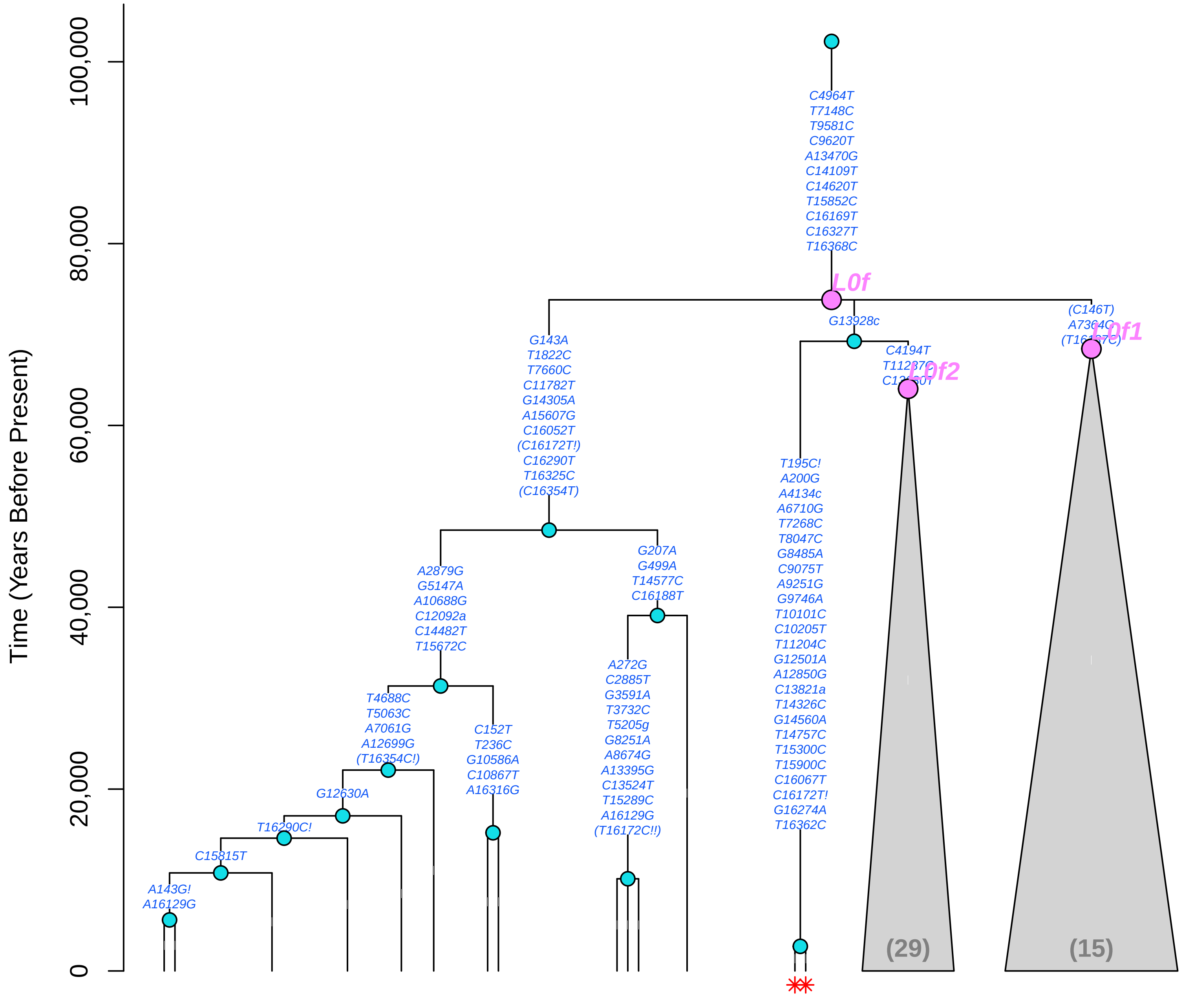

### L1c1 (Mitotree 2026−05−06)

*(Splitters >30,000 Years)*

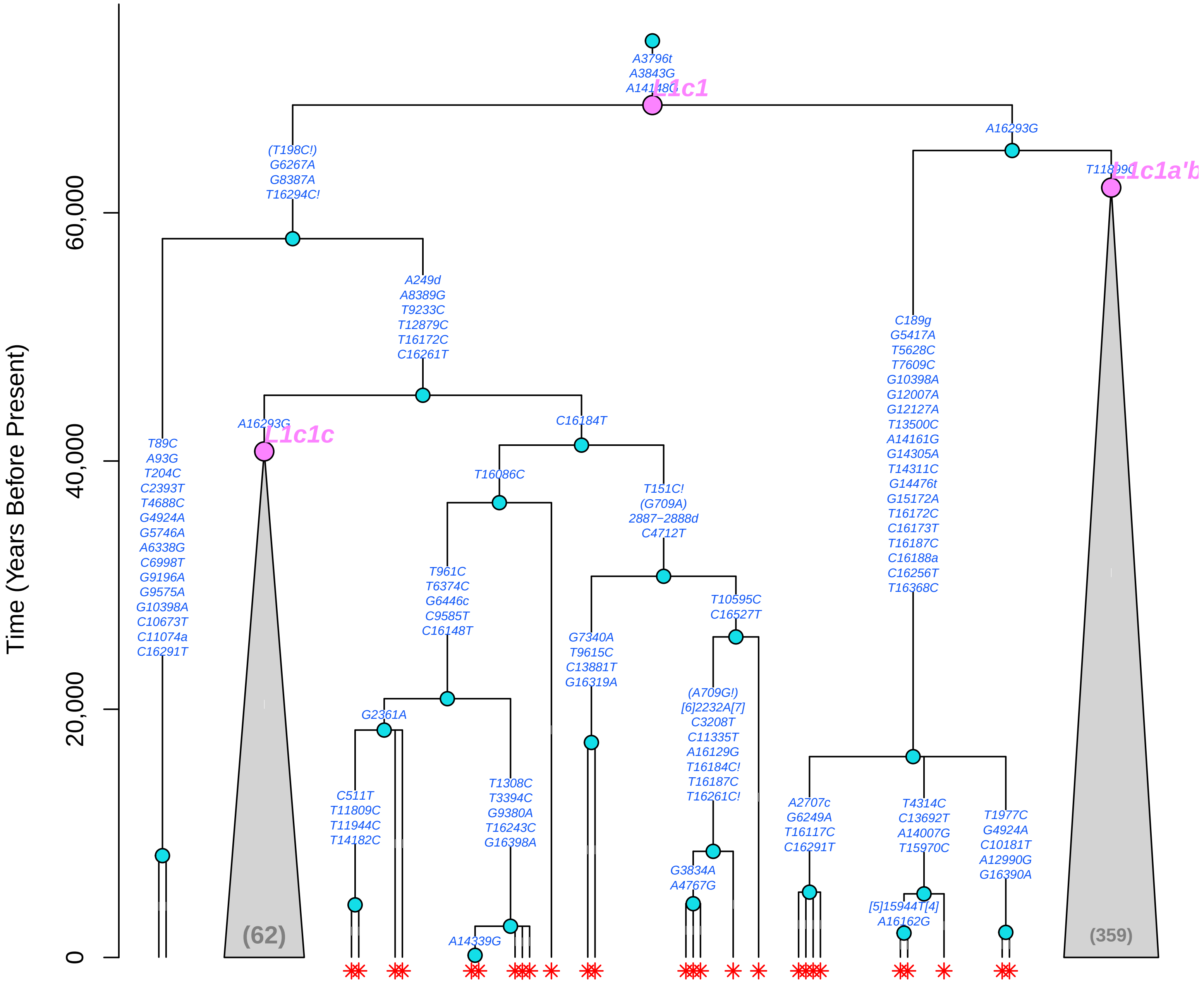

### L1c1a (Mitotree 2026−05−06)

*(Splitters >30,000 Years)*

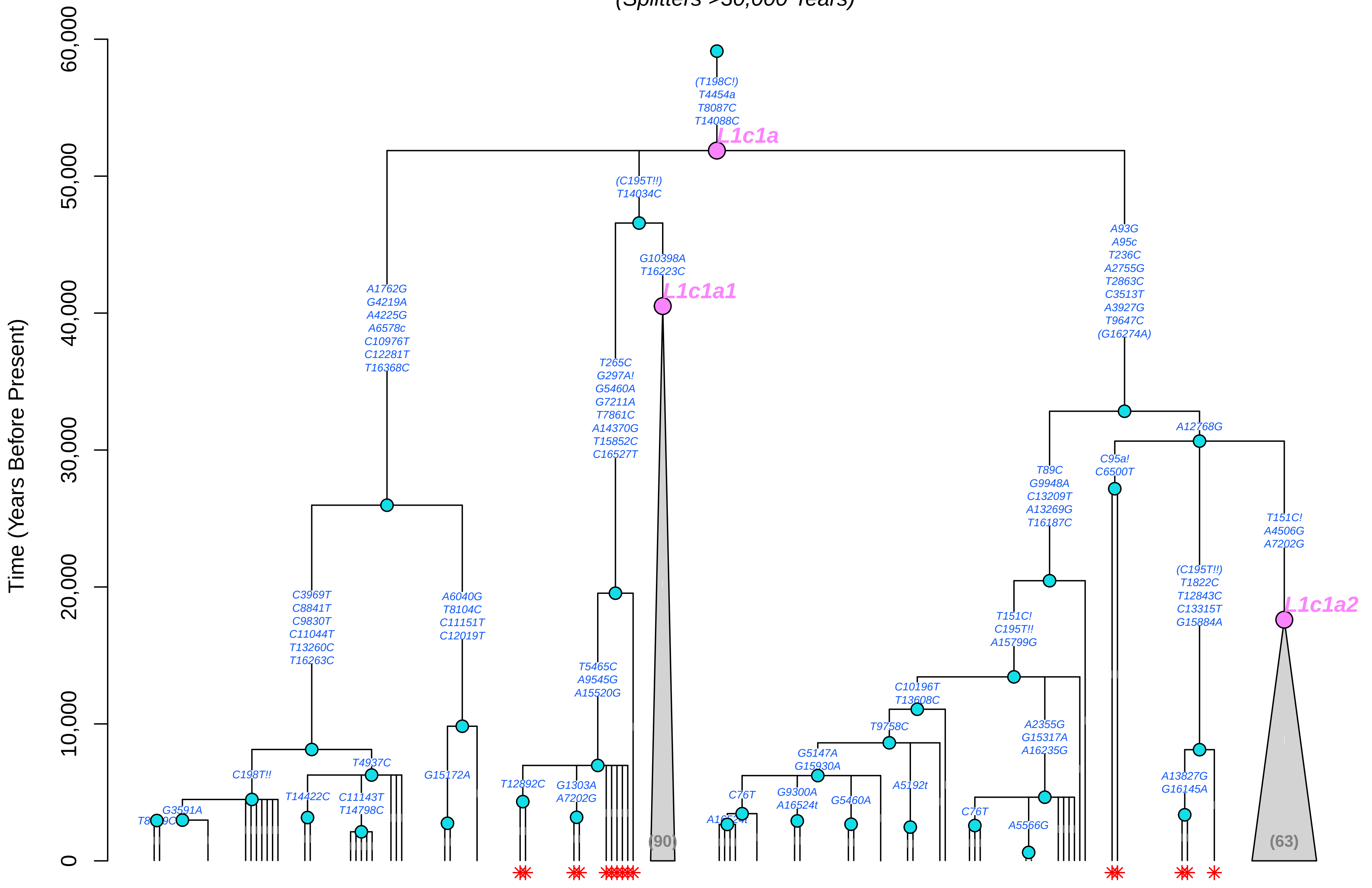

### L1c1a1 (Mitotree 2026−05−06)

*(Splitters >30,000 Years)*

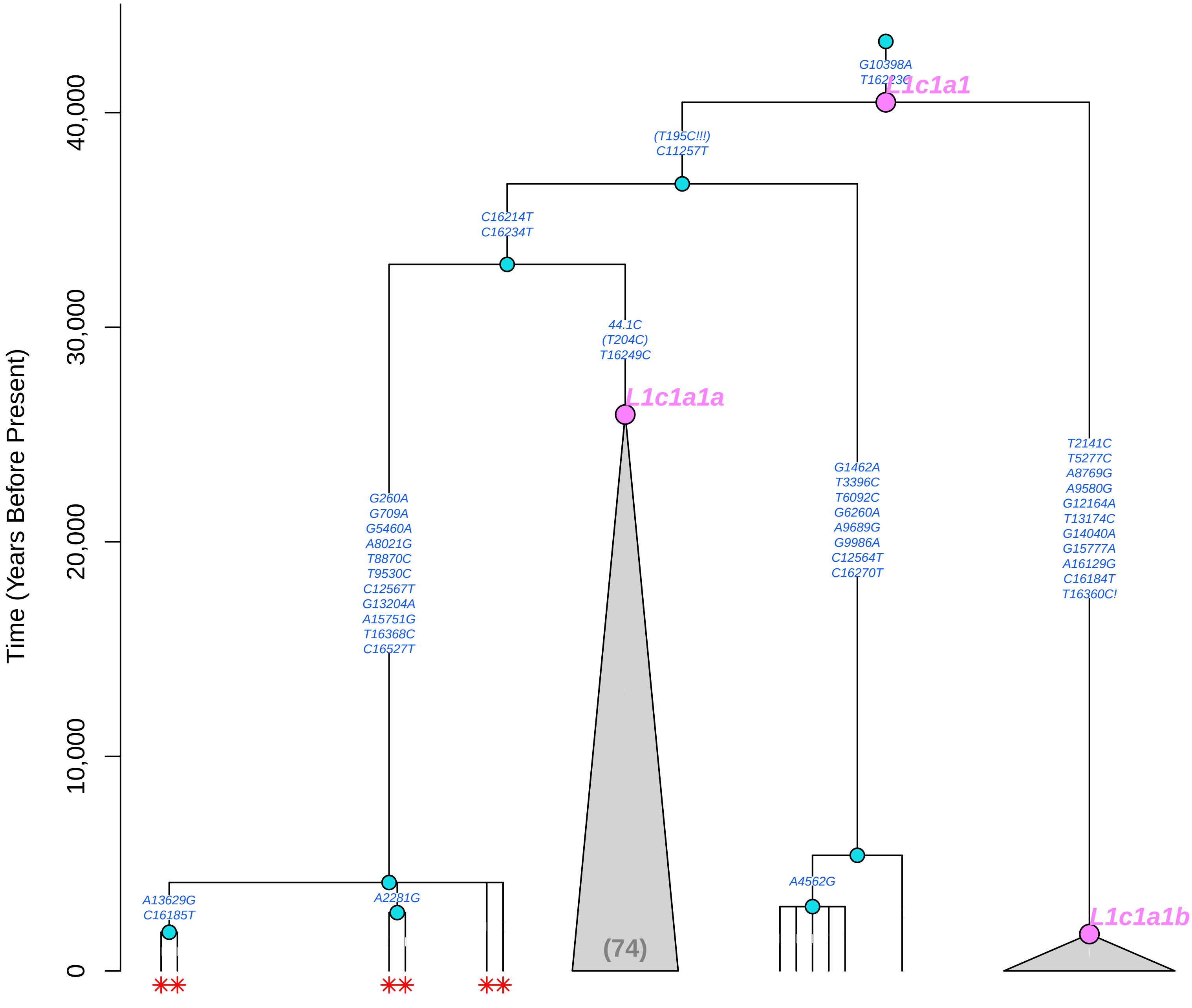

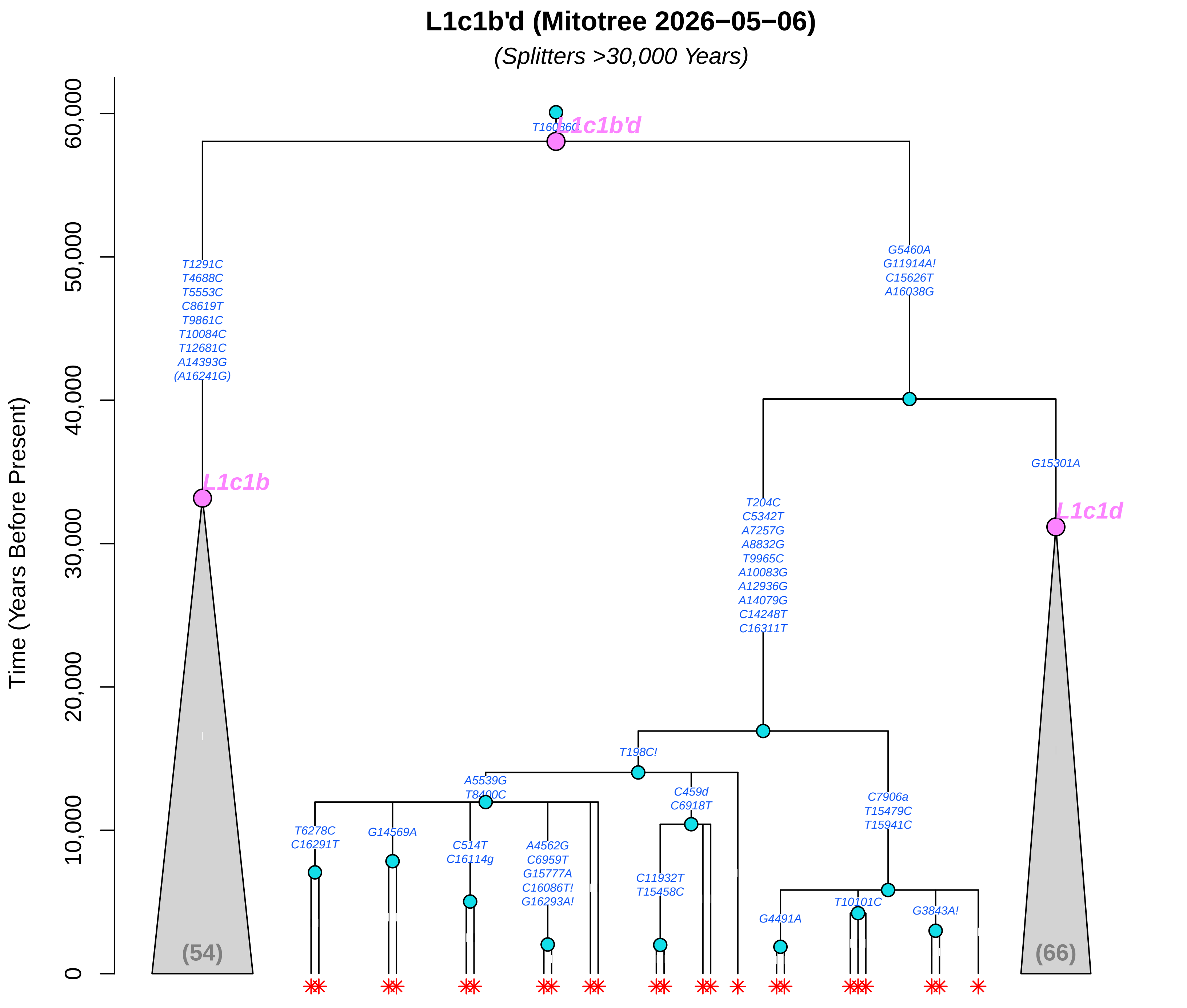

L1c1b'd (Mitotree 2026−05−06)
(Splitters >30,000 Years)
Time (Years Before Present)
0
10,000
20,000
30,000
40,000
50,000
60,000
T16036C
L1c1b'd
T1291C
T4688C
T5553C
C8619T
T9861C
T10084C
T12681C
A14393G
(A16241G)
L1c1b
(54)
G5460A
G11914A!
C15626T
A16038G
G15301A
L1c1d
(66)
T204C
C5342T
A7257G
A8832G
T9965C
A10083G
A12936G
A14079G
C14248T
C16311T
T198C!
A5539G
T8400C
C459d
C6918T
C7906a
T15479C
T15941C
T6278C
C16291T
G14569A
C514T
C16114g
A4562G
C6959T
G15777A
C16086T!
G16293A!
C11932T
T15458C
G4491A
T10101C
G3843A!

### L1c2'4 (Mitotree 2026−05−06)

*(Splitters >30,000 Years)*

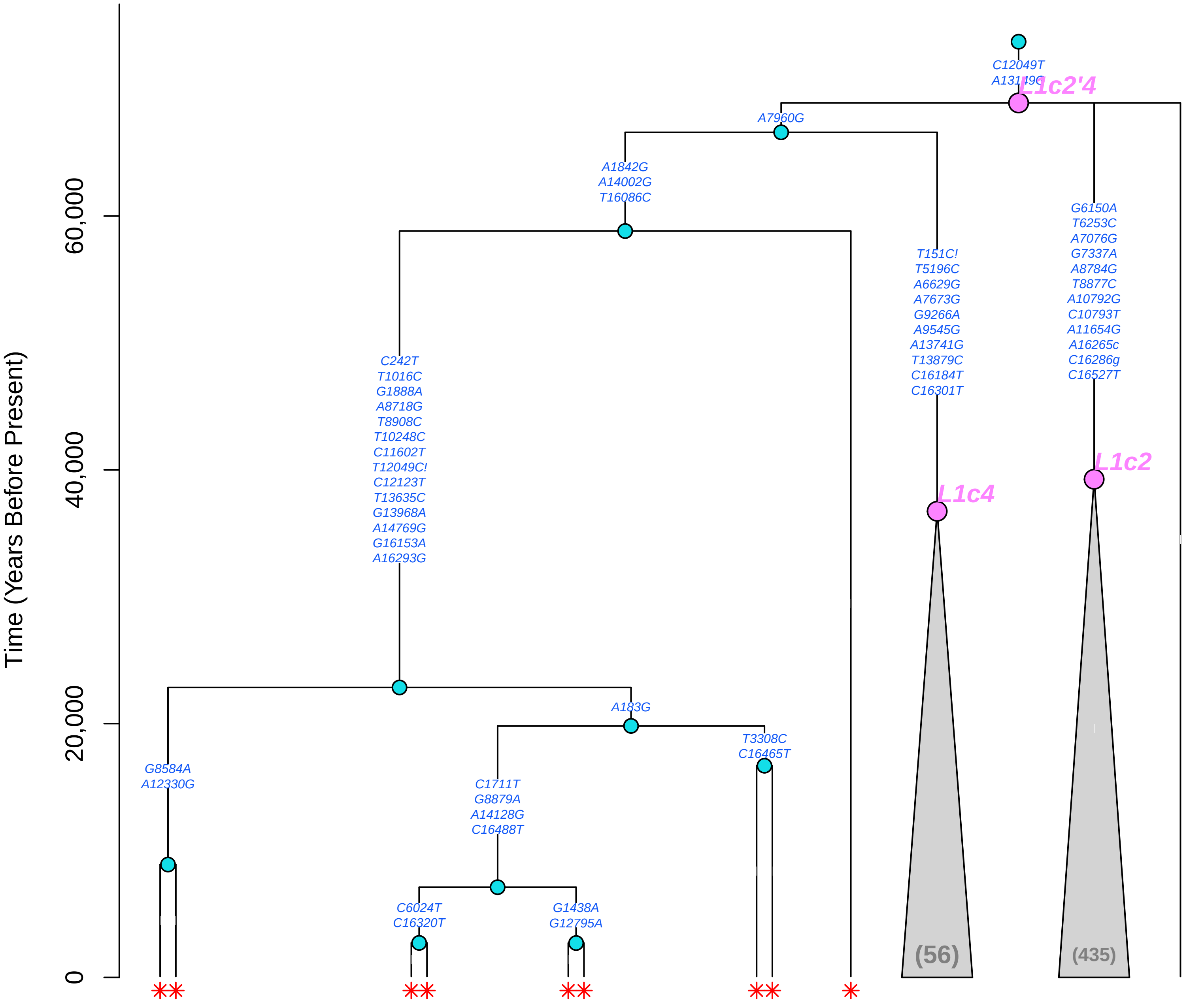

#### L2 (Mitotree 2026−05−06)

*(Splitters >30,000 Years)*

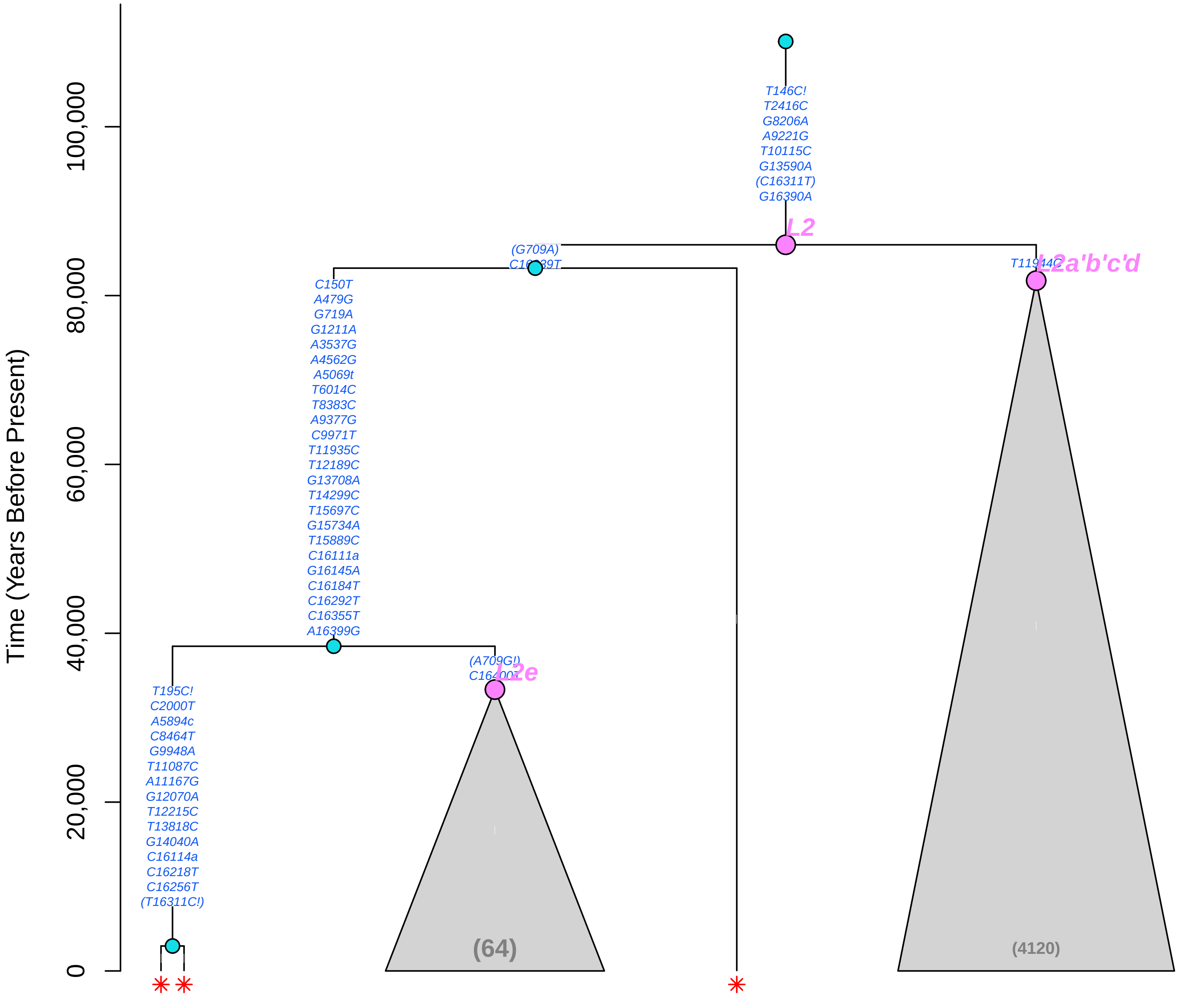

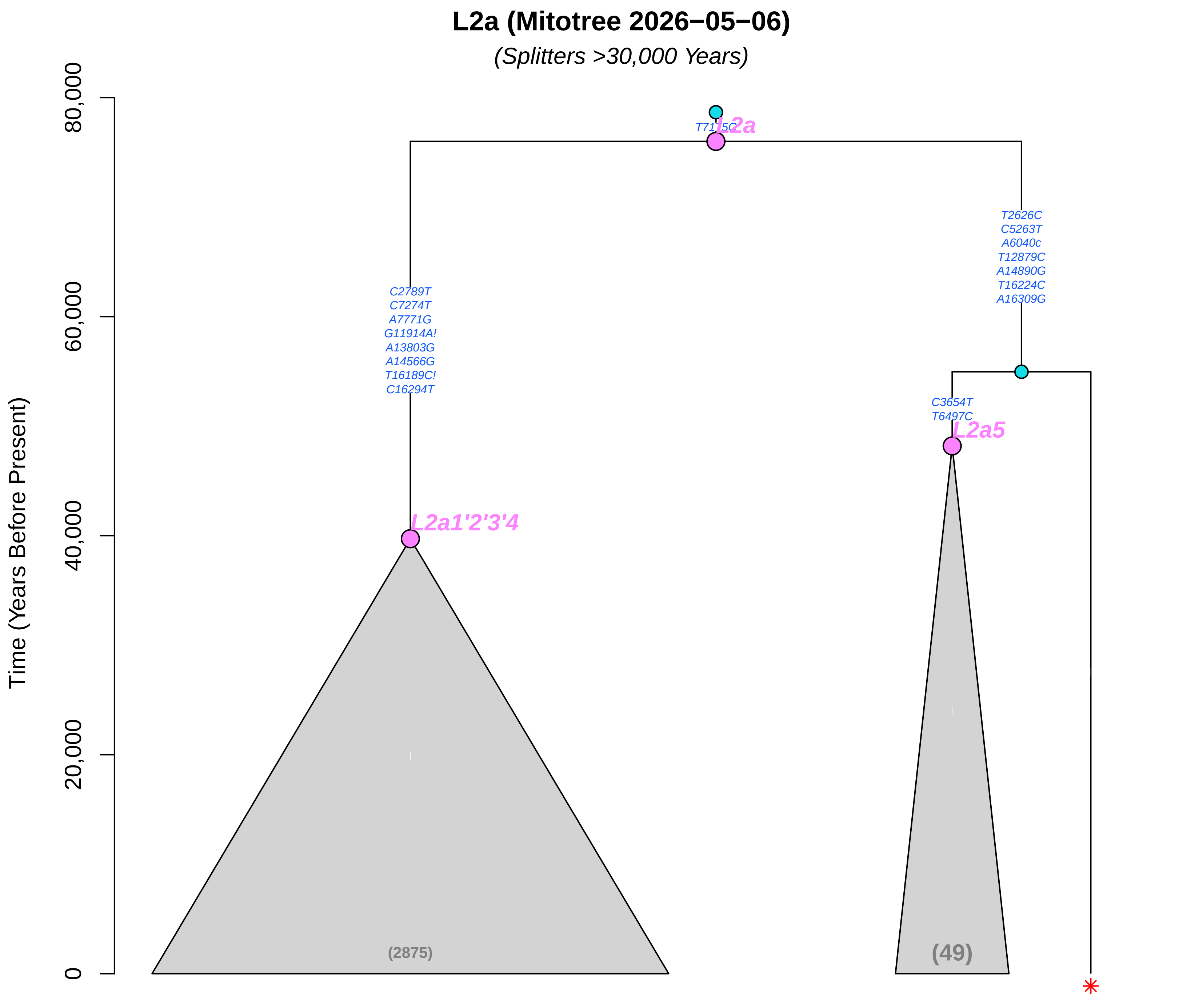
L2a (Mitotree 2026−05−06)
(Splitters >30,000 Years)
Time (Years Before Present)
80,000
60,000
40,000
20,000
0
T7175C
L2a
C2789T
C7274T
A7771G
G11914A!
A13803G
A14566G
T16189C!
C16294T
L2a1'2'3'4
T2626C
C5263T
A6040c
T12879C
A14890G
T16224C
A16309G
C3654T
T6497C
L2a5
(2875)
(49)

### L3e'i'k'x (Mitotree 2026−05−06)

*(Splitters >30,000 Years)*

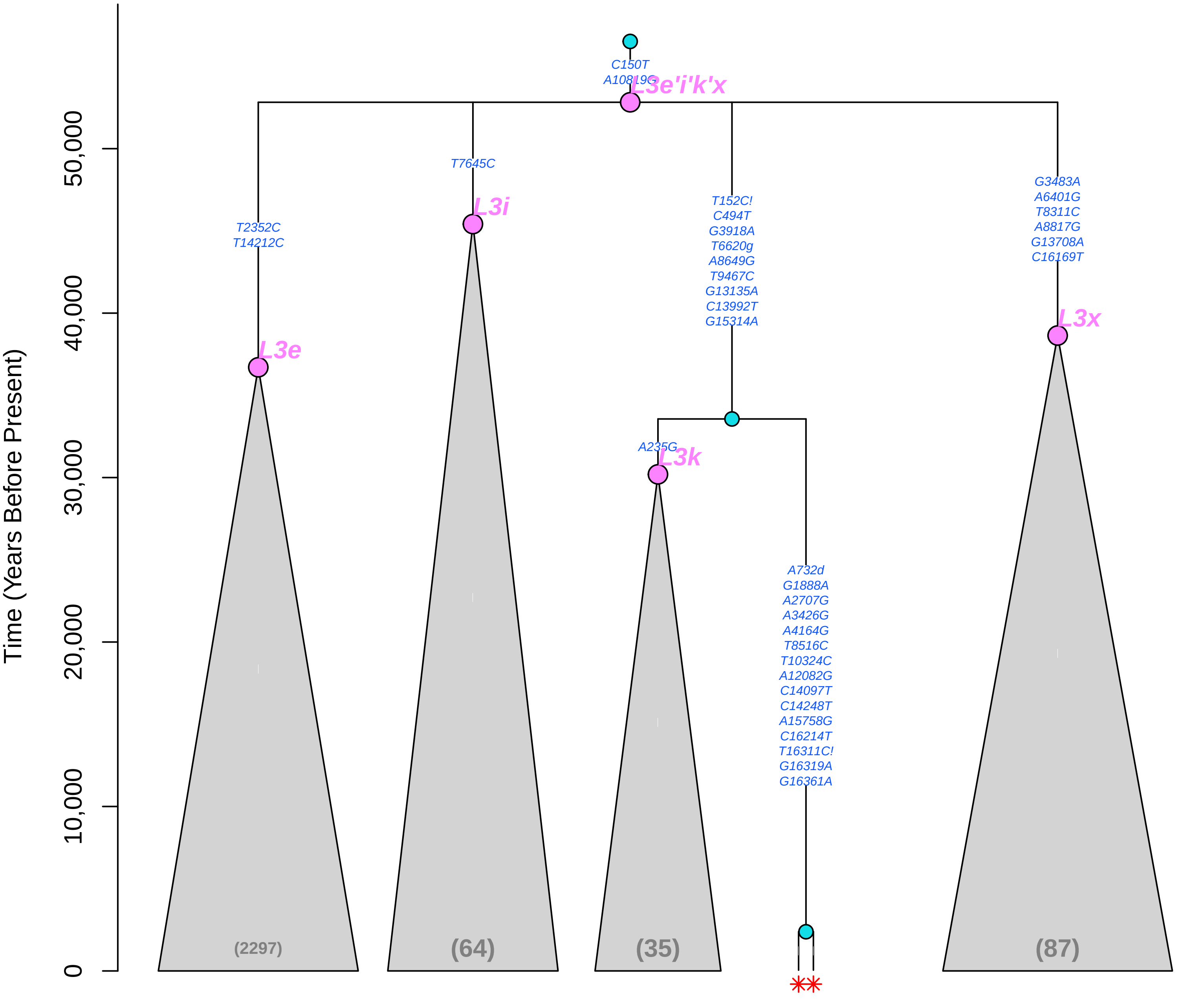

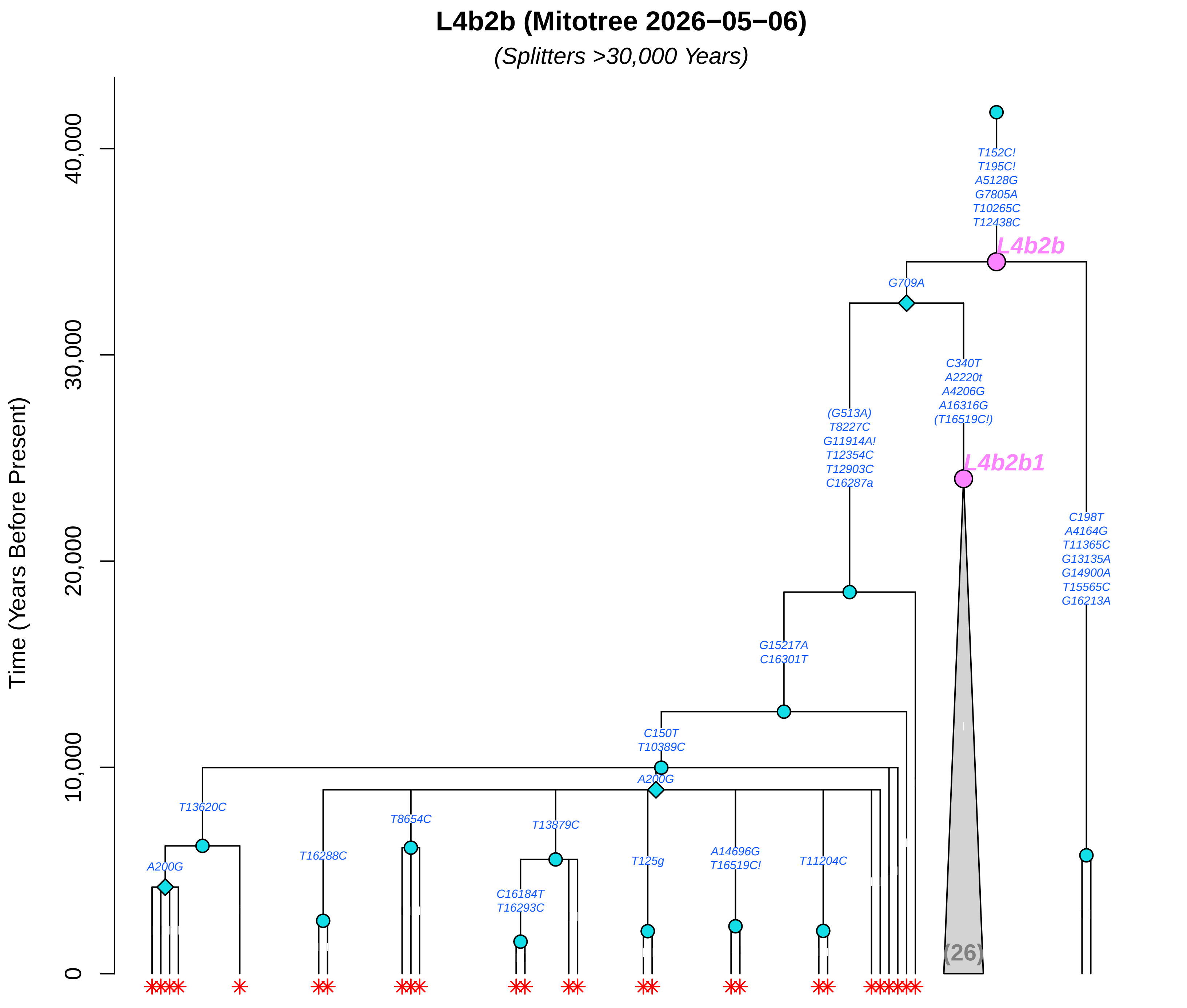

L4b2b (Mitotree 2026−05−06)
(Splitters >30,000 Years)
Time (Years Before Present)
40,000
30,000
20,000
10,000
0
T152C!
T195C!
A5128G
G7805A
T10265C
T12438C
L4b2b
G709A
C340T
A2220t
A4206G
A16316G
(T16519C!)
L4b2b1
(G513A)
T8227C
G11914A!
T12354C
T12903C
C16287a
C198T
A4164G
T11365C
G13135A
G14900A
T15565C
G16213A
G15217A
C16301T
C150T
T10389C
A200G
T13620C
T8654C
T13879C
A200G
T16288C
C16184T
T16293C
T125g
A14696G
T16519C!
T11204C
(26)

### L7a (Mitotree 2026−05−06)

*(Splitters >30,000 Years)*

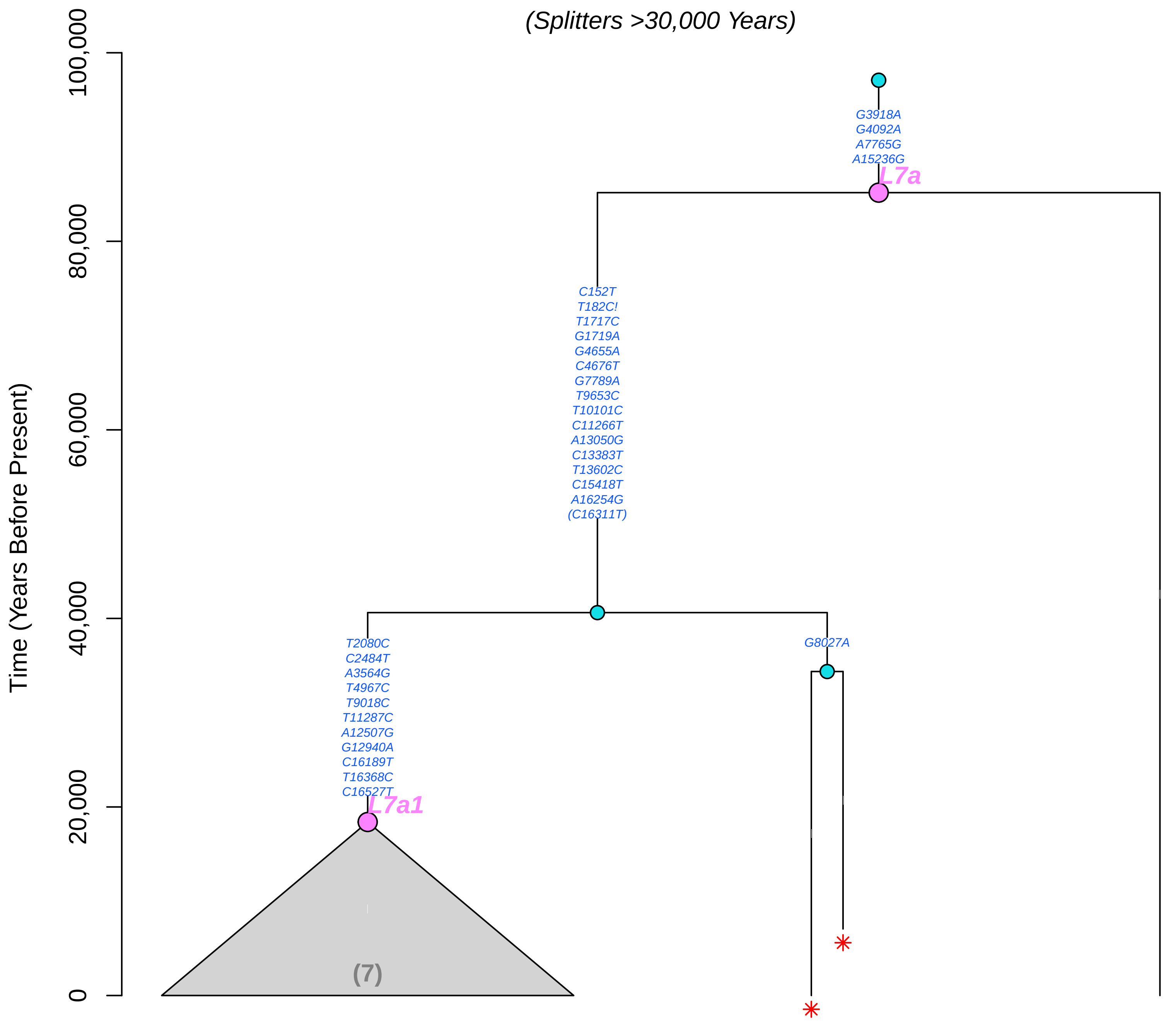

**M (Mitotree 2026-05-06)**

*(Splitters >30,000 Years)*

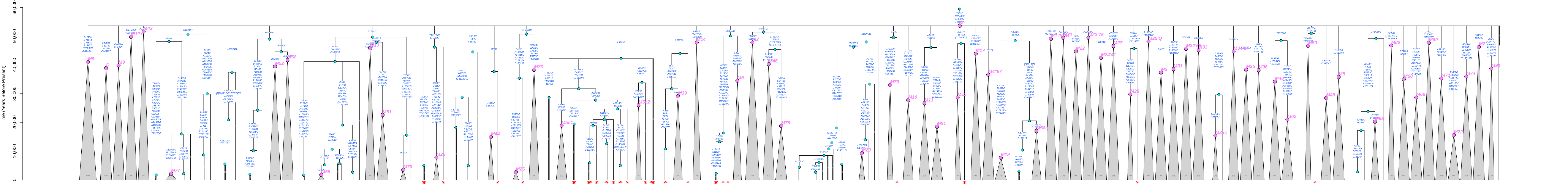

**M13 (Mitotree 2026−05−06)**

*(Splitters >30,000 Years)*

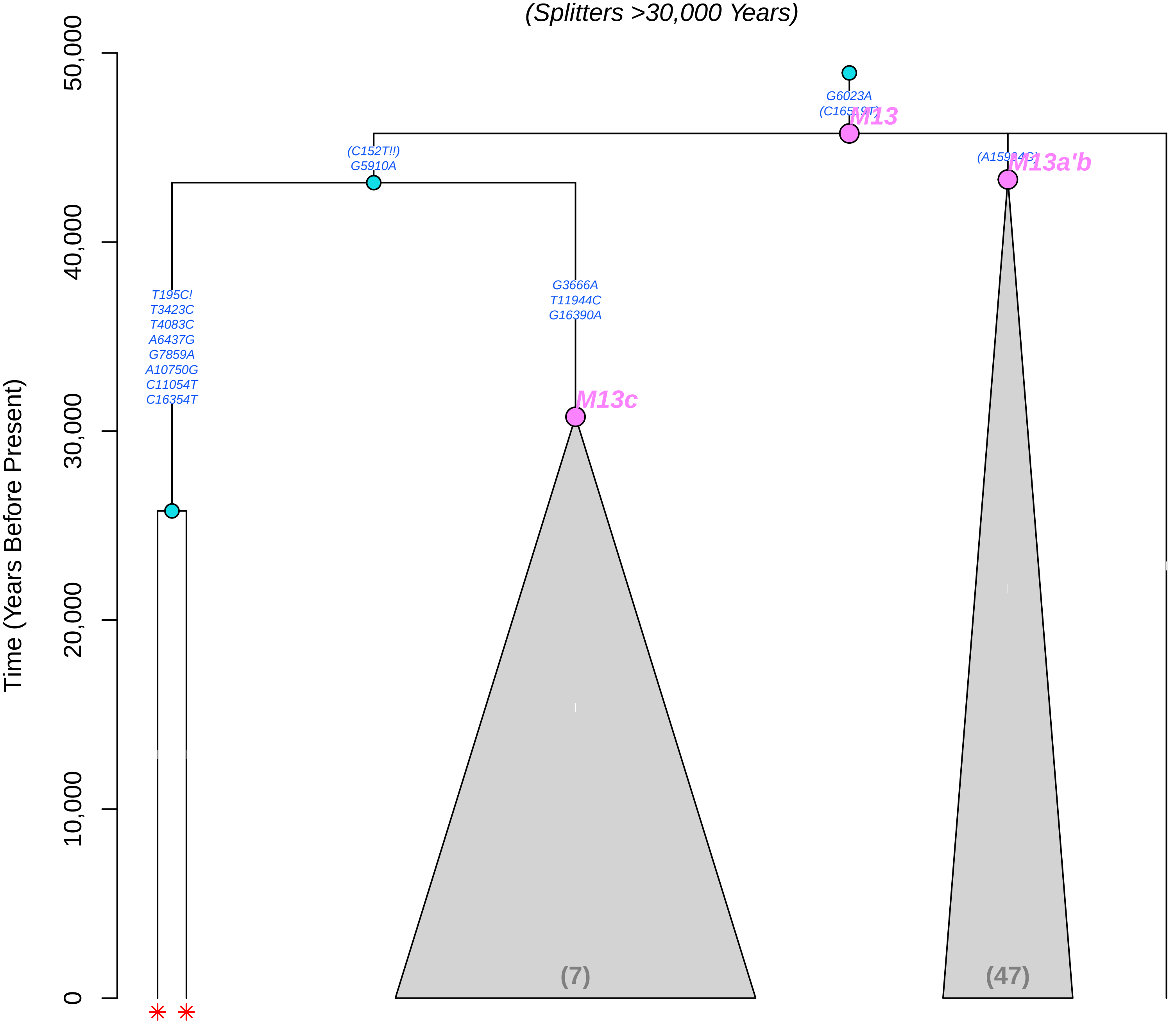

### M17c (Mitotree 2026−05−06)

*(Splitters >30,000 Years)*

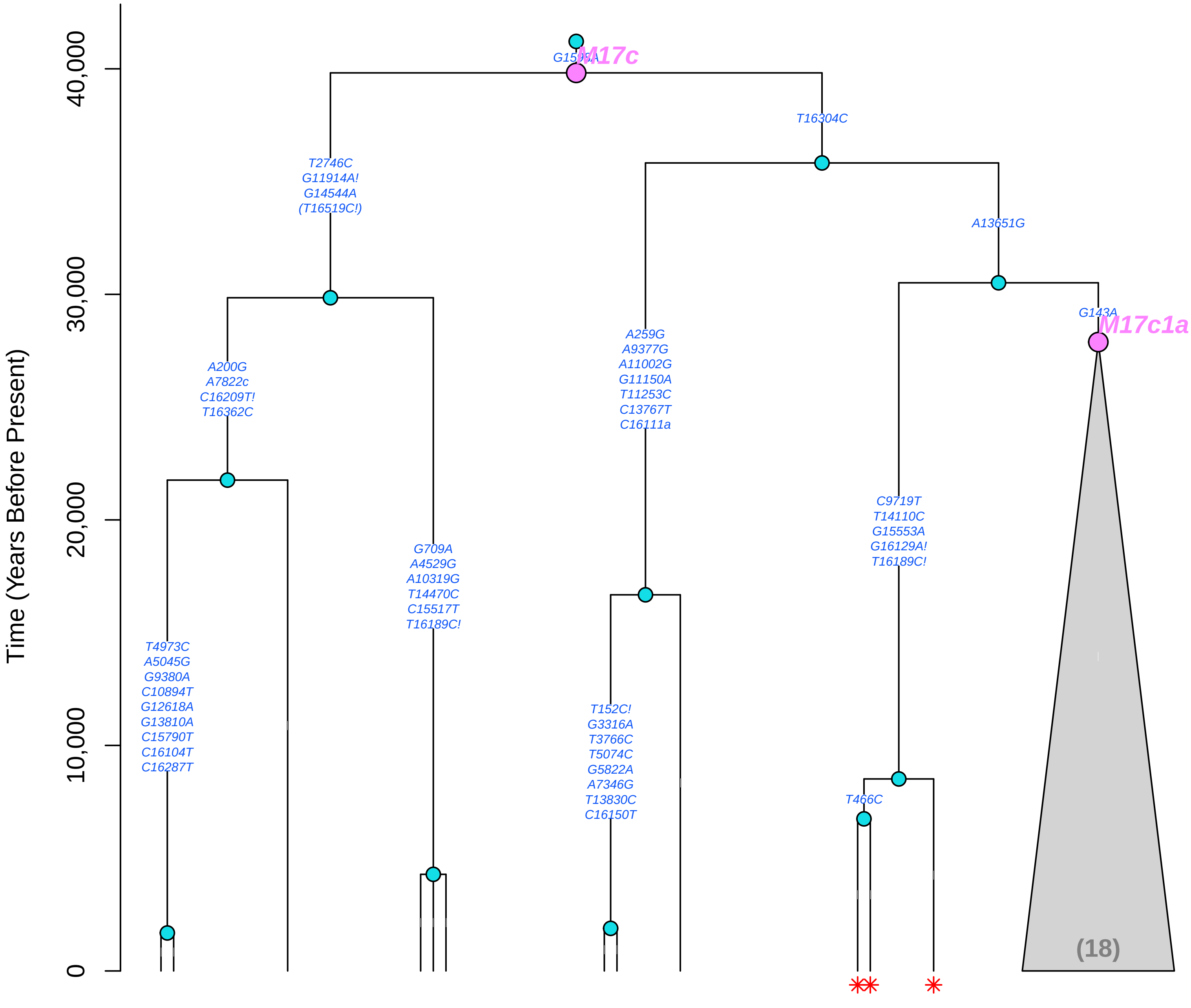

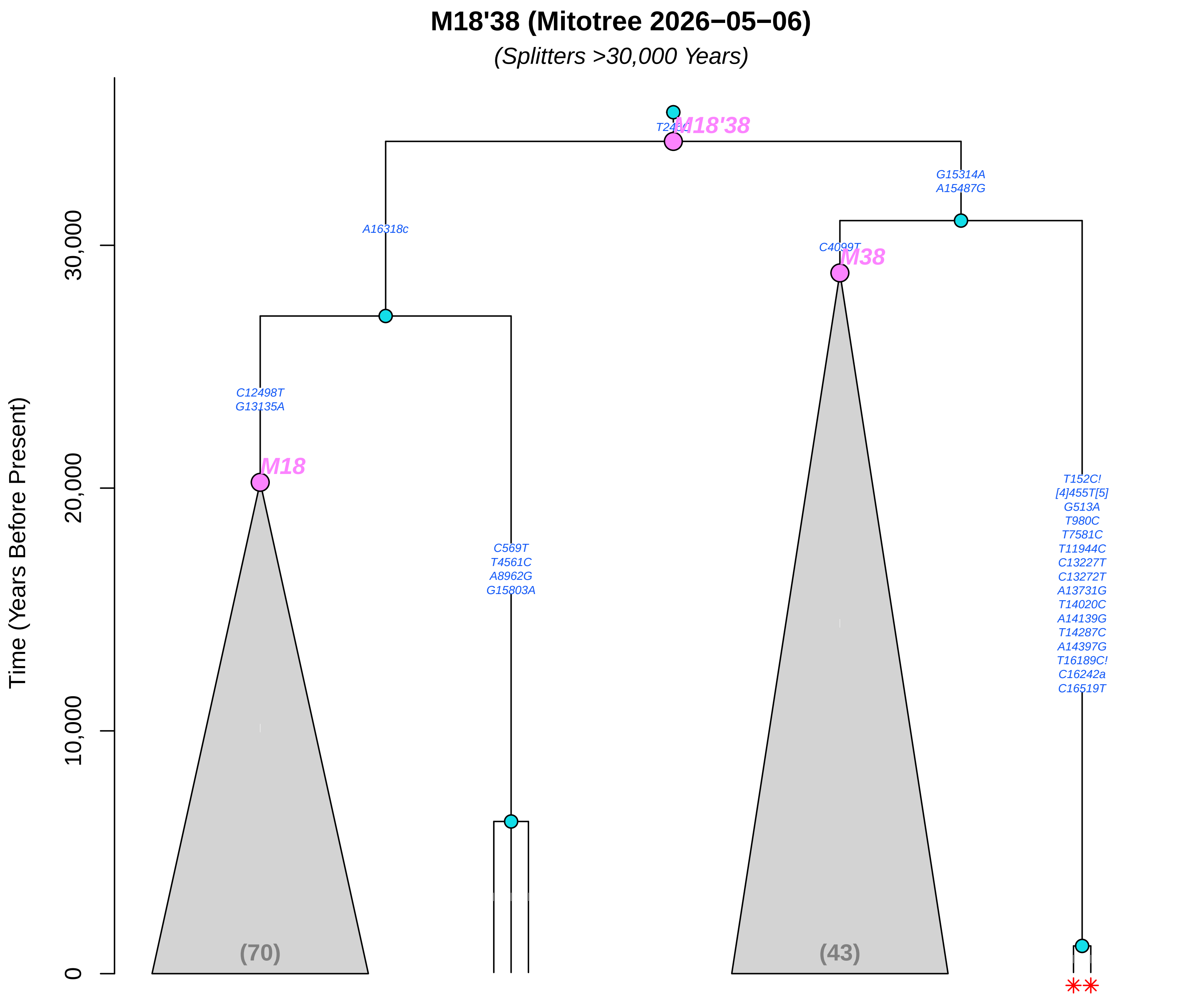

M18'38 (Mitotree 2026−05−06)
(Splitters >30,000 Years)
Time (Years Before Present)
30,000
20,000
10,000
0
M18'38
G15314A
A15487G
A16318c
C4099T
M38
C12498T
G13135A
M18
C569T
T4561C
A8962G
G15803A
T152C!
[4]455T[5]
G513A
T980C
T7581C
T11944C
C13227T
C13272T
A13731G
T14020C
A14139G
T14287C
A14397G
T16189C!
C16242a
C16519T
(70)
(43)

### M2 (Mitotree 2026−05−06)

*(Splitters >30,000 Years)*

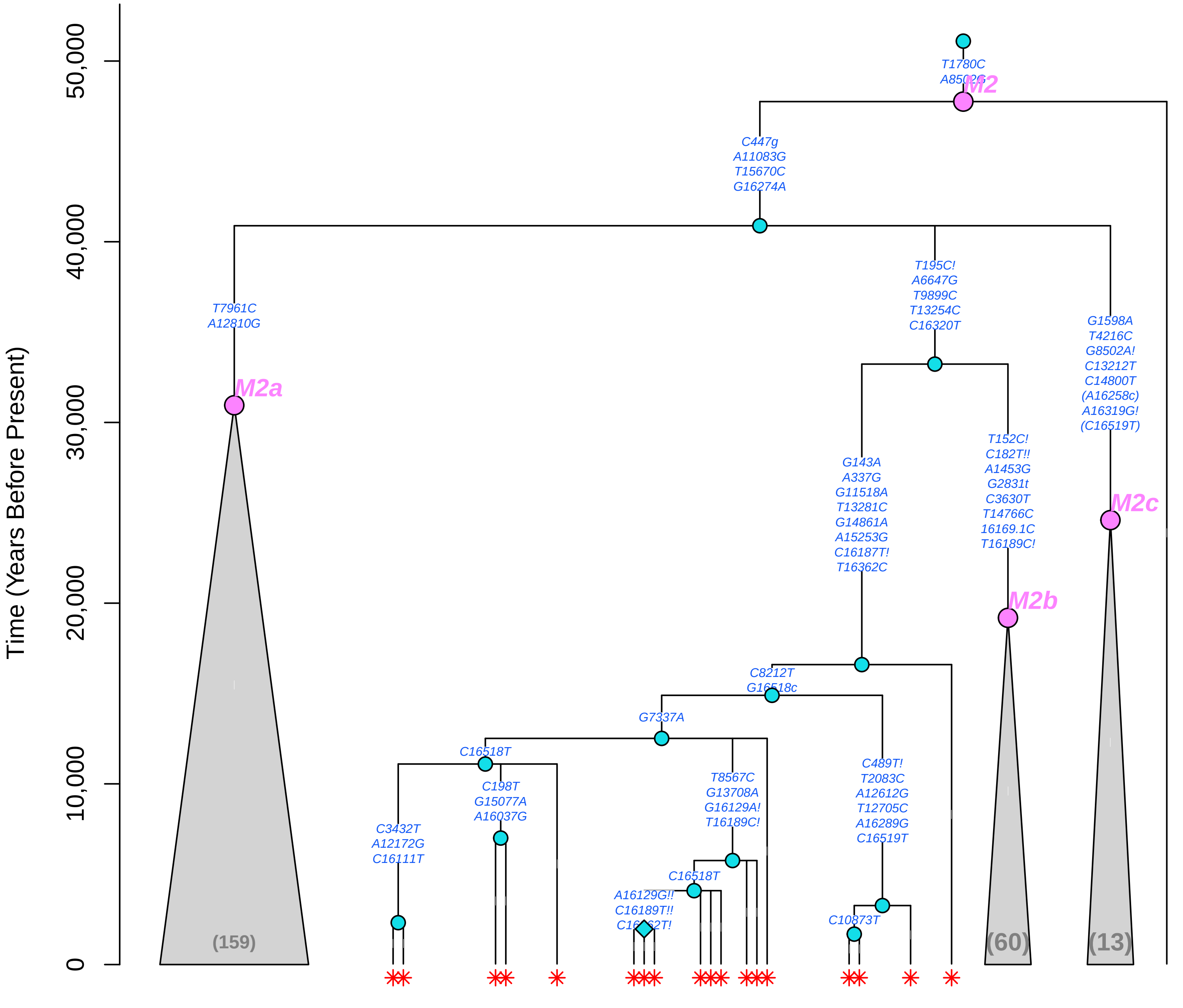

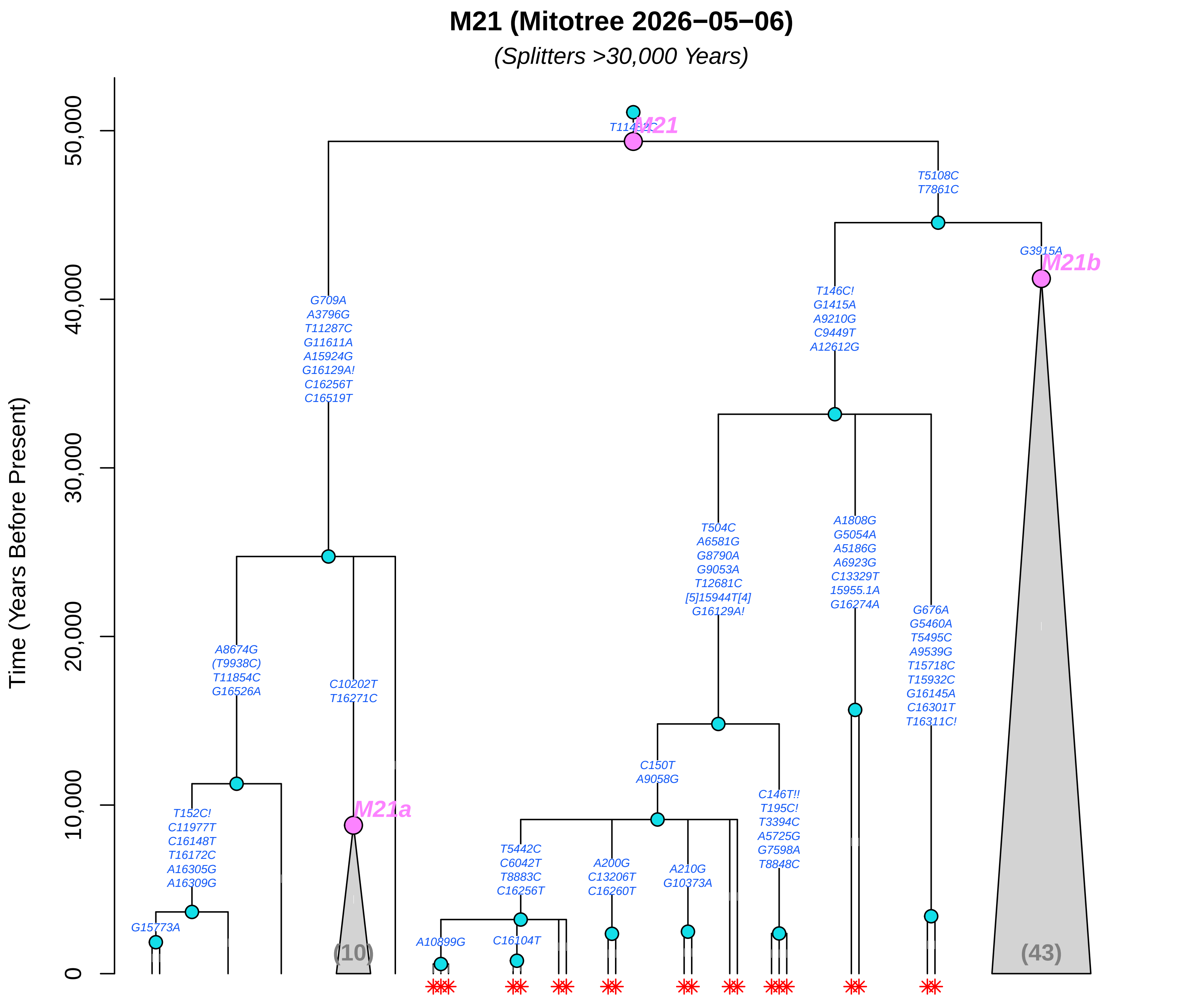

M21 (Mitotree 2026−05−06)
(Splitters >30,000 Years)
Time (Years Before Present)
50,000
40,000
30,000
20,000
10,000
0
M21
M21a
M21b
T5108C
T7861C
G3915A
G709A
A3796G
T11287C
G11611A
A15924G
G16129A!
C16256T
C16519T
T146C!
G1415A
A9210G
C9449T
A12612G
T504C
A6581G
G8790A
G9053A
T12681C
[5]15944T[4]
G16129A!
A1808G
G5054A
A5186G
A6923G
C13329T
15955.1A
G16274A
G676A
G5460A
T5495C
A9539G
T15718C
T15932C
G16145A
C16301T
T16311C!
A8674G
(T9938C)
T11854C
G16526A
C10202T
T16271C
C150T
A9058G
C146T!!
T195C!
T3394C
A5725G
G7598A
T8848C
T152C!
C11977T
C16148T
T16172C
A16305G
A16309G
G15773A
T5442C
C6042T
T8883C
C16256T
A200G
C13206T
C16260T
A210G
G10373A
A10899G
C16104T
(10)
(43)

#### M23'75 (Mitotree 2026−05−06)

*(Splitters >30,000 Years)*

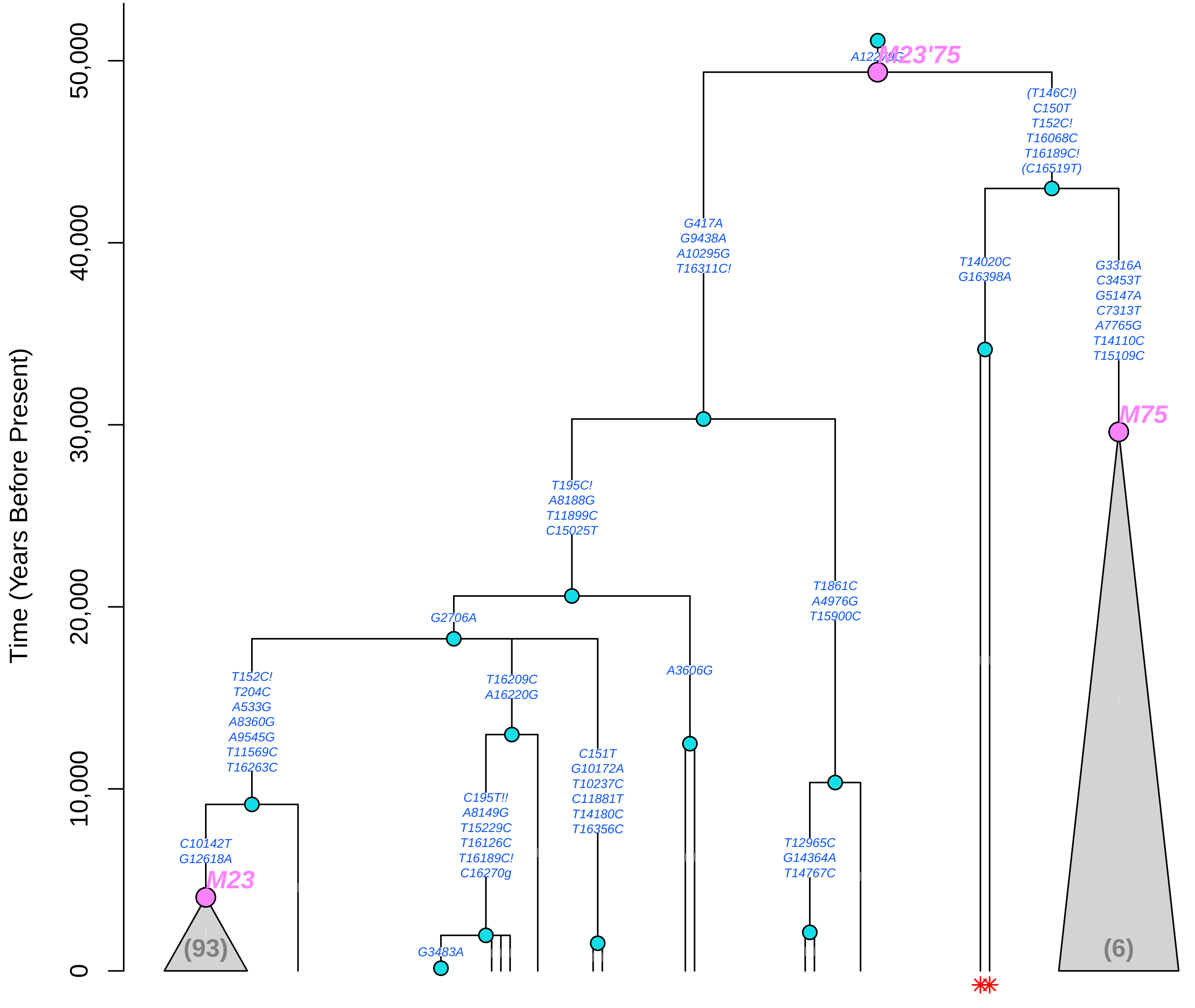

#### M33 (Mitotree 2026−05−06)

*(Splitters >30,000 Years)*

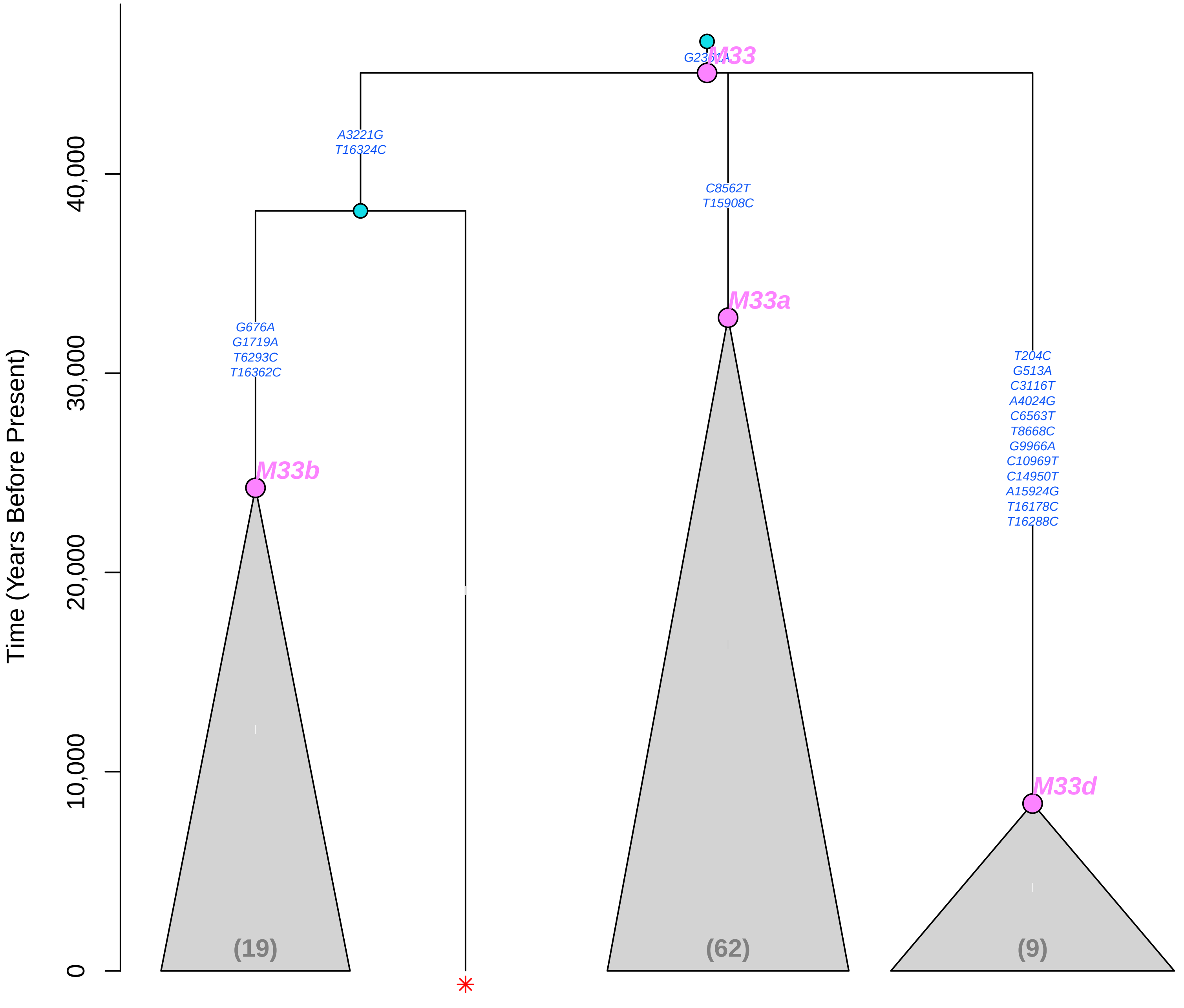

#### M34'57 (Mitotree 2026−05−06)

*(Splitters >30,000 Years)*

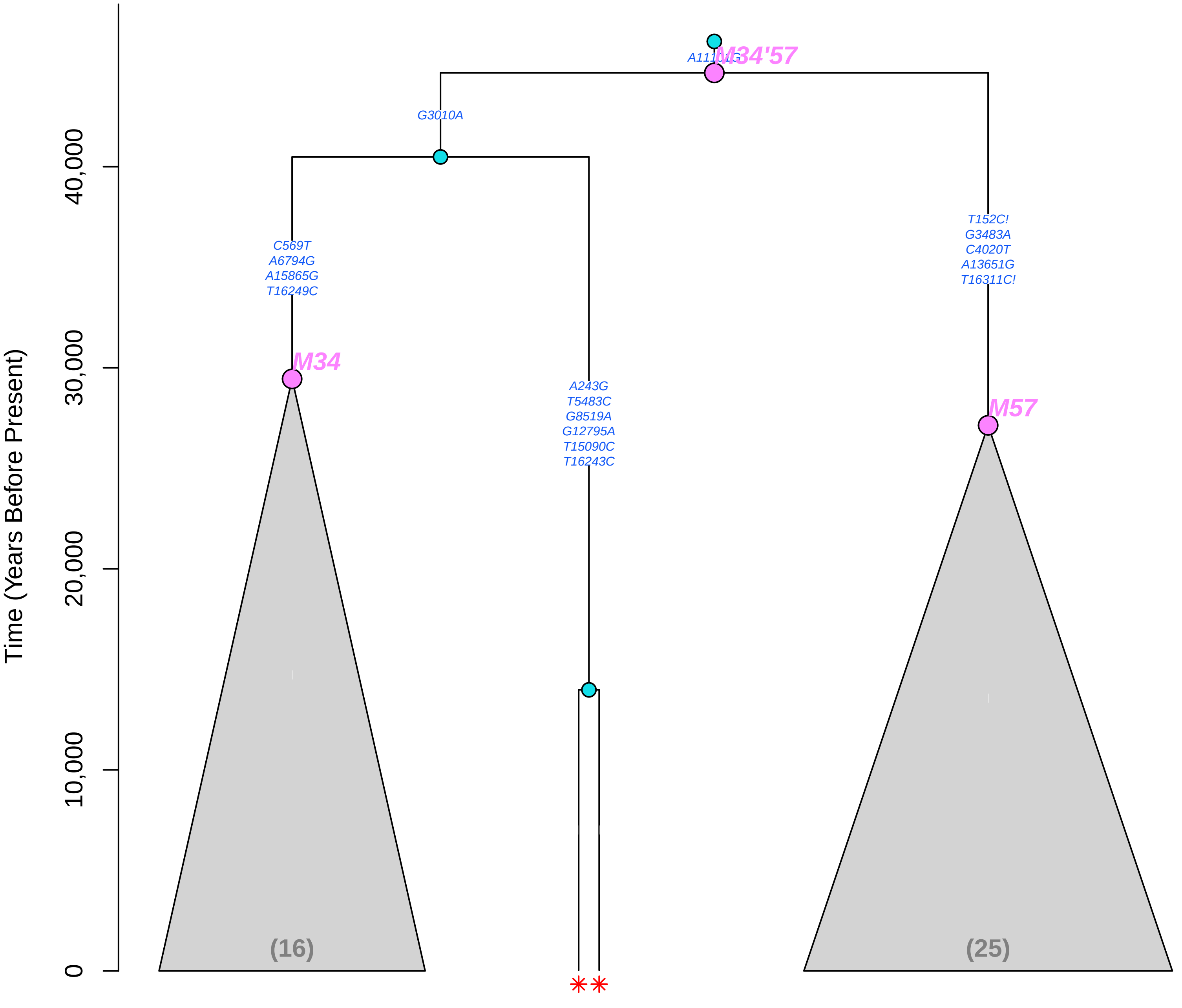

### M4"67 (Mitotree 2026−05−06)

*(Splitters >30,000 Years)*

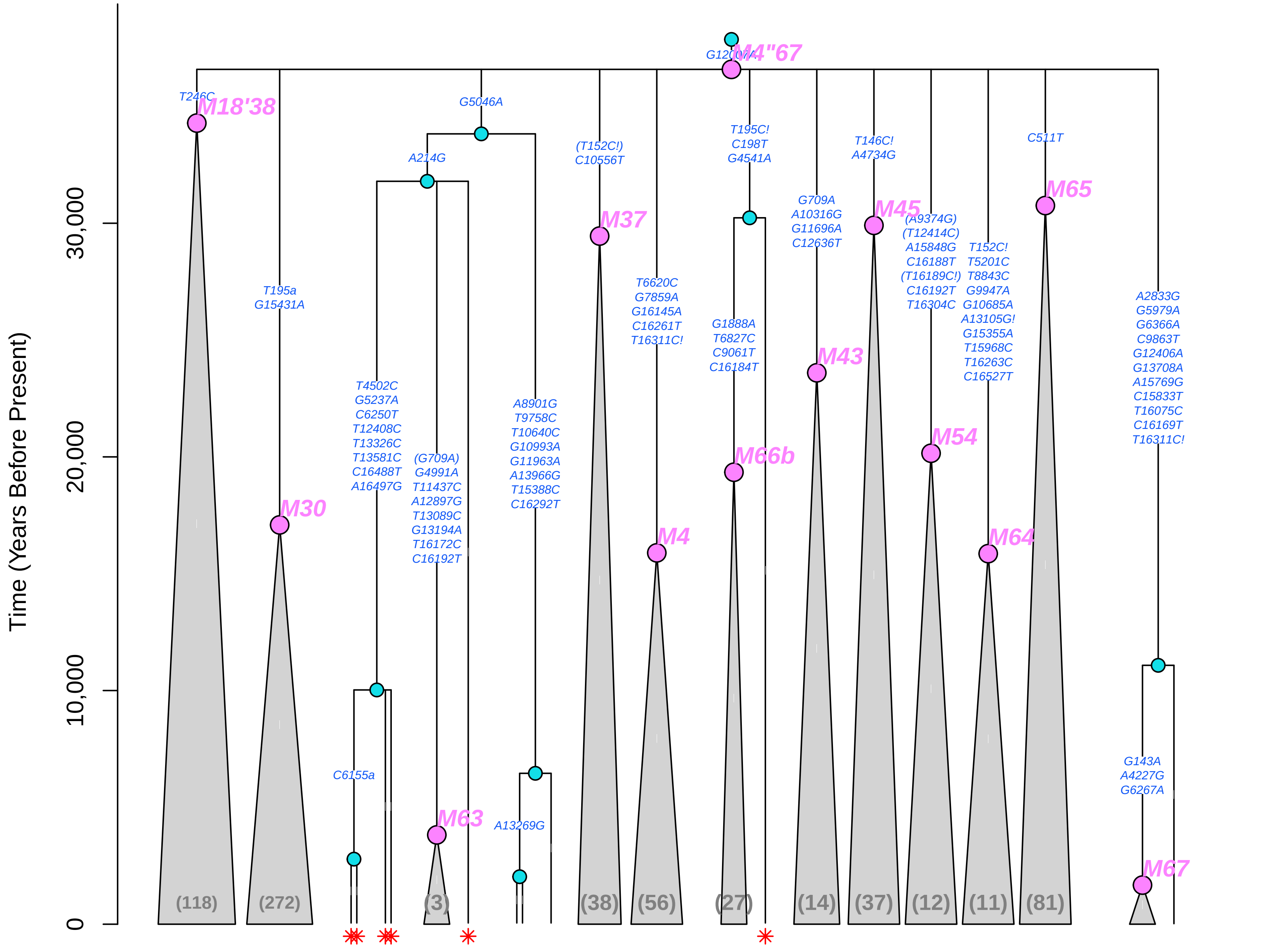

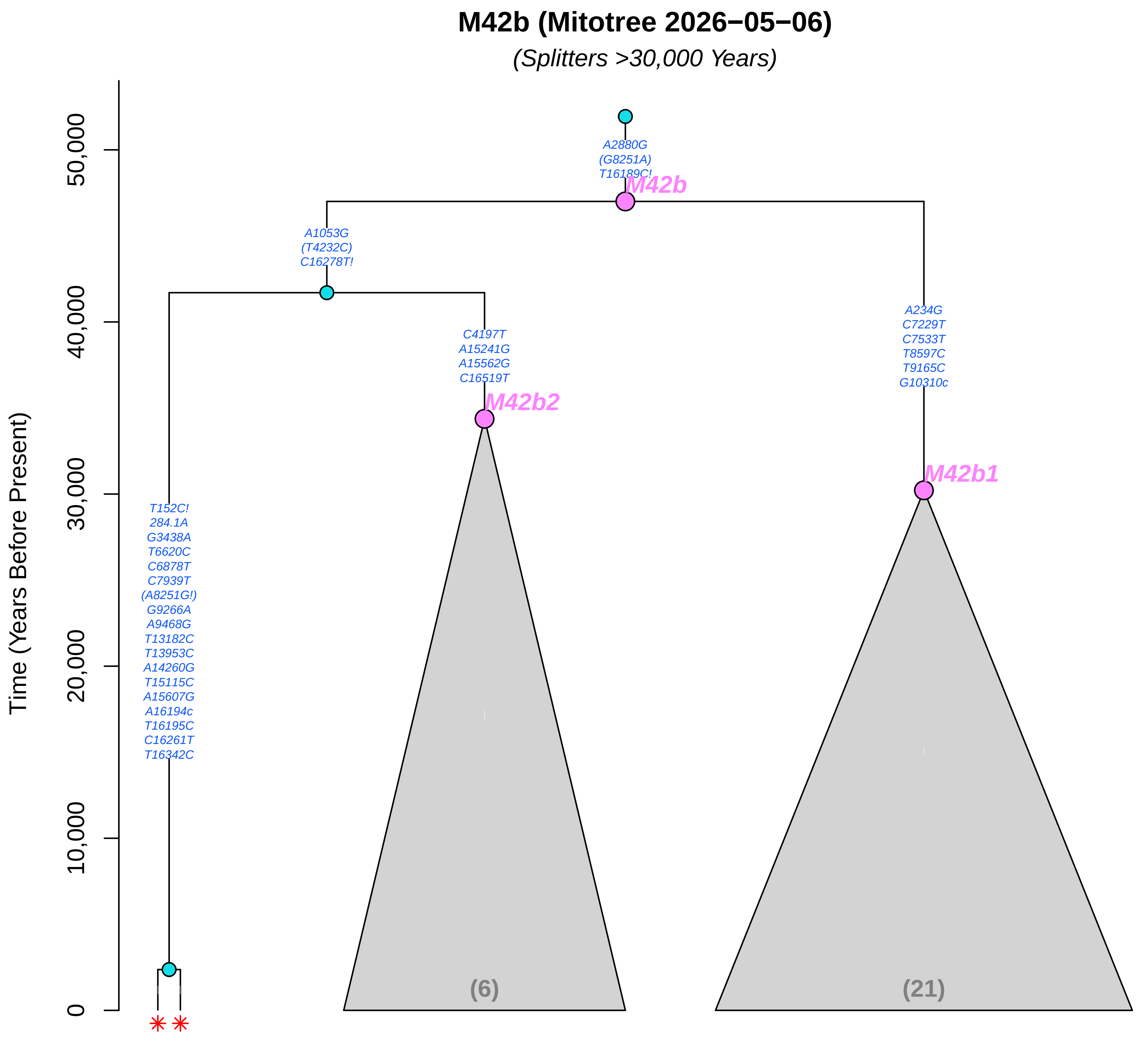

M42b (Mitotree 2026−05−06)
(Splitters >30,000 Years)
Time (Years Before Present)
0
10,000
20,000
30,000
40,000
50,000
A2880G
(G8251A)
T16189C!
M42b
A1053G
(T4232C)
C16278T!
C4197T
A15241G
A15562G
C16519T
M42b2
A234G
C7229T
C7533T
T8597C
T9165C
G10310c
M42b1
T152C!
284.1A
G3438A
T6620C
C6878T
C7939T
(A8251G!)
G9266A
A9468G
T13182C
T13953C
A14260G
T15115C
A15607G
A16194c
T16195C
C16261T
T16342C
(6)
(21)

**M5 (Mitotree 2026−05−06)**

*(Splitters >30,000 Years)*

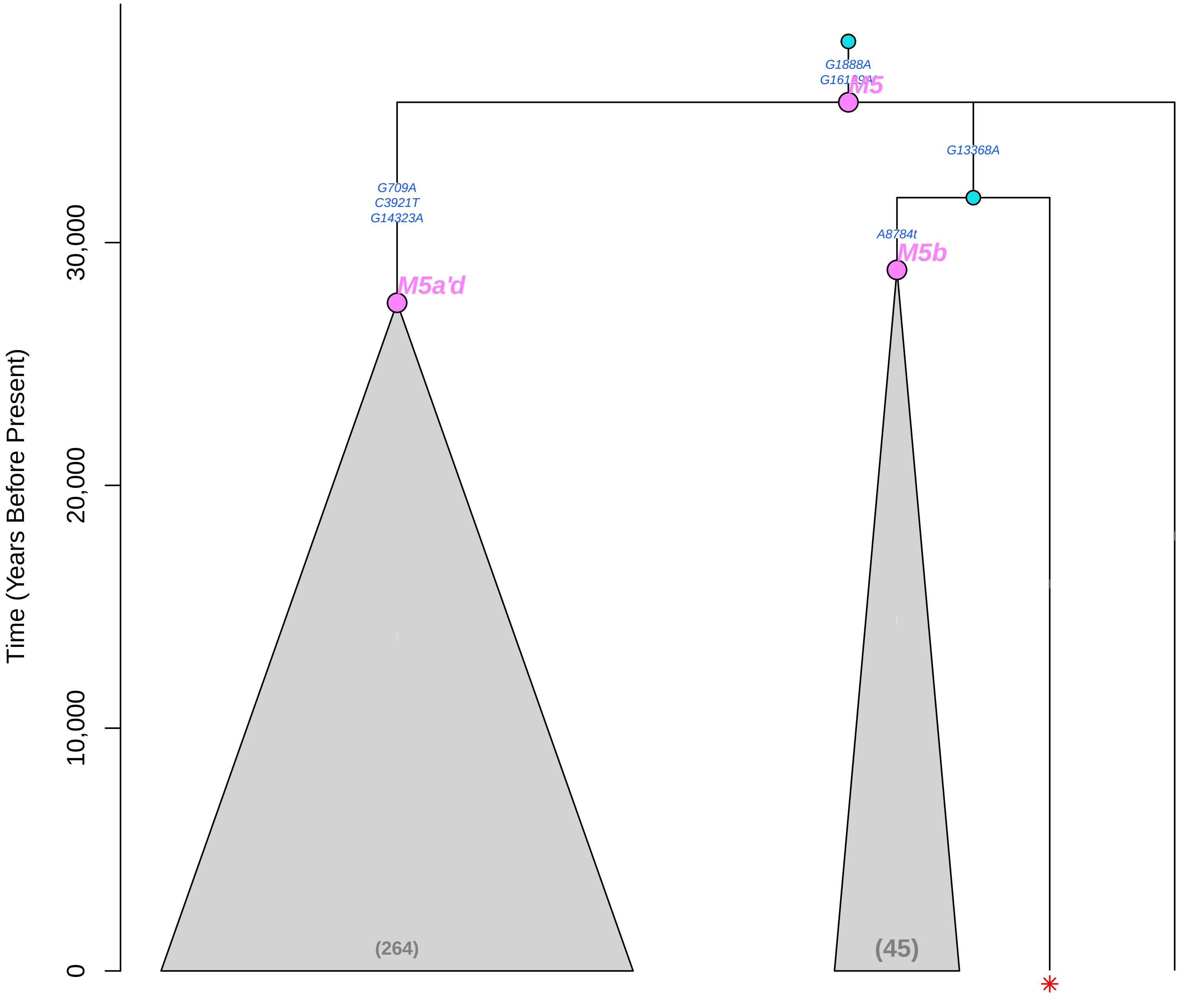

#### M52b (Mitotree 2026−05−06)

*(Splitters >30,000 Years)*

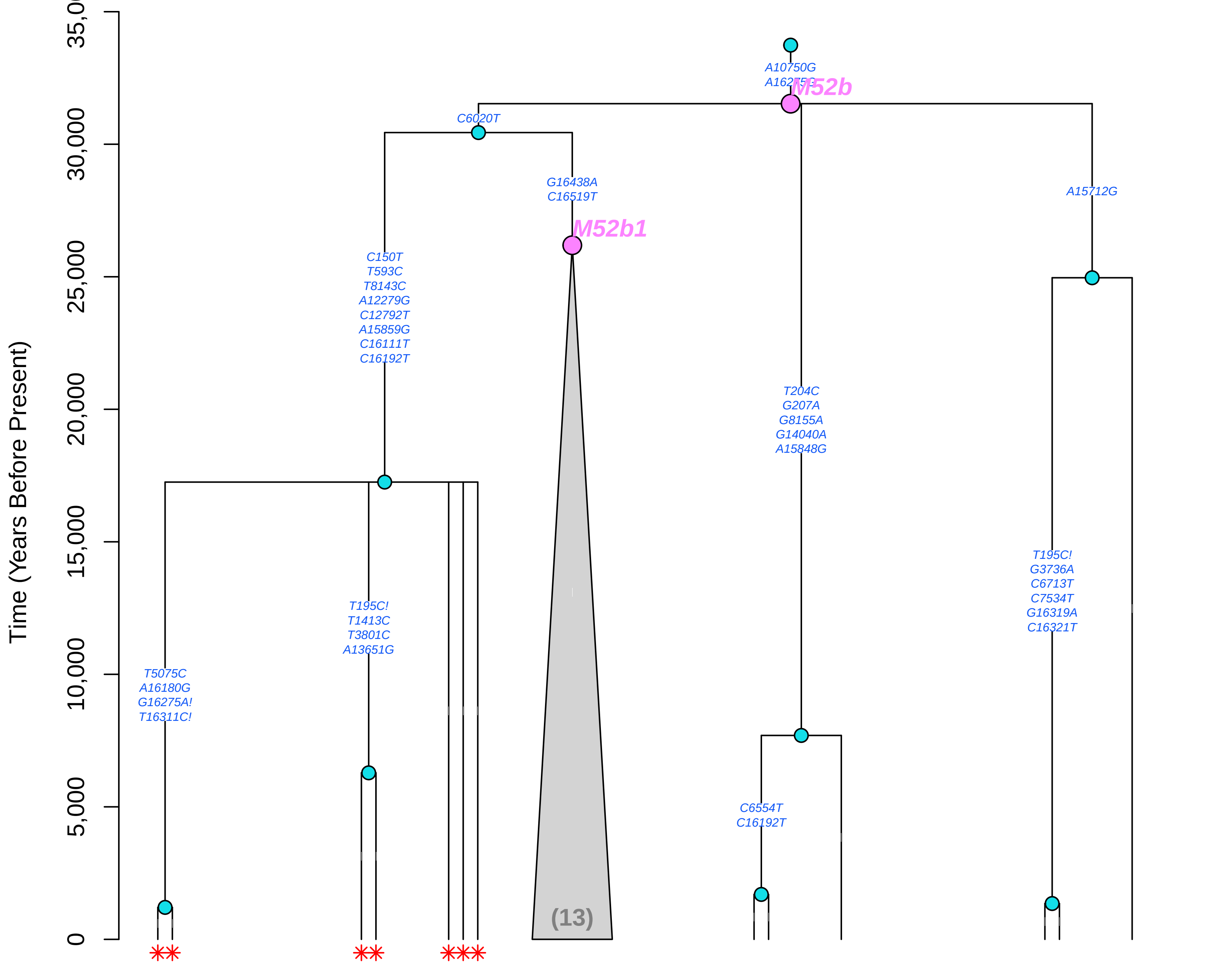

#### M60 (Mitotree 2026−05−06)

*(Splitters >30,000 Years)*

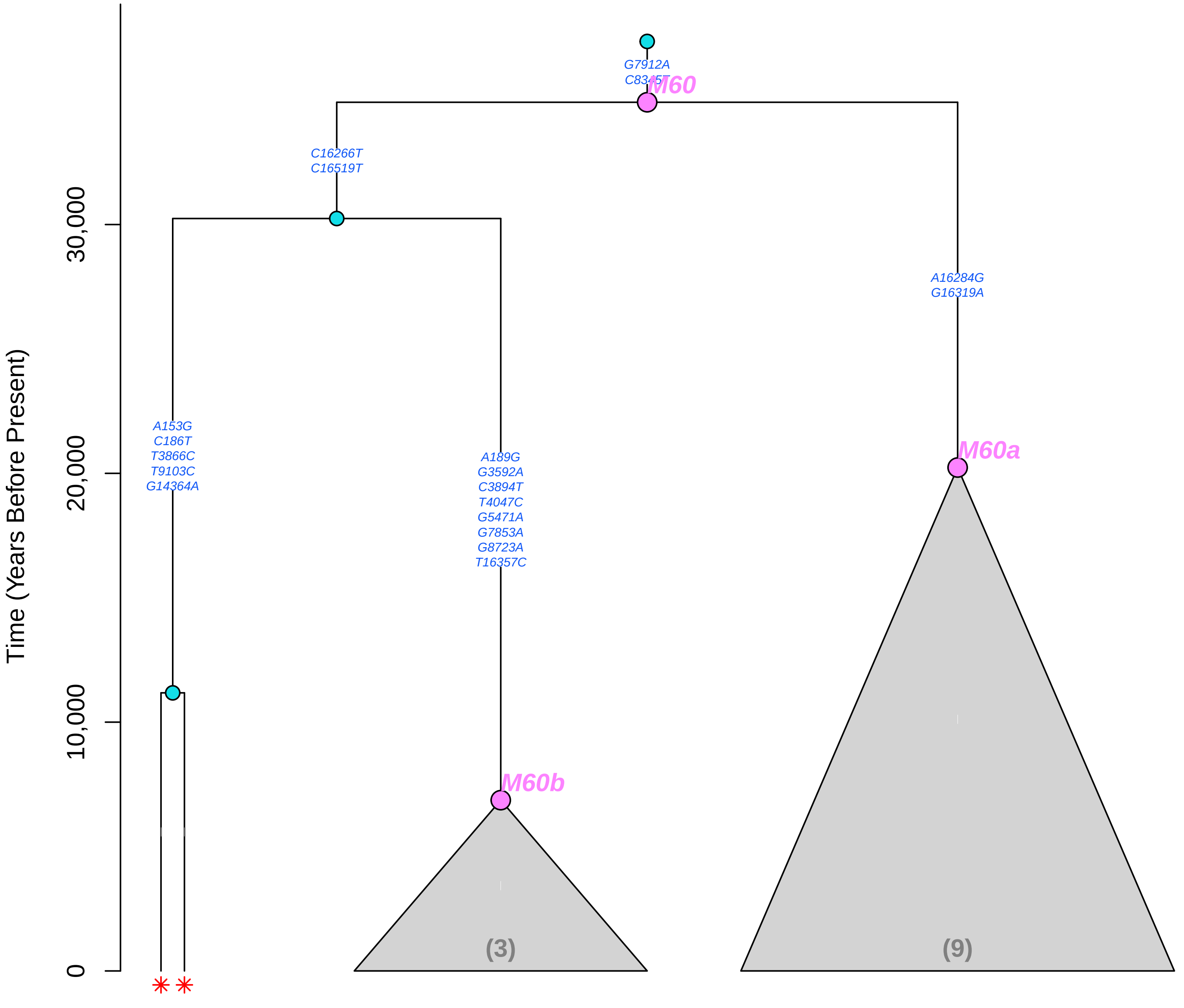

#### M7b (Mitotree 2026−05−06)

*(Splitters >30,000 Years)*

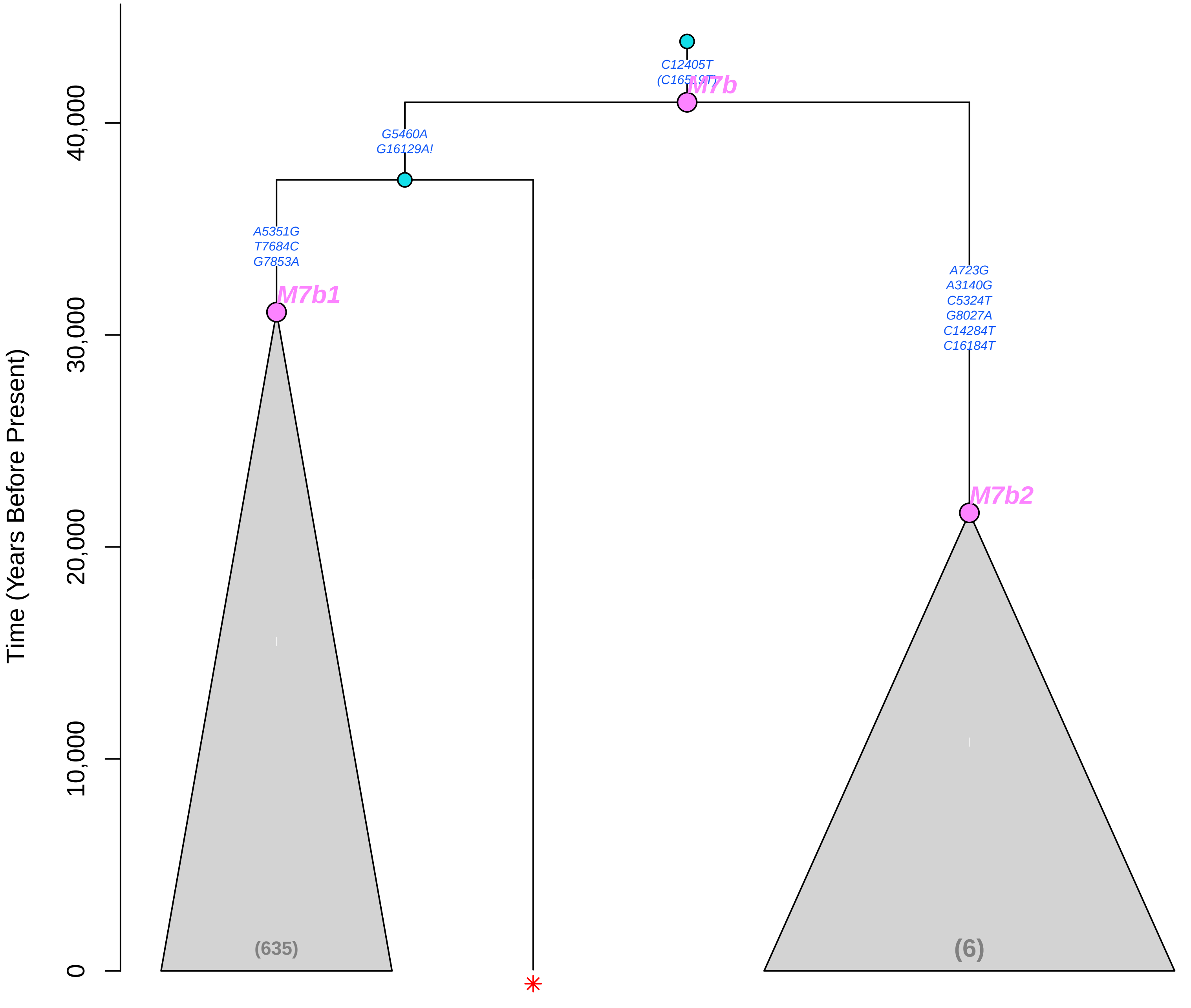

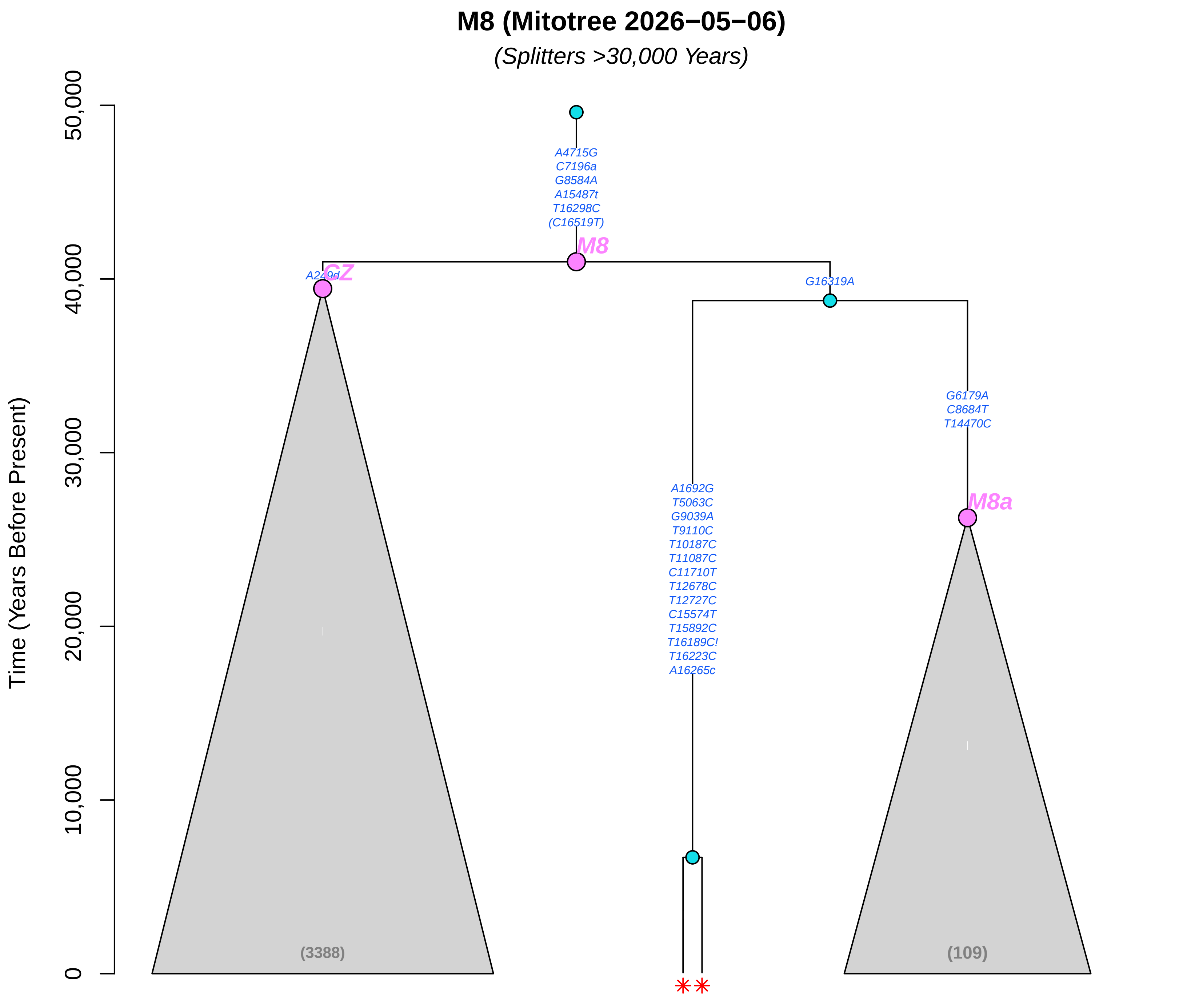
M8 (Mitotree 2026−05−06)
(Splitters >30,000 Years)
Time (Years Before Present)
50,000
40,000
30,000
20,000
10,000
0
A4715G
C7196a
G8584A
A15487t
T16298C
(C16519T)
M8
A249d
CZ
G16319A
G6179A
C8684T
T14470C
M8a
A1692G
T5063C
G9039A
T9110C
T10187C
T11087C
C11710T
T12678C
T12727C
C15574T
T15892C
T16189C!
T16223C
A16265c
(3388)
(109)

### N (Mitotree 2026−05−06)

*(Splitters >30,000 Years)*

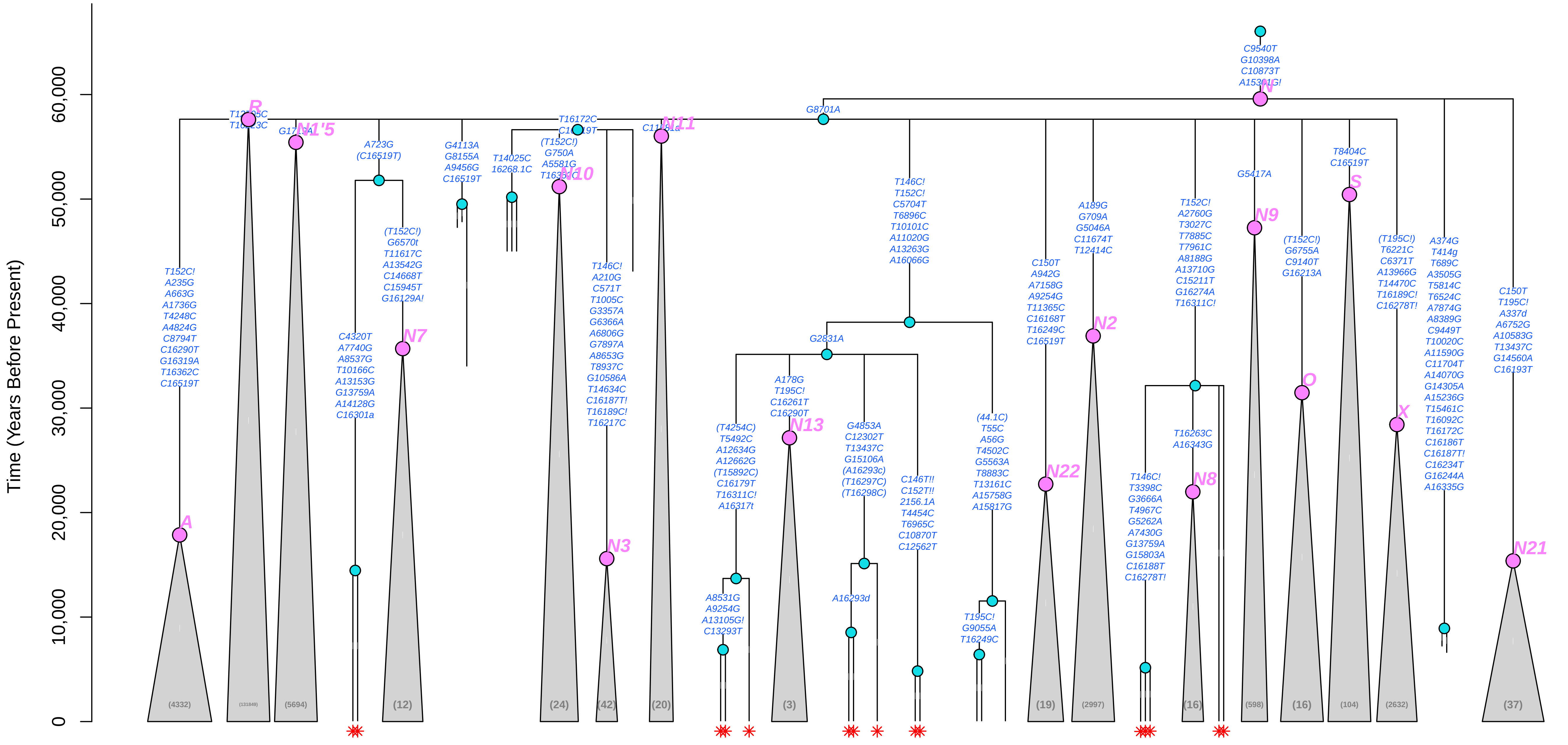

**N10 (Mitotree 2026−05−06)**

*(Splitters >30,000 Years)*

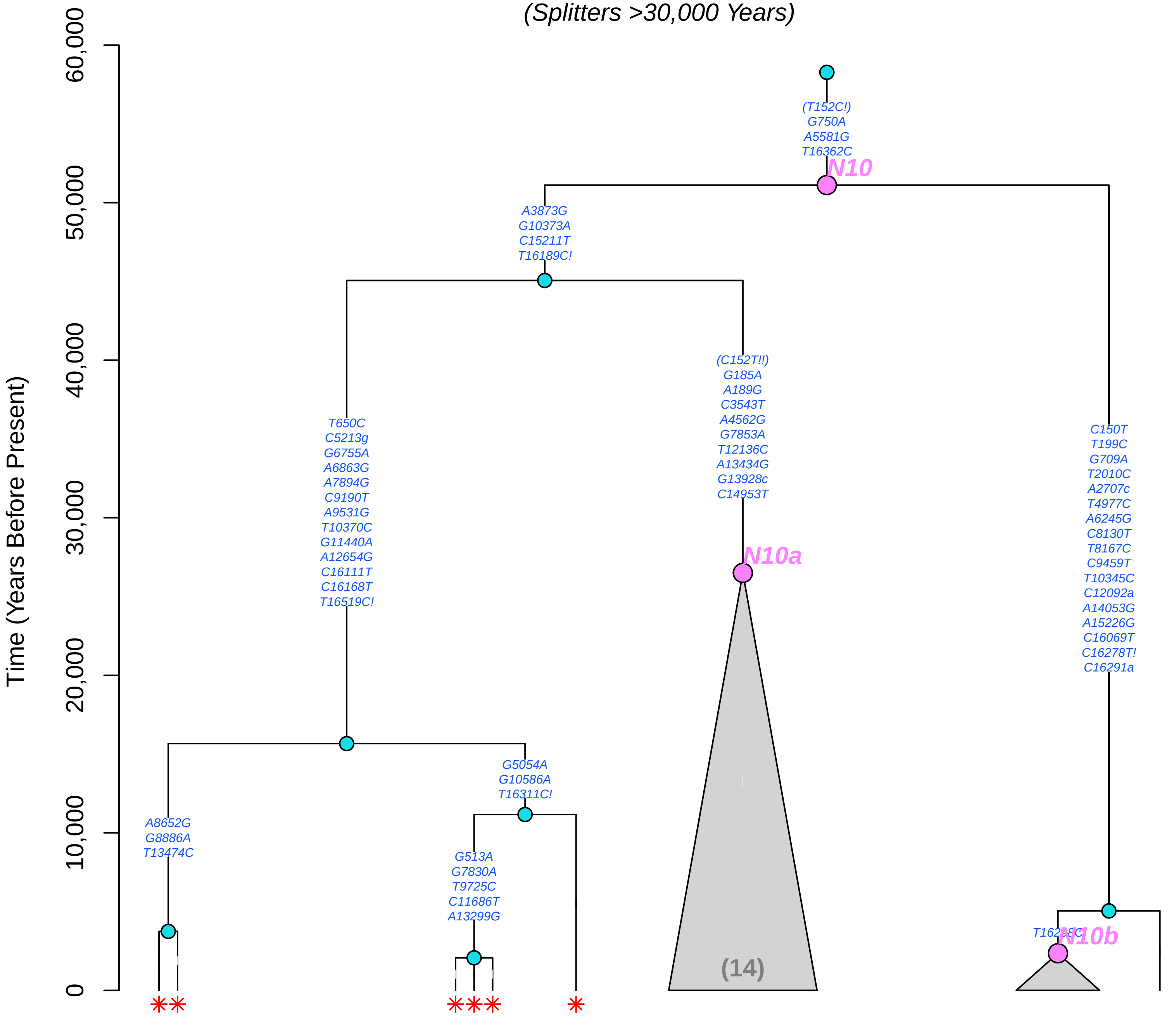

### P (Mitotree 2026−05−06)

*(Splitters >30,000 Years)*

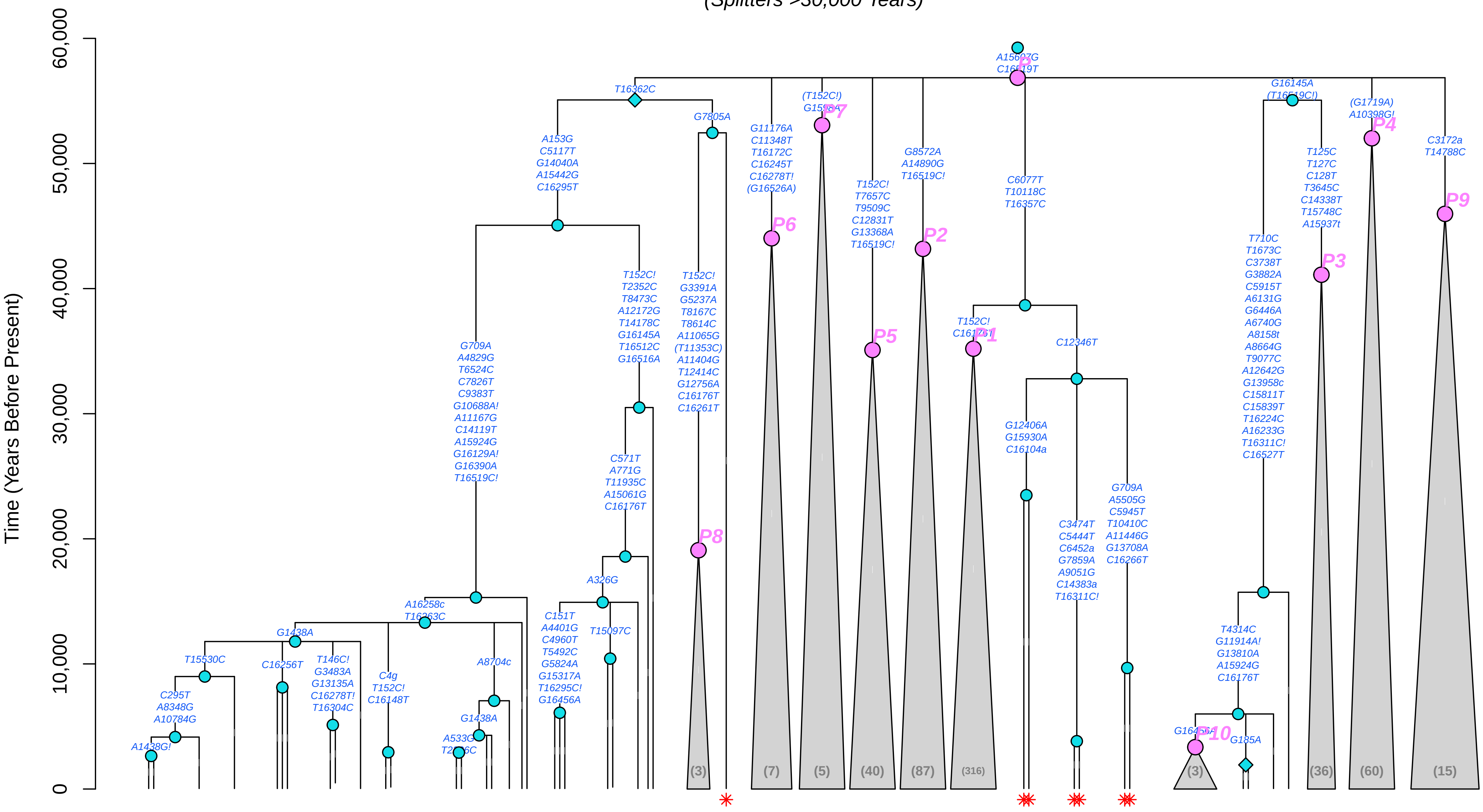

#### P3b (Mitotree 2026−05−06)

*(Splitters >30,000 Years)*

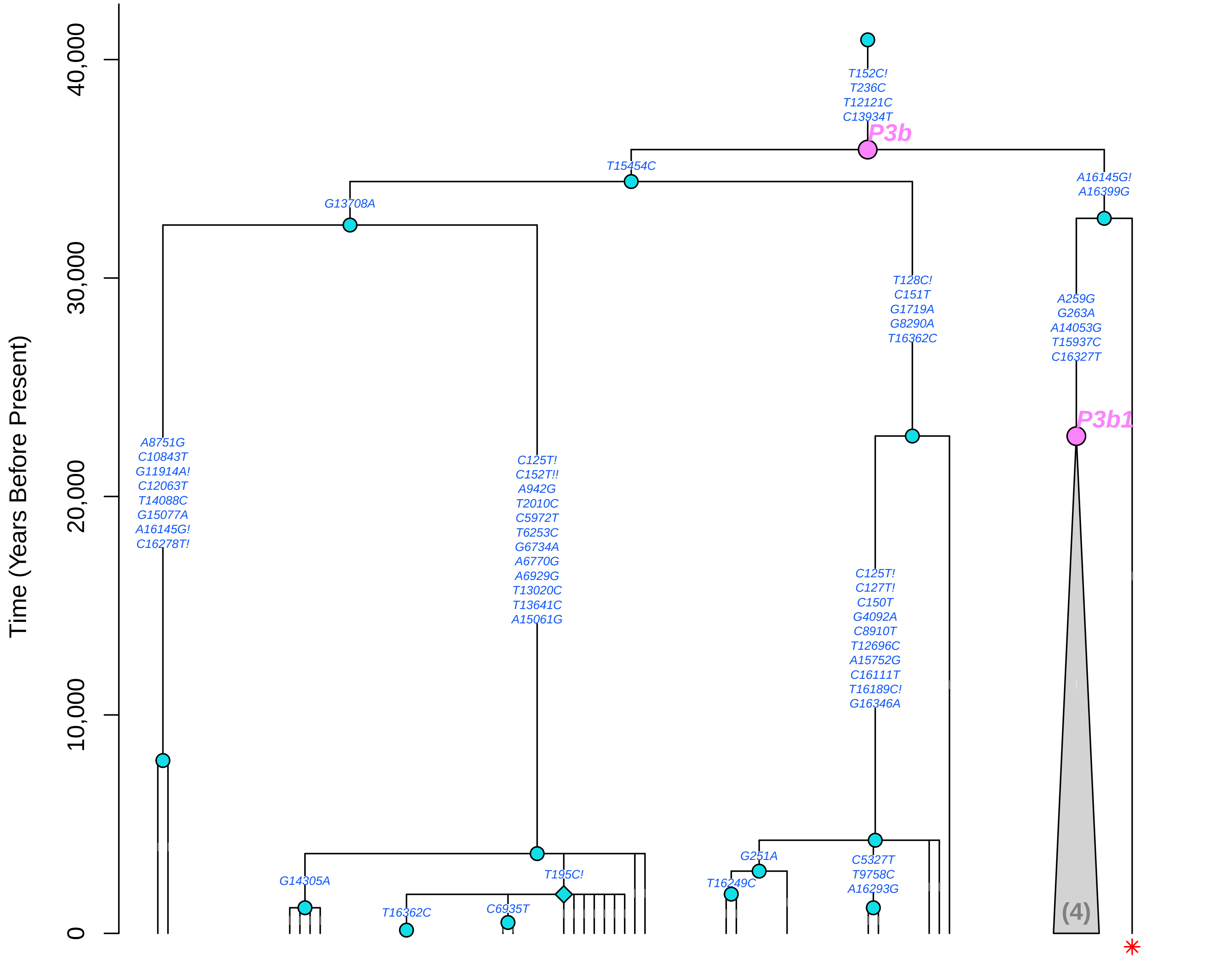

### R (Mitotree 2026−05−06)

*(Splitters >30,000 Years)*

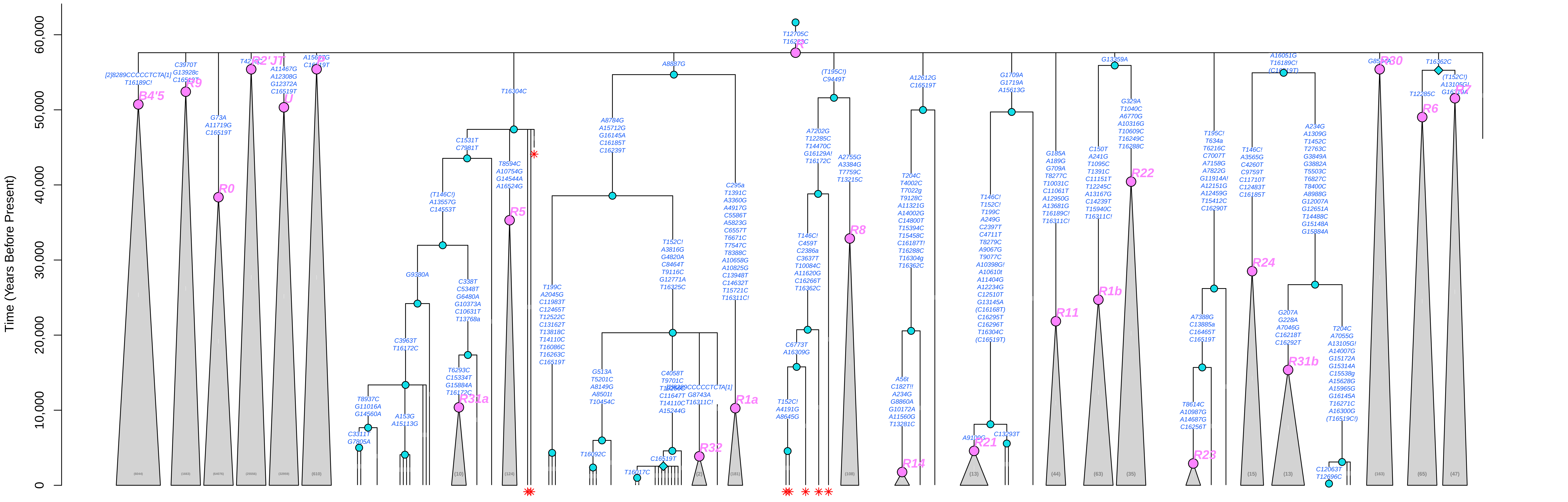

### R30a (Mitotree 2026−05−06)

*(Splitters >30,000 Years)*

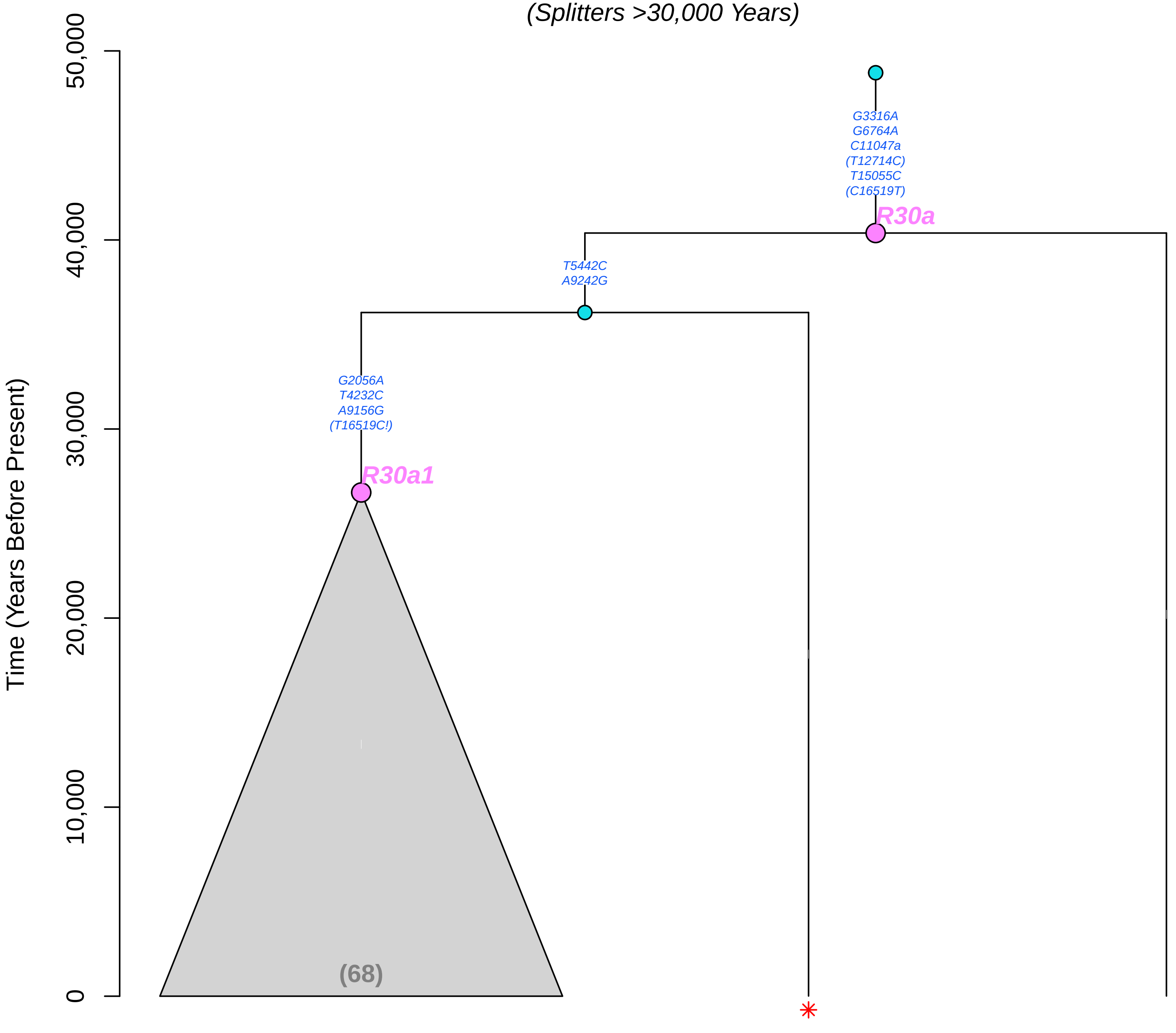

### R7 (Mitotree 2026−05−06)

*(Splitters >30,000 Years)*

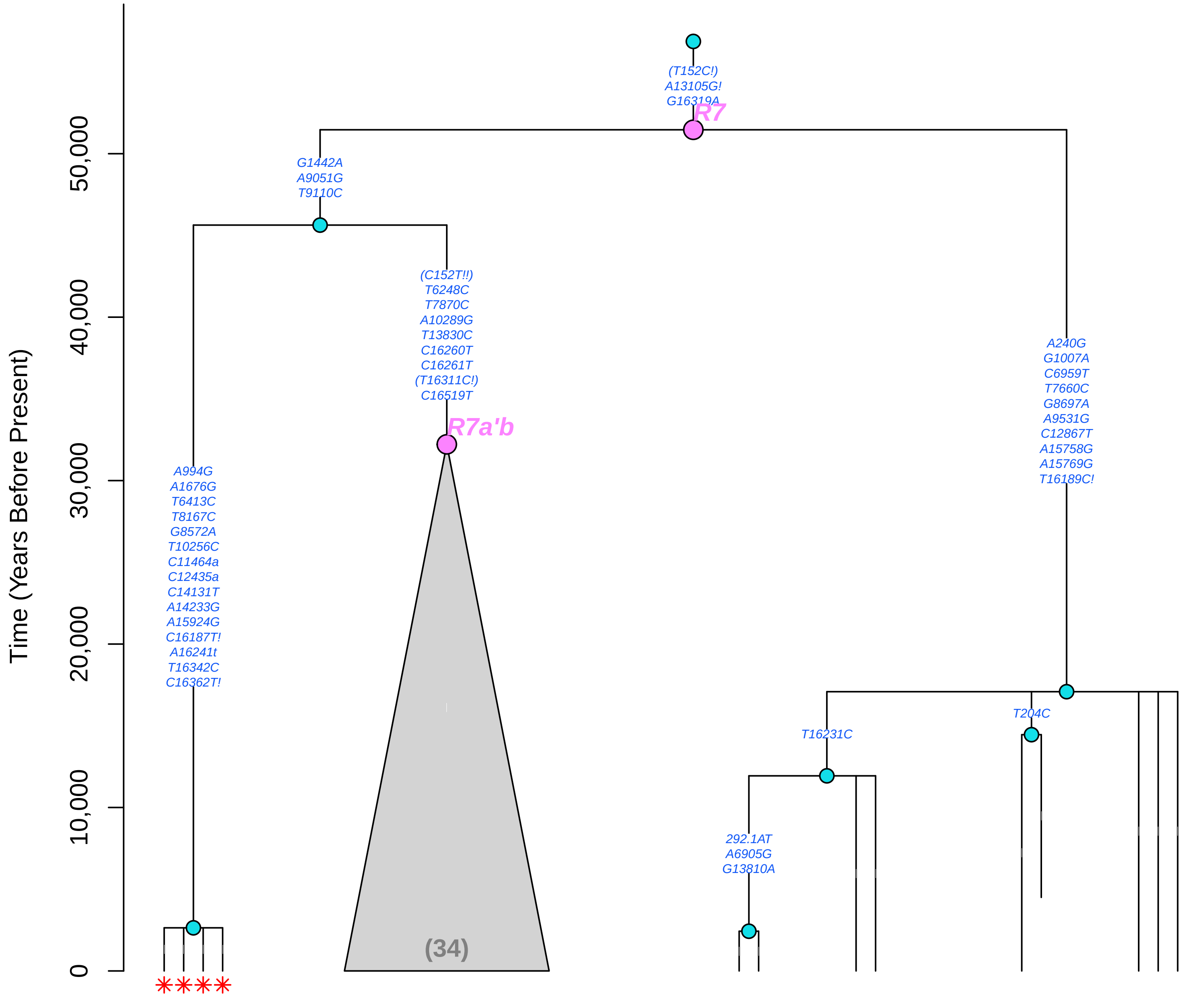

### S (Mitotree 2026−05−06)

*(Splitters >30,000 Years)*

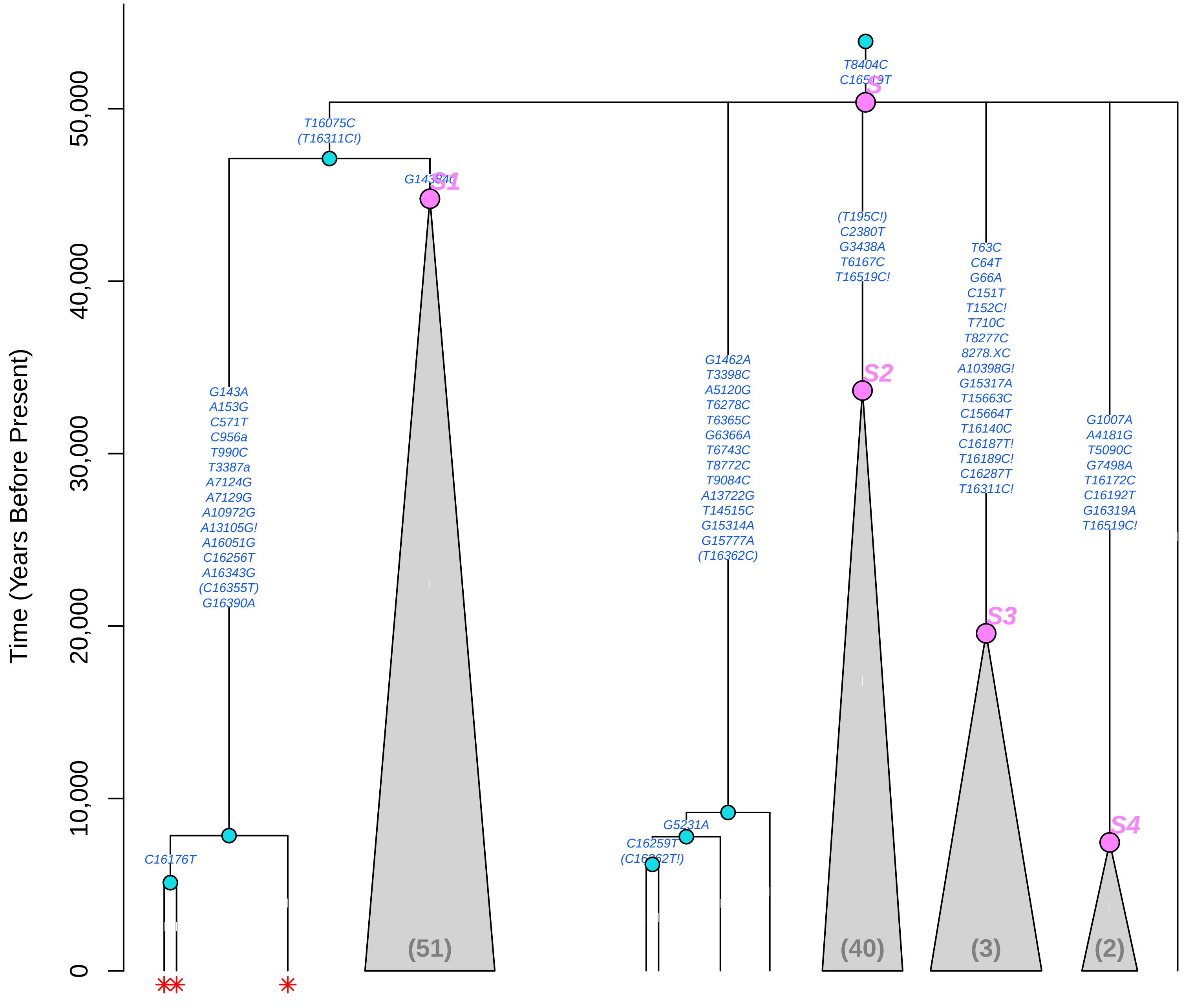

### U2 (Mitotree 2026−05−06)

*(Splitters >30,000 Years)*

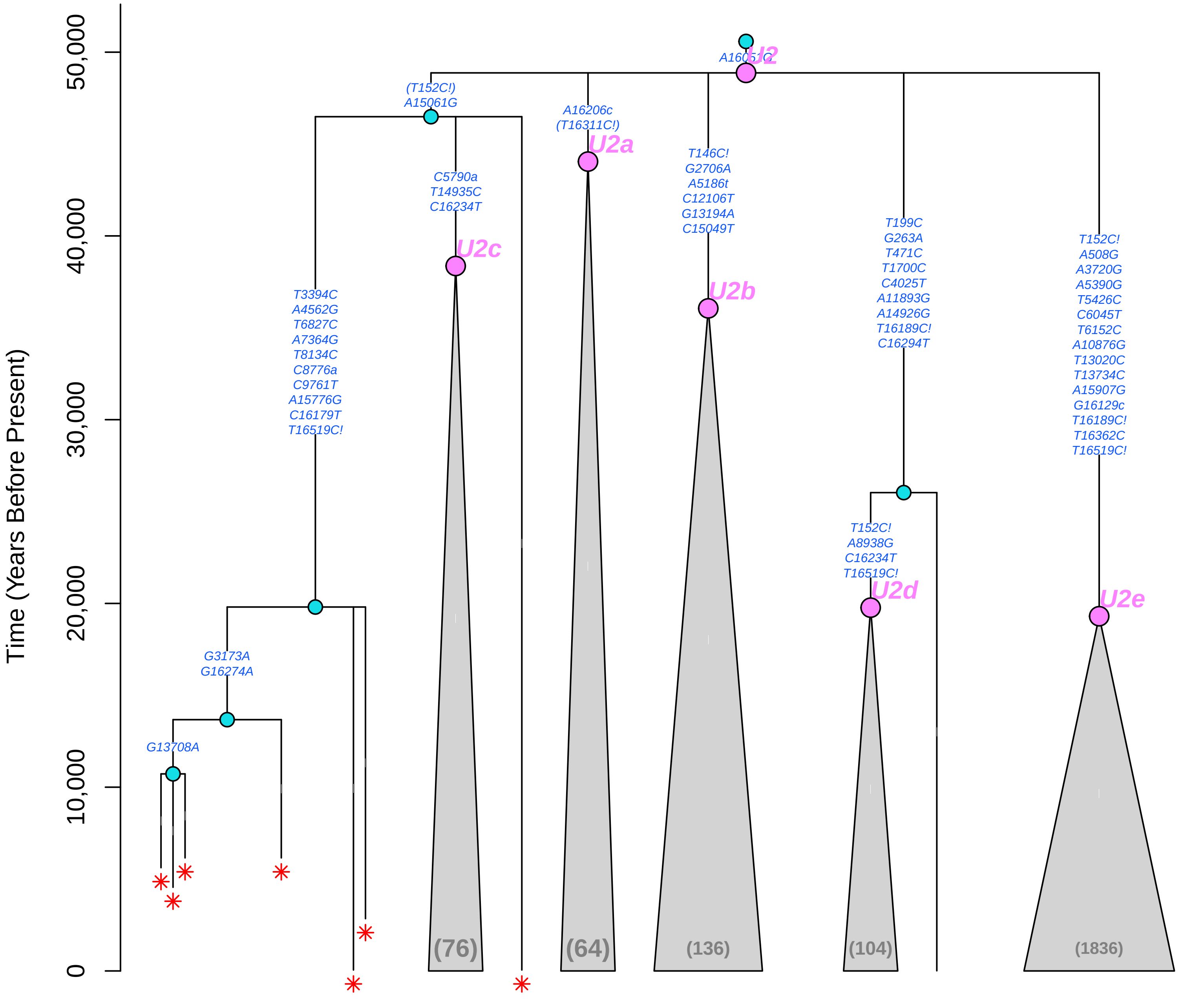

#### U2'3'4'7'8'9 (Mitotree 2026−05−06)

*(Splitters >30,000 Years)*

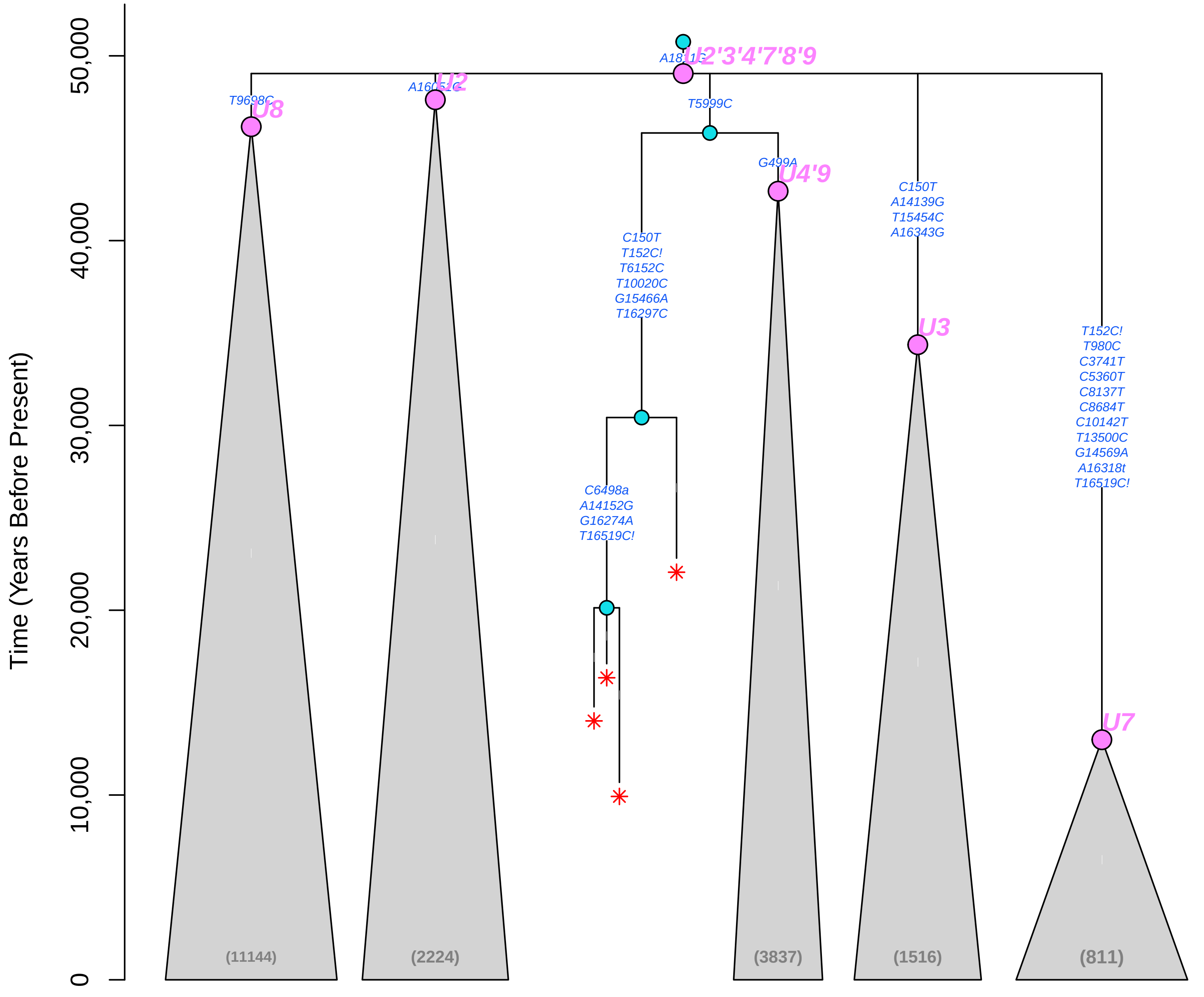

### U4'9 (Mitotree 2026−05−06)

*(Splitters >30,000 Years)*

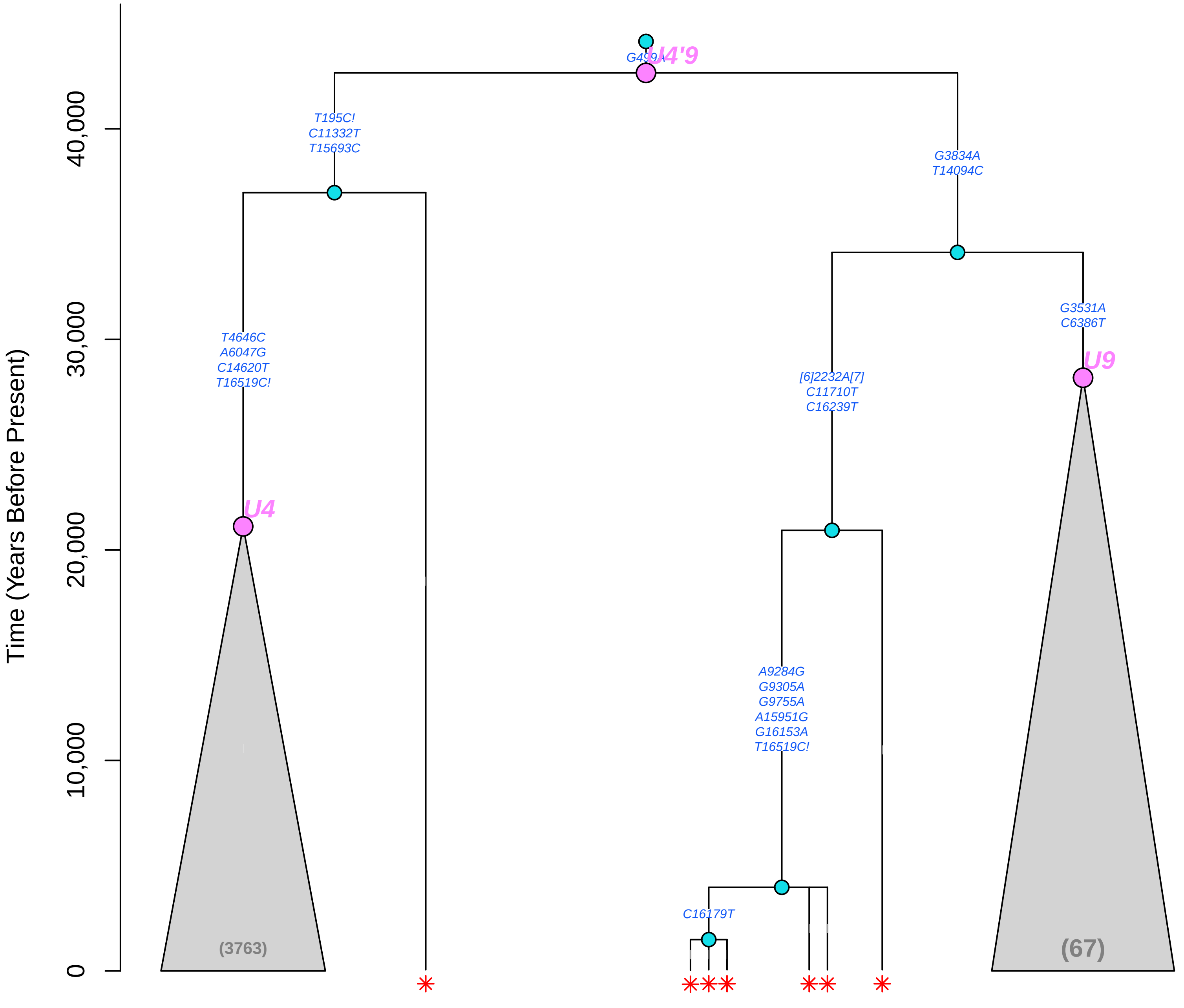
